# Priming versus propagating: distinct immune effects of an alpha- versus beta-particle emitting radiopharmaceutical when combined with immune checkpoint inhibition

**DOI:** 10.1101/2024.12.26.630430

**Authors:** Caroline P. Kerr, Won Jong Jin, Peng Liu, Joseph J. Grudzinski, Carolina A. Ferreira, Hansel Comas Rojas, Alejandro J. Oñate, Ohyun Kwon, Meredith Hyun, Malick Bio Idrissou, René Welch Schwartz, Jessica M. Vera, Paul A. Clark, Maya Takashima, Amy K. Erbe, Amanda G. Shea, Maria Powers, Anatoly N. Pinchuk, Christopher F. Massey, Cynthia Choi, Reinier Hernandez, Bryan P. Bednarz, Irene M. Ong, Jamey P. Weichert, Zachary S. Morris

## Abstract

Radiopharmaceutical therapy (RPT) enhances tumor response to immune checkpoint inhibitors (ICI) in preclinical models, but the effects of different radioisotopes have not been thoroughly compared. To evaluate mechanisms of response to RPT+ICI, we used NM600, an alkylphosphocholine selectively taken up by most tumors. Effects of ^90^Y-, ^177^Lu-, and ^225^Ac-NM600 + ICIs were compared in syngeneic murine models, B78 melanoma (poorly immunogenic) and MC38 colorectal cancer (immunogenic). ^90^Y-/^177^Lu-/or ^225^Ac-NM600 delivering 2 Gy mean tumor dose promoted tumor regression and improved survival when combined with ICIs in syngeneic mice bearing B78 or MC38 tumors. Regardless of the administered isotope, this combination was optimized with early ICI administration (days -3/0/3) relative to day 1 RPT. ^90^Y-NM600+ICI produced the greatest anti-tumor response for MC38, whereas high linear energy transfer (LET) alpha particle radiation from ^225^Ac-NM600+ICI was most effective against poorly immunogenic B78 tumors. Flow cytometry and single cell RNA and T cell receptor (TCR) sequencing illuminated distinct mechanisms of ^90^Y- or ^177^Lu-NM600 in promoting expansion of existing adaptive immunity and of ^225^Ac-NM600 in promoting immune priming when combined with ICI. Antitumor immune response can be achieved with appropriate application of α- or β- emitting RPT in combination with ICIs in diverse murine tumor models.

## INTRODUCTION

Combinations of radiation therapy (RT) and immune checkpoint inhibitors (ICI) have shown efficacy preclinically and in certain clinical scenarios^1–5^. Preclinical mechanisms by which RT enhances response to immunotherapy include inducing immunogenic cell death, releasing tumor specific antigens and damage-associated molecular pattern molecules, and causing phenotypic changes on surviving tumor cells (upregulation of MHC-I and FAS, type I interferon secretion)^6–11^. However, adding RT to a single tumor site in patients with metastatic disease has not improved the rate or duration of systemic response to standard-of-care ICIs^12^. Several mechanisms have been proposed to account for this translational discrepancy, including patient selection (e.g., patients with immune exhausted tumor microenvironments have worse outcomes^2^), radiation of tumor draining lymph nodes necessary for propagation of an adaptive immune response^13^, and non-optimal dosing or timing of radiation^14,15^.

With a growing understanding of how RT enhances responses to immunotherapy preclinically and clinically in locally advanced disease settings, it is important to consider how these therapies may be combined to improve outcomes in patients with metastatic disease. The role of external beam radiation therapy (EBRT) is limited in treatment of metastatic disease due to potentially prohibitive toxicities associated with irradiating multiple tumor sites or the whole body to cover radiographically occult micrometastatic disease. Additionally, unirradiated distant tumor sites have been shown to abrogate the *in situ* vaccine effect and resulting adaptive immune response following RT at one tumor site in certain preclinical studies^16,17^.

Radiopharmaceutical therapy (RPT) is a modality of RT whereby a radioactive isotope conjugated to a tumor targeting vector is administered systemically to deposit radiation dose at all sites of uptake. RPT agents have the potential to overcome some of the limitations of EBRT in metastatic disease settings. Physical properties of radiopharmaceuticals vary widely, with beta (β)- (e.g., ^90^Y, ^177^Lu) and alpha (α)-particle (e.g., ^223^Ra, ^225^Ac) emitting radioisotopes being the focus of much clinical interest presently. β particles are characterized by low linear energy transfer (LET) (0.2 keV/µm) and travel millimeters in tissue with sparse ionization events, whereas α particles are high LET particles (50-230 keV/µm) that travel micrometers in tissue with densely clustered ionization events along their track^18^. RPT agents have been studied both preclinically and in early clinical studies in combination with immunotherapy^19^. To capitalize on the therapeutic potential of combinations of RPT + ICIs, it is critical to understand the effects of RPT properties (e.g., radionuclide, vector), disease characteristics (e.g., cancer type, immune environment), and RPT dosing and timing so that combinations with ICI may be optimized prior to advanced phase clinical investigation.

We aimed to determine the dose of RPT and sequencing of administration of combination RPT and ICI therapies resulting in effective anti-tumor immune responses through comparative studies of ^90^Y-, ^177^Lu-, and ^225^Ac-NM600. NM600 is an alkylphosphocholine analog with selective uptake and retention in a wide array of tumors^20^. The capacity of ^90^Y-, ^177^Lu-, and ^225^Ac-NM600 to generate type I interferon (IFN1) responses and immunomodulate the tumor microenvironment has been reported^15,21–23^. Here, we use ^90^Y-, ^177^Lu-, and ^225^Ac-NM600 to examine RPT + ICI combinations in both poorly immunogenic (B78 melanoma) and immunogenic (MC38 colorectal carcinoma) syngeneic murine tumor models. B78, derived from the spontaneous melanocytic tumor B16F10, is characterized as poorly immunogenic given its low levels of MHC-I expression and limited response to immune checkpoint inhibition alone^15,24,25^. MC38, a carcinogen-induced model, exhibits mismatch repair deficiency^26^ and is characterized as immunogenic, having the highest mutational load amongst ten syngeneic tumor models and a propensity for response to anti-CTLA4^27^ and anti-PD-L1 alone^28^.

Using these models and NM600, we sought to evaluate the development and efficacy of anti-tumor immunity in response to differing radionuclides and varied timing of ICI delivery. We further sought to identify key mechanisms of therapeutic response. We hypothesized that ^225^Ac- NM600 may enhance therapeutic response to dual ICI therapy (anti-PD-L1 and anti-CTLA4) to a greater extent than ^90^Y- or ^177^Lu-NM600 in syngeneic murine tumor models and that the immunogenicity of the tumor model may impact the optimal radioisotope for antitumor responses.

## RESULTS

### Murine tumor models exhibit tumor growth delay and survival improvement following low dose ^225^Ac-NM600+ICI

To perform dose-matched comparative immunologic studies of ^90^Y-, ^177^Lu-, and ^225^Ac- NM600, image-based radiation dosimetry studies were performed (NM600 chemical structure: **Figure 1A**). Tumor uptake and dosimetry of ^86^Y/^90^Y-NM600^15^ and ^177^Lu/^225^Ac-NM600 in mice bearing B78 melanoma tumors were reported previously^23^. Maximum intensity projections of PET or SPECT scans showing MC38 selective uptake of ^86^Y-NM600 (imaging surrogate for ^90^Y-NM600) and ^177^Lu-NM600, respectively, are shown in **Figure 1B-C**. Absorbed doses (mean +/- standard deviation) of 2.03 +/- 0.05, 0.50 +/- 0.01, 1.24 +/- 0.08, and 2.11 +/- 0.21 Gy/MBq ^90^Y-NM600 and 0.47 +/- 0.04, 0.08 +/- 0.01, 0.46 +/- 0.05, and 1.40 +/- 0.24 Gy/MBq ^177^Lu-NM600 were determined for blood, bone, spleen, and MC38 tumors, respectively (**Figure 1D-E**). These dose estimates were obtained using a Monte Carlo voxel-based dosimetry method^29^ and serial PET/CT or SPECT/CT data as well as *ex vivo* biodistribution data at one timepoint (**Figure S1A-B**). Estimates of absorbed dose to MC38 and normal tissues for ^225^Ac-NM600 were obtained by S-value calculations based on the Medical Internal Radiation Dose (MIRD) formalism using serial *ex vivo* biodistribution data (**Figure S1C**)^30^. Absorbed doses of 0.163, 0.248, and 0.437 Gy/kBq ^225^Ac-NM600 were estimated for blood, bone, and MC38 tumor, respectively (**Figure 1F**).

**Figure 1.**
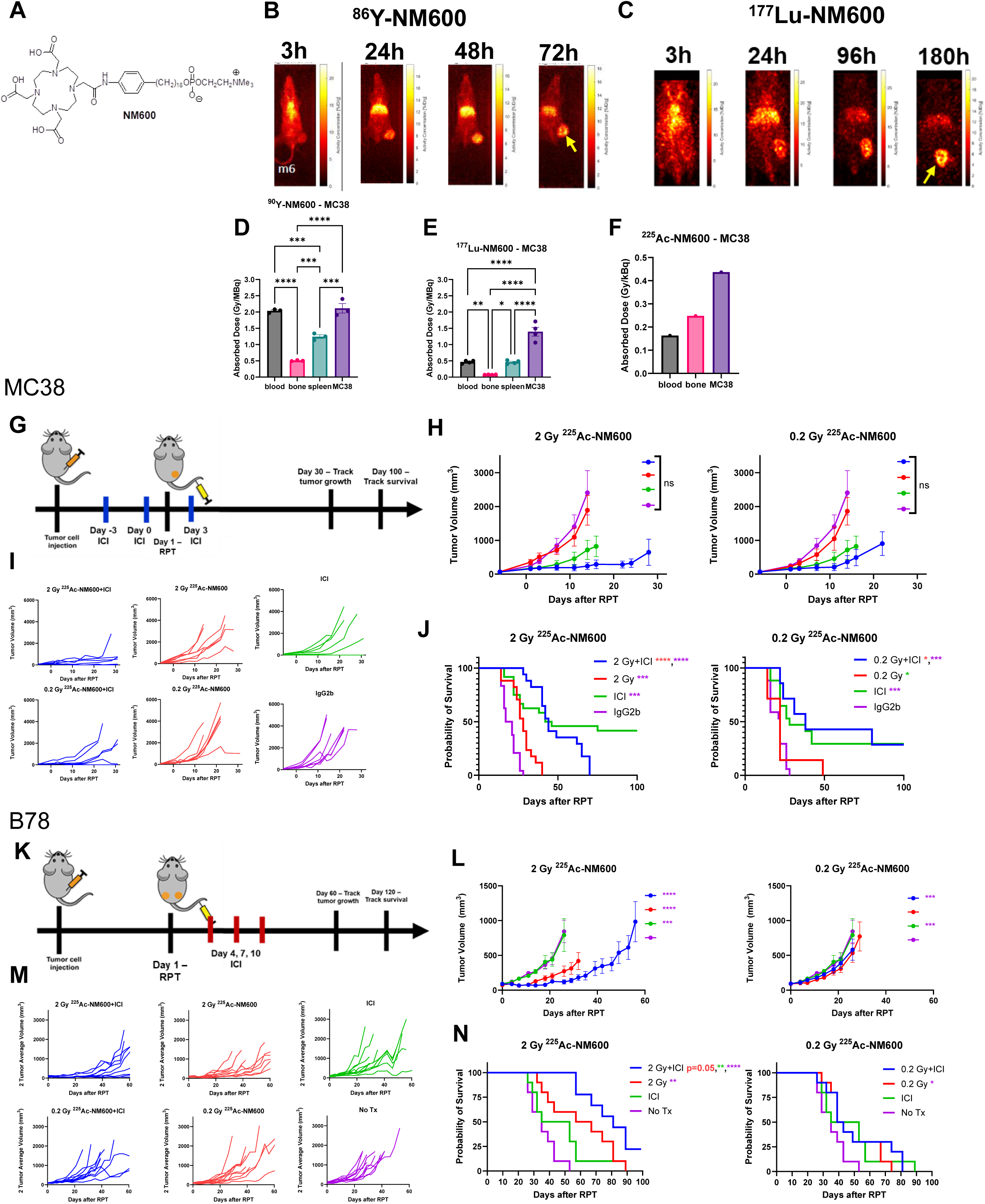
Murine tumor models exhibit tumor growth delay and survival improvement following low dose ^225^Ac-NM600+ICI. A) Chemical structure of NM600. B-C) Maximum intensity projections (MIPs) of B) serial PET/CT images 3h, 24h, 48h, and 72h post injection of 9.25 MBq ^86^Y-NM600 or C) serial SPECT/CT images 3h, 24h, 96h, and 180h post injection of 18.5 MBq ^177^Lu-NM600 for MC38 colorectal carcinoma tumor-bearing mice (tumor indicated by arrow) D-F) Tissue dosimetry performed using Monte Carlo methods (n=3: ^90^Y-NM600; n=4: ^177^Lu- NM600; n=9: ^225^Ac-NM600). G) Treatment scheme for MC38 ^225^Ac-NM600 dose *in vivo* therapy studies. MC38 tumor-bearing mice were randomized to IgG2b control on days -5/0/5 (IgG2b), or 0.2 Gy (0.4625 kBq), 2 Gy (4.625 kBq) ^225^Ac-NM600, or no RPT on day 1 +/- dual anti-CTLA4 and anti-PD-L1 on days -3/0/3 (ICI). H-J) Effects of ^225^Ac-NM600 dose on tumor growth and overall survival. K) Treatment scheme for B78 ^225^Ac-NM600 dose *in vivo* therapy studies. B78 two tumor-bearing mice were randomized to untreated control (No Tx), 0.2 Gy (1.85 kBq), 2 Gy (18.5 kBq) ^225^Ac-NM600, or no RPT on day 1 +/- dual anti-CTLA4 and anti-PD-L1 on days 4/7/10 (ICI). L-N) Effects of ^225^Ac-NM600 dose on tumor growth and overall survival. H-I) N=7: all groups. J) n=17: 2 Gy ^225^Ac-NM600 +/- ICI, ICI, IgG2b; n=7: 0.2 Gy ^225^Ac-NM600 +/- ICI. L-N) N=9: 2 Gy ^225^Ac-NM600+ICI; n=10: all other treatment groups. One-way ANOVA with Tukey’s HSD post hoc test was used to compare ^90^Y-NM600 and ^177^Lu- NM600 absorbed doses between tissues. Log-rank test was used to compare survival.

Blood and additional normal tissue toxicity assessments for ^90^Y-NM600 and ^177^Lu- NM600 have previously been reported^22,31^. The acute toxicity profile for the maximum tolerated activity (MTA) of ^225^Ac-NM600 in naïve C57BL/6 mice (18.5 kBq) is shown in **Figure S2**. ^225^Ac-NM600 could be delivered to naïve mice without physical signs of toxicity during the therapy studies but induced transient cytopenias (**Figure S2**). We also examined the cytotoxicity of ^225^Ac relative to EBRT in both MC38 and B78, observing an expected high relative biological effectiveness (RBE) of ^225^Ac (**Figure S3**). Given that ^225^Ac-NM600 was tolerable with evidence of hematologic recovery and stable weight, we began testing ^225^Ac-NM600 in combination with ICI (**Figure 1G-N**).

In MC38 (**Figure 1G-J, Tables S1-S3)** and B78 (**Figure 1K-N, Tables S4-S6**), dose finding studies were conducted to determine a dose of ^225^Ac-NM600 that could be administered in each tumor model with a therapeutic benefit. From our prior observation that a 2 Gy mean tumor dose was optimal for response to ^90^Y-NM600 + anti-CTLA-4^15^, we investigated 2 and 0.2 Gy ^225^Ac-NM600 in MC38 and B78. B78 has lower uptake of ^225^Ac-NM600 (0.10 Gy/kBq)^23^ than MC38 (0.437 Gy/MBq), and an estimated tumor absorbed dose greater than 2 Gy could not be achieved without exceeding toxicity limits. For both tumor models, 2 Gy ^225^Ac-NM600 was tolerable and demonstrated an immune contribution to tumor response (^225^Ac-NM600+ICI vs. ^225^Ac-NM600 alone, overall survival) (**Figure 1H-J, L-N**). Observed differences in the patterns of response led us to hypothesize that the effects of RPT+ICI may be different based on the immunogenicity of the tumor model.

### ^90^Y-NM600 + early/intermediate-timed ICI enhances therapeutic response in the immunogenic MC38 colorectal carcinoma model

Three different timings of ICI (dual anti-CTLA4 and anti-PD-L1) were investigated relative to treatment day 1 injection of activities of ^90^Y-NM600, ^177^Lu-NM600, or ^225^Ac-NM600 that were estimated from dosimetry to deliver 2 Gy mean tumor dose. ICI delivery was defined as early (days -3/0/3), intermediate (days 4/7/10), or delayed (days 11/14/17). The intermediate-timed ICI has been shown to be safe and effective in combination with all three RPT compounds and EBRT in murine tumor models^15,21,23,32^. Considering the open clinical question of optimal sequencing of RT+ICI and effects of radioisotope half-life on tumor immunomodulation^23^, we compared the sequencing of RPT+ICI (**Figure 2**; 16 treatment groups, repeated two separate times). Comparisons between varied ICI timings are shown in **Figure 2F, H** and **Tables S7-S13,** and comparisons between varied radioisotopes are shown in **Figure 2G, I** and **Tables S14-S20**.

**Figure 2.**
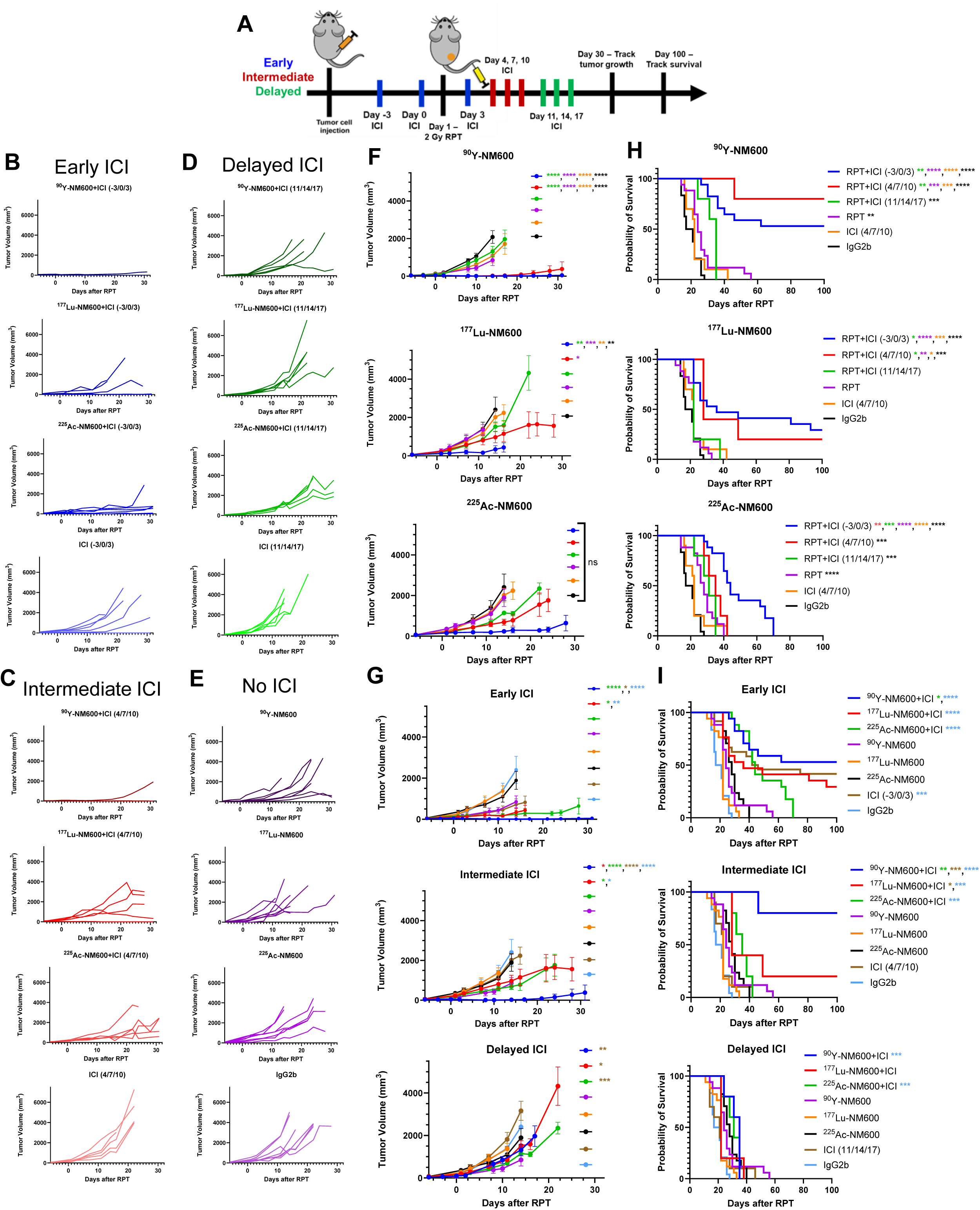
^90^Y-NM600 + early/intermediate-timed ICI enhances therapeutic response in the immune sensitive MC38 colorectal carcinoma model. A) Treatment regimen options for *in vivo* therapy studies. Mice either received early (days -3/0/3), intermediate (days 4/7/10), or delayed (days 11/14/17) dual anti-CTLA4 and anti-PD-L1 relative to RPT on day 1. B-I) MC38 tumor-bearing mice were randomized to one of 16 treatment groups: IgG2b control on days -5/0/5 (IgG2b), dual anti-CTLA4 and anti-PD-L1 (ICI) on days -3/0/3, days 4/7/10, or days 11/14/17, 2 Gy ^90^Y-NM600 (0.925 MBq), ^177^Lu-NM600 (1.85 MBq), or ^225^Ac-NM600 (4.625 kBq) on day 1, or combination 2 Gy ^90^Y-, ^177^Lu-, ^225^Ac-NM600 on day 1 with dual ICI on days -3/0/3, days 4/7/10, or days 11/14/17. B-D) Individual MC38 tumor growth curves corresponding to F & G. F, H) Effects of varied ICI timing for each radionuclide on tumor growth and overall survival. G, I) Effects of varied radionuclide for each ICI timing of combination therapy on tumor growth and overall survival. ^90^Y-, ^177^Lu-, ^225^Ac-NM600, and no RPT + early ICI extended overall survival compared to IgG2b. B-G) N=7: ^90^Y-, ^177^Lu-, ^225^Ac-NM600 +/- ICI −3/0/3, ICI −3/0/3, IgG2b; n=5: ^90^Y-, ^177^Lu-, ^225^Ac-NM600 + ICI 4/7/10 or + ICI 11/14/17, ICI 4/7/10, ICI 11/14/17. H-I) N=24: ICI −3/0/3, IgG2b; n=17: ^90^Y-, ^177^Lu-, ^225^Ac-NM600 +/- ICI −3/0/3; n=10: ICI 4/7/10, ICI 11/14/17; n=5: ^90^Y-, ^177^Lu-, ^225^Ac-NM600 + ICI 4/7/10 or + ICI 11/14/17. Log-rank test was used to compare survival.

^90^Y- and ^177^Lu-NM600 + early ICI reduced MC38 growth rate and prolonged overall survival relative to all other groups. For ^225^Ac-NM600, overall survival prolongation was observed following only ^225^Ac-NM600 + early ICI (**Figure 2F, H**). For early ICI, ^90^Y- NM600+ICI slowed tumor growth rate and prolonged overall survival compared to ^225^Ac- NM600+ICI (**Figure 2G, I**). For intermediate-timed ICI, ^90^Y-NM600 combination therapy showed the greatest therapeutic benefit. For delayed ICI, there were few differences between any RPT+ICI combination therapy and the control groups (**Figure 2G, I**). Rates of complete response (CR) for all mice included in **Figure 2** therapy studies are shown in **Figure S4A-4**. **Figure S4E** demonstrates the rates at which mice rendered disease-free subsequently rejected re-engraftment on the contralateral flank with the same tumor cell line they had been cured from (as a measure of immunologic memory), relative to age-matched naïve control mice. This re-engraftment was performed on day 100 after initial RPT treatment and rates of rejection were: ^90^Y-NM600+early ICI: 8/9; ^90^Y-NM600+intermediate ICI: 4/4; ^177^Lu-NM600+early ICI: 5/5; ^177^Lu-NM600+intermediate ICI: 1/1; early ICI: 8/10; naïve C57BL/6 mice: 0/15.

### ^225^Ac-NM600 + early ICI prolongs overall survival in the poorly immunogenic B78 melanoma model

We next turned to a poorly immunogenic tumor model, B78 melanoma^24,25^, to test whether the immunogenicity of a tumor model may impact patterns of response to RPT+ICI (**Figure 3**; 14 treatment groups). Comparisons between varied ICI timings are shown in **Figure 3F, H** and **Tables S21-S27,** and comparisons between varied radioisotopes are shown in **Figure 3G, I** and **Tables S28-S34**. For each radioisotope delivering 2 Gy mean tumor dose, early ICI administration led to a statistically significant reduction in tumor growth rate and prolongment of overall survival compared to RPT monotherapy and an untreated control group. ^225^Ac- NM600+ICI groups demonstrated the slowest tumor growth rates for each ICI timing. ^225^Ac- NM600+delayed ICI resulted in a significant extension of overall survival compared to ^90^Y- NM600+delayed ICI. **Figure S5** shows the complete response and rechallenge rejection rates for B78 tumor cells injected on the contralateral flank of the complete responders or naïve mice on day 100: ^90^Y-NM600+early ICI: 2/2; ^90^Y-NM600+intermediate ICI: 2/2; ^177^Lu-NM600+early ICI: 1/1; ^225^Ac-NM600+intermediate ICI: 1/1; naïve C57BL/6 mice: 0/5.

**Figure 3.**
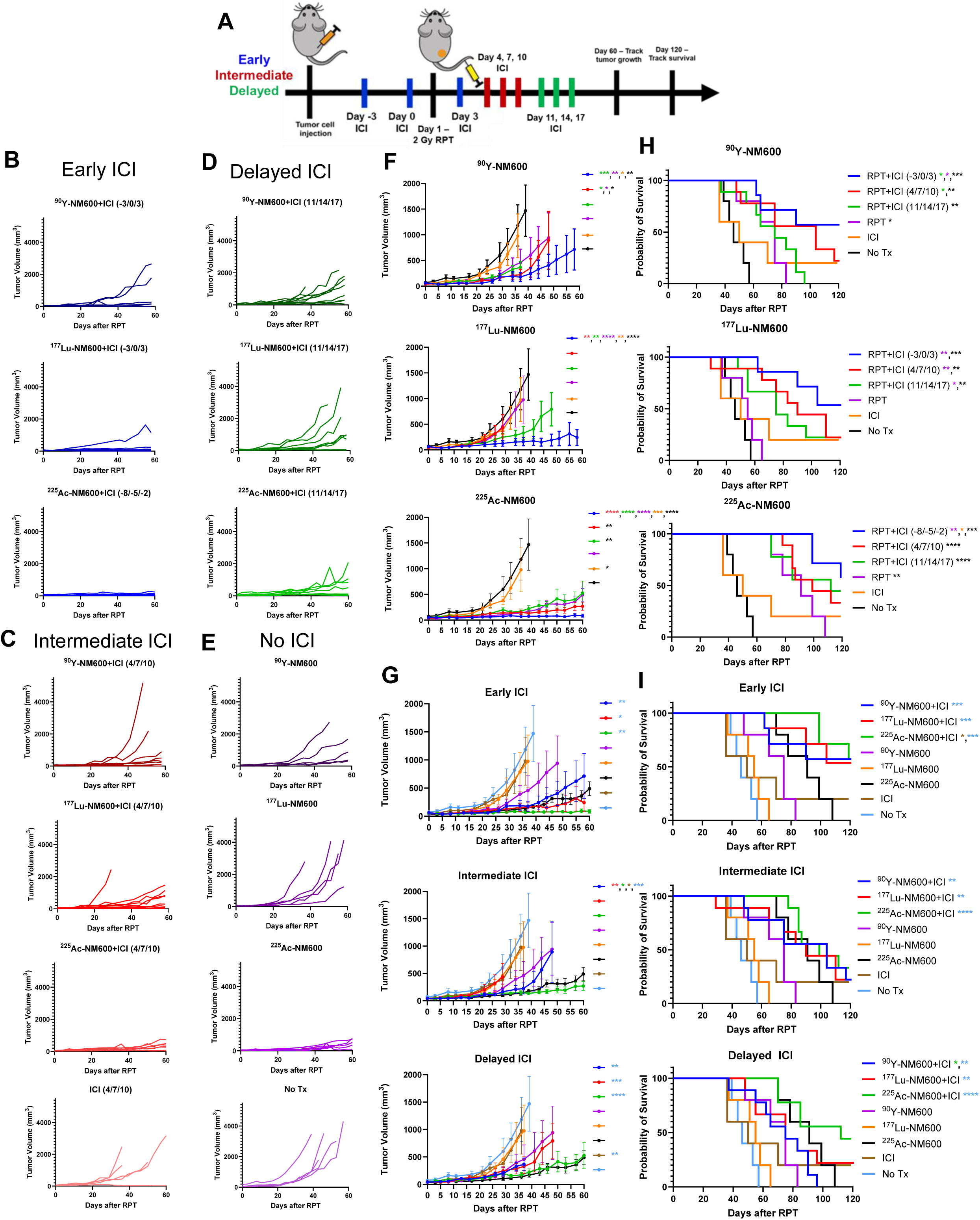
Early ICI prolongs overall survival following RPT+ICI in the immune resistant B78 melanoma model. A) Treatment regimen options for *in vivo* therapy studies. Mice either received early (days -3/0/3), intermediate (days 4/7/10), or delayed (days 11/14/17) dual anti-CTLA4 and anti-PD-L1 relative to RPT on day 1. B-I) B78 tumor-bearing mice were randomized to one of 14 treatment groups: untreated control (No Tx), dual anti-CTLA4 and anti-PD-L1 (ICI) on days 4/7/10, or 2 Gy ^90^Y-NM600 (0.7326 MBq), ^177^Lu-NM600 (1.702 MBq), or ^225^Ac-NM600 (18.5 kBq) on day 1, or combination 2 Gy ^90^Y-, ^177^Lu-, ^225^Ac-NM600 on day 1 with dual ICI on days - 3/0/3 (^90^Y/^177^Lu only), days 4/7/10, or days 11/14/17, or ^225^Ac-NM600+ICI on days -8/-5/-2. B-E) Individual tumor growth curves corresponding to F & G. F, H) Effects of varied ICI timing for each radionuclide on tumor growth and overall survival. G, I) Effects of varied radionuclide for each ICI timing of combination therapy on tumor growth and overall survival. B-I) N=9: ^90^Y-, ^177^Lu-, ^225^Ac-NM600 + ICI 4/7/10 or + ICI 11/14/17; n=7: ^90^Y-, ^177^Lu-, ^225^Ac-NM600 + ICI - 3/0/3; n=5: ^90^Y-, ^177^Lu-, ^225^Ac-NM600, ICI 4/7/10, No Tx. Log-rank test was used to compare survival.

### CD8^+^ T cells are key immune mediators of effector and memory immune response to ^90^Y-, ^177^Lu-, and ^225^Ac-NM600 + ICI

In B78 melanoma, we have shown the necessity of T cells for the therapeutic response to ^90^Y-NM600+anti-CTLA4^15^. Separately, in response to immunotherapy, including ICIs, CD8^+^ T cells have been shown to be major mediators of the antitumor and memory response in mice bearing MC38 tumors^33–35^. Radiation therapy directed against MC38 tumors has also been shown to enhance the antitumor effect of adoptively transferred CD8^+^ T cells^36^. In MC38 colorectal carcinoma, the therapeutic benefit of ^90^Y-, ^177^Lu-, and ^225^Ac-NM600+ICI as compared to RPT monotherapy was abrogated when CD8^+^ T cells were depleted (**Figure 4A-K, Tables S35-S41**). This suggests the necessity of functional CD8^+^ T cells for mediation of antitumor responses to these combination systemic therapies in mice bearing MC38 tumors.

**Figure 4.**
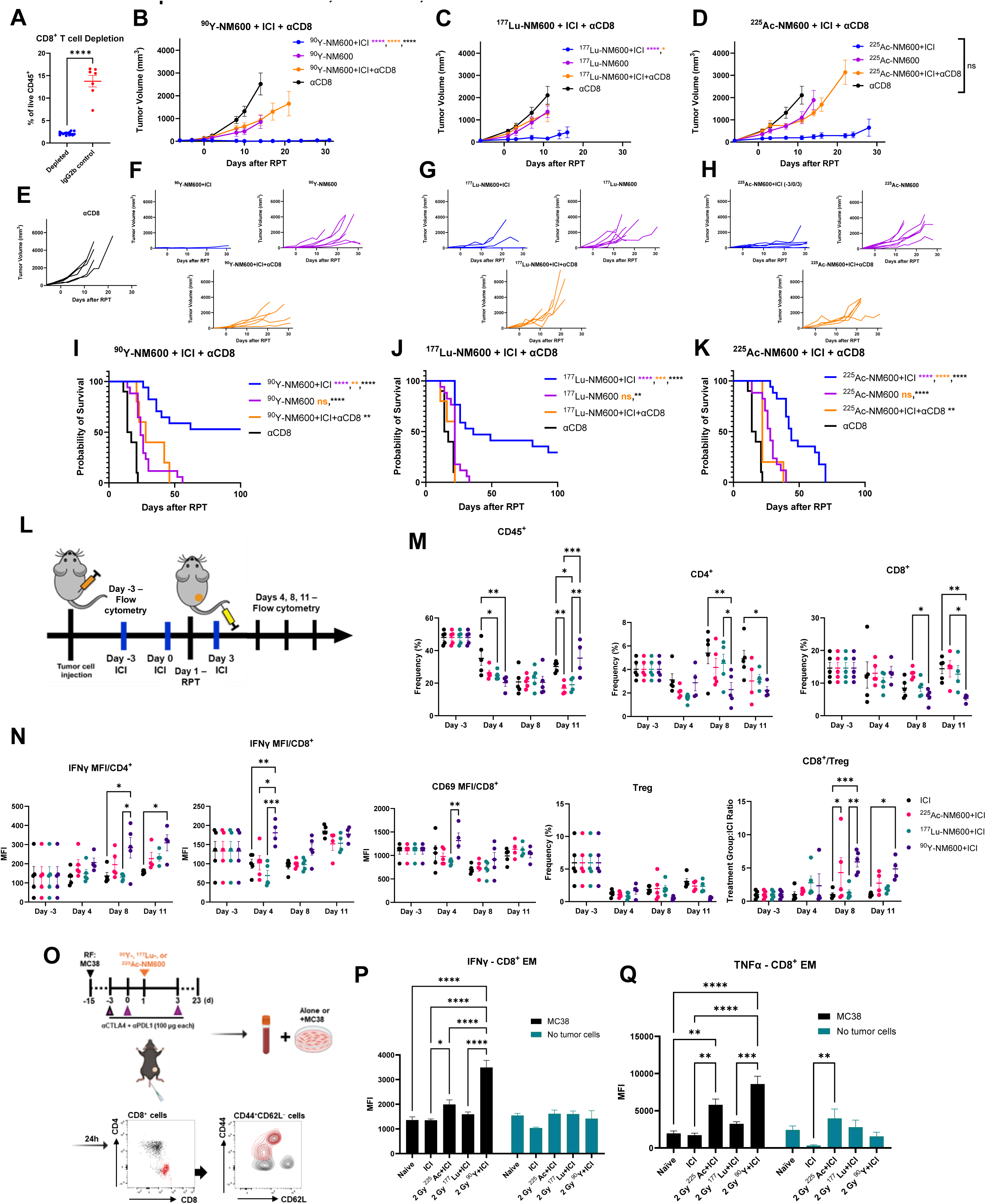
CD8^+^ T cells are key immune mediators of effector and memory immune response to ^90^Y-, ^177^Lu-, and ^225^Ac-NM600 + ICI. A) Confirmation of CD8^+^ depletion by flow cytometry. B-K) MC38 tumor-bearing mice were randomized to receive 2 Gy ^90^Y-, ^177^Lu-, or ^225^Ac-NM600 + ICI (anti-CTLA4 + anti-PD-L1 on days -3/0/3) + αCD8 (anti-CD8 on days -5/0/5), RPT+ICI, RPT alone, or αCD8 alone. Effects of CD8^+^ depletion on tumor growth (B-H) and overall survival (I-K) are shown. L) Flow cytometry treatment scheme. MC38 tumor-bearing mice were randomized to receive either 2 Gy ^90^Y-, ^177^Lu-, or ^225^Ac-NM600 + ICI (anti-CTLA4 and anti-PD-L1) on days -3/0/3, ICI alone, or untreated control (No Tx). M-N) Tumors were harvested on days -3, 4, 8, or 11 and dissociated. The tumor immune infiltrate was analyzed by flow cytometry. O) Co-culture experiment scheme. P-Q) TNFα and IFNγ expression were measured by intracellular flow cytometry staining and analysis of peripheral CD8^+^ effector memory cells (CD8^+^CD44^+^CD62L^-^) harvested from naïve C57BL/6 mice or treated MC38 tumor-bearing mice following 24h of co-culture with MC38 or no tumor cells. A) N=15: depleted; n=7: IgG2b control. B-H) N=7: ^90^Y-, ^177^Lu-, ^225^Ac-NM600 +/- ICI; n=5: ^90^Y-, ^177^Lu-, ^225^Ac- NM600+ICI+αCD8, αCD8. I-K) N=17: ^90^Y-, ^177^Lu-, ^225^Ac-NM600 +/- ICI; n=10: αCD8; n=5: ^90^Y-, ^177^Lu-, ^225^Ac-NM600+ICI+αCD8. M-N) N=5/treatment group and timepoint except n=4: ^90^Y-NM600+ICI Day 4; ^90^Y-, ^177^Lu-, ^225^Ac-NM600+ICI Day 11. P-Q) N=5: naïve, 2 Gy ^90^Y-NM600+ICI; n=6: ICI, 2 Gy ^177^Lu-NM600+ICI; n=7: 2 Gy ^225^Ac-NM600+ICI. Unpaired two-tailed t-test was used to compare CD8^+^ frequency between groups. One-way ANOVA with Tukey’s HSD post hoc test was used to compare flow cytometry results between treatment groups for each marker and timepoint and TNFα or IFNγ expression between treatment groups within each co-culture condition. Log-rank test was used to compare survival.

MC38 tumors were harvested and dissociated for flow cytometry at various timepoints over the course of RPT + early ICI to evaluate longitudinal changes in the tumor immune infiltrate, including pre-treatment (day −3), and one harvest timepoint at the half-life of each radioisotope: day 4 (^90^Y half-life: 2.7 days), day 8 (^177^Lu: 6.7 days), and day 11 (^225^Ac: 10 days), following 2 Gy RPT administration on day 1 (**Figure 4L-N**). Overall, the frequency of CD45^+^ tumor infiltrating immune cells was similar with or without RPT, with a slight reduction in CD45^+^ cell frequency in the ^177^Lu- and ^225^Ac-NM600+ICI tumors compared to ICI and ^90^Y- NM600+ICI on day 11. The ^90^Y-NM600+ICI treated tumors additionally demonstrated an increased CD8^+^/Treg ratio compared to ICI alone, and increased activation marker expression on CD8^+^ T cells and interferon-γ (IFNγ) production by CD8^+^ and CD4^+^ T cells compared to other groups. These results suggest that ^90^Y-NM600+ICI treatment leads to an inflamed tumor immune microenvironment in MC38 colorectal carcinoma. Notably, all groups had a reduction in Tregs following ICI on days -3/0/3, as expected following treatment with a Treg depleting anti-CTLA4 antibody (IgG2c, clone 9D9).

We next aimed to determine whether a tumor-specific memory response was elicited following RPT+ICI. To do this, we investigated the functionality of peripheral CD8^+^ effector memory (EM) cells 23 days after 2 Gy mean tumor dose from ^90^Y-, ^177^Lu-, or ^225^Ac-NM600 + ICI or ICI alone compared to CD8^+^ EM cells isolated from naïve C57BL/6 mice (**Figure 4O-Q**). In this assay, peripheral blood cells were co-cultured with or without MC38 tumor cells for 24 hours, and flow cytometry was performed to measure TNFα and IFNγ effector cytokine production by CD8^+^ EM cells (CD8^+^CD44^+^CD62L^-^). Overall, ^90^Y-NM600+ICI resulted in the greatest induction of TNFα and IFNγ compared to other treatment groups, and treatment with ^177^Lu-NM600 did not induce increased TNFα and IFNγ production by CD8^+^ EM cells. There were few or no differences in the level of TNFα and IFNγ production by peripheral CD8^+^ EM cells between the groups in the absence of MC38 co-culture, consistent with tumor-specific activation.

### Long term immune memory response observed following ^225^Ac-NM600 dose escalation in the immunogenic MC38 tumor model

In prior studies we had observed no improvement in tumor response to ^90^Y-NM600 RPT+ICI combo with dose escalation beyond 2 Gy of ^90^Y-NM600 - a low LET, longer range RPT^15^. Given that dose escalation of ^225^Ac-NM600 to a mean tumor dose of 8 Gy was tolerable with evidence of hematologic recovery and stable weight in mice bearing MC38 tumors, we began testing ^225^Ac-NM600 in combination with ICI in MC38 (**Figure 5A-B**). A dose-dependent increase in TNFα and IFNγ production by peripheral CD8^+^ EM cells following 0.2, 2, or 8 Gy mean tumor dose from ^225^Ac-NM600 + ICI was observed (**Figure 5C**).

**Figure 5.**
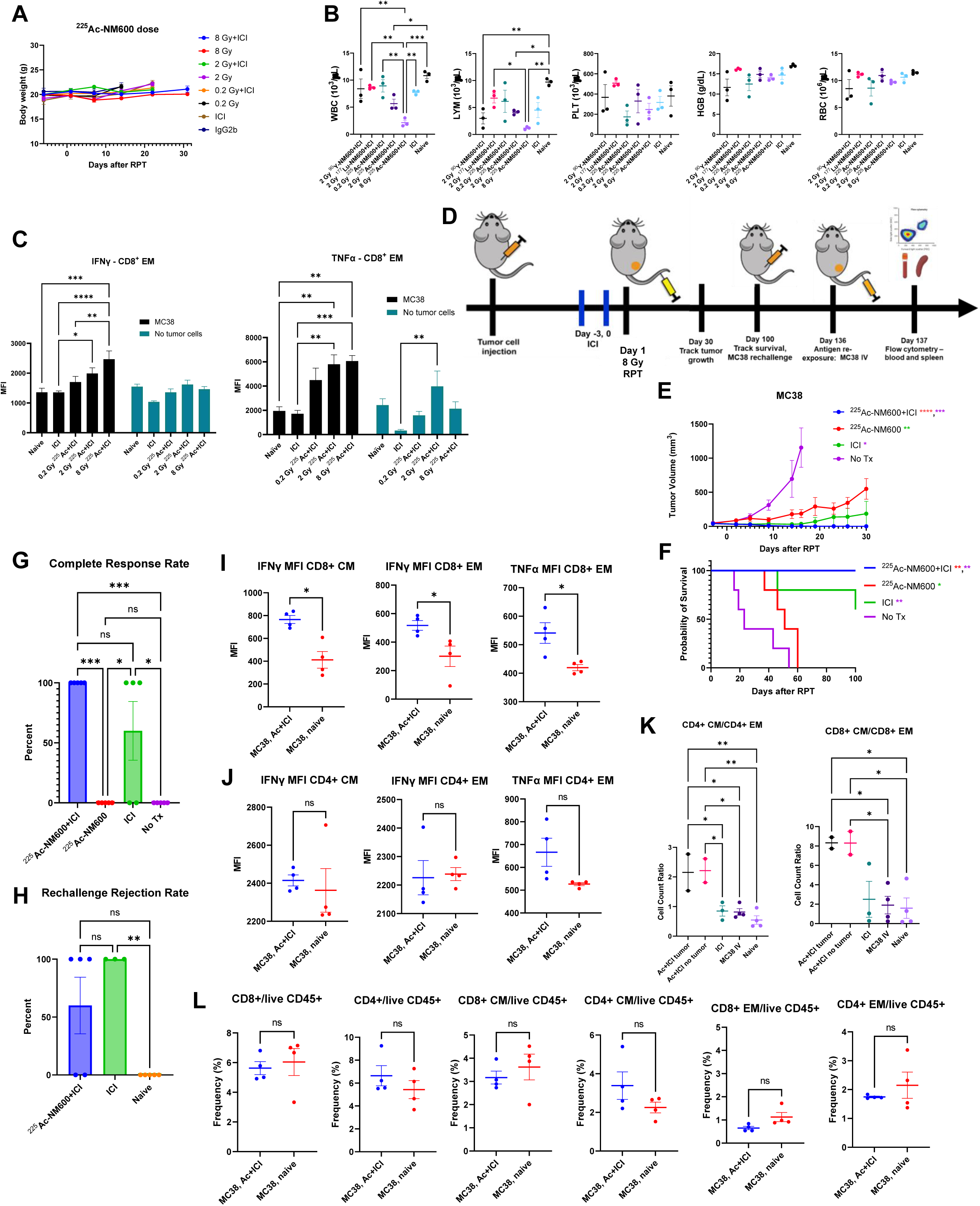
Long term immune memory response observed following ^225^Ac-NM600 dose escalation in the immunogenic MC38 tumor model. A) MC38 tumor-bearing mice were randomized to IgG2b control on days -5/0/5 (IgG2b), or 0, 0.2 Gy (0.4625 kBq), 2 Gy (4.625 kBq), 8 Gy (18.5 kBq) ^225^Ac-NM600 on day 1 +/- dual anti-CTLA4 and anti-PD-L1 on days - 3/0/3 (ICI). Overall mouse body weights over time. B) Complete blood counts on day 23 following RPT day 1 + ICI days -3/0/3. WBC: white blood cells, LYM: lymphocytes, PLT: platelets, HGB: hemoglobin, RBC: red blood cells. C) Samples from co-culture experiment depicted in **Figure 4O**. TNFα and IFNγ expression were measured by intracellular flow cytometry staining and analysis of peripheral CD8^+^ effector memory cells (CD8^+^CD44^+^CD62L^-^) harvested from naïve C57BL/6 mice or dual ICI +/- 0.2, 2, or 8 Gy ^225^Ac-NM600 treated MC38 tumor-bearing mice following 24h of co-culture with MC38 or no tumor cells. D) Scheme for MC38 immune memory response experiment. G-K) MC38 tumor-bearing mice received 8 Gy ^225^Ac-NM600 + ICI (anti-CTLA4 and anti-PD-L1 on days -3, 0), 8 Gy ^225^Ac-NM600 alone, ICI alone, or untreated control (No Tx). G) CR rates. H) All CR mice and naïve controls were rechallenged with 1*10^6^ MC38 cells on the left flank on day 100. I-J, L) Flow cytometry of blood from CR ^225^Ac-NM600+ICI mice (MC38, Ac+ICI) or naïve C57BL/6 mice (MC38, naïve) 24h following IV injection of 2*10^5^ MC38 tumor cells. K) Flow cytometry of splenocytes from CR ^225^Ac-NM600+ICI mice that did not reject rechallenge (Ac+ICI tumor), ^225^Ac-NM600+ICI mice that did reject rechallenge (Ac+ICI no tumor), CR ICI mice (ICI), naïve C57BL/6 mice following IV injection of 2*10^5^ MC38 tumor cells (MC38 IV), and completely naïve C5BL/6 mice (Naïve). A) N=7: all groups. B) N=5: naïve, 2 Gy ^90^Y-NM600+ICI; n=6: ICI, 2 Gy ^177^Lu-NM600+ICI; n=7: 0.2, 2, 8 Gy ^225^Ac-NM600+ICI. C) N=5: naïve; n=6: ICI; n=7: 0.2, 2, 8 Gy ^225^Ac-NM600+ICI. E-G) N=5/treatment group. H) N=5: ^225^Ac-NM600+ICI, naïve; n=3: ICI. I-J, L) N=4/treatment group. K) N=4: MC38 IV, Naïve; n=3: ICI; n=2: Ac+ICI tumor, Ac+ICI no tumor. One-way ANOVA with Tukey’s HSD post hoc test was used to compare ^90^Y-NM600 and ^177^Lu-NM600 One-way ANOVA with Tukey’s HSD post hoc test was used to compare blood counts, CR rates, rechallenge rejection rates, and spleen flow cytometry between groups and TNFα or IFNγ expression between treatment groups within each co-culture condition. Unpaired two-tailed t-test was used to compare blood flow cytometry between groups. Log-rank test was used to compare survival.

We sought to investigate whether a memory response could be observed following 8 Gy mean tumor dose from ^225^Ac-NM600 + ICI. Mice bearing MC38 colorectal carcinoma were treated with ^225^Ac-NM600 on day 1 +/- ICI on days -3/0 (**Figure 5D-L, Tables S42-S44**). Combination therapy significantly prolonged overall survival compared to RPT monotherapy and control (**Figure 5F**). ^225^Ac-NM600+ICI enhanced the CR rate of primary MC38 tumors (5/5 CR) relative to RPT (0/5 CR) and ICI (3/5 CR) monotherapies (**Figure 5G**). CRs were rechallenged with injection of MC38 cells in the flank on day 100. However, only 3/5 ^225^Ac- NM600+ICI CR mice rejected tumor rechallenge as compared to 3/3 ICI CR mice (**Figure 5H**).

From mice showing rejection of this rechallenge, blood and splenocyte cell populations were evaluated by flow cytometry 24 hours following an intravenous injection of MC38 tumor cells on day 136 after RPT. Flow cytometry revealed a significant increase in activation of circulating CD8^+^ memory T cells by IFNγ and TNFα expression compared to naïve mice following MC38 re-exposure for all ^225^Ac-NM600+ICI mice regardless of rechallenge response (**Figure 5I**). No CD4^+^ memory cell activation differences were observed relative to age-matched naïve mice (**Figure 5J**), suggesting the memory response may be mediated by CD8^+^ cells. Additionally, we noted increased CD4^+^ and CD8^+^ central memory (CM)/EM ratios in spleens of all ^225^Ac-NM600+ICI CR mice as compared to ICI monotherapy CR mice (**Figure 5K**), regardless of whether the CR mice rejected rechallenge (Ac+ICI no tumor) or not (Ac+ICI tumor). Increased CD8^+^ CM/EM cell ratios have been reported to be associated with long-term immunologic memory^37^. Circulating lymphocyte percentages were maintained in ^225^Ac- NM600+ICI treated mice (**Figure 5L**). Thus, long-term memory immune memory responses were observed following ^225^Ac-NM600 dose escalation in the immune sensitive MC38 tumor model.

### Tumor immune cell infiltrate and gene expression changes following ^90^Y-, ^177^Lu-, ^225^Ac- NM600, or EBRT + ICI combination therapy in B78 melanoma

In the immune resistant B78 melanoma, we sought to identify tumor immune microenvironment changes following ^90^Y-, ^177^Lu-, or ^225^Ac-NM600 + ICI combination therapy that would explicate the relative therapeutic benefit of alpha-emitting low dose ^225^Ac-NM600 compared to low dose ^90^Y-NM600 observed in **Figure 3**. We performed paired single cell (sc)RNA-sequencing (seq) and T cell receptor (TCR) seq among tumor-infiltrating immune cells from mice bearing B78 melanoma tumors receiving 2 Gy mean tumor dose from either ^90^Y-, ^177^Lu-, ^225^Ac-NM600, or EBRT + early ICI or ICI alone. Tumors were harvested and 3 tumors/treatment group were pooled for CD45^+^ cell isolation and scRNA-seq and TCR seq on treatment day 15 (**Figure 6A**). **Figure S6** shows the tumor growth curves of the B78 melanoma tumors harvested and pooled for scRNA-seq analysis. Following filtering and quality control, data were analyzed for 40,672 cells with 11,312 mean reads per cell. Dimensionality reduction and graph-based clustering of all cells were performed to distinguish cells with unique transcriptional profiles. UMAPs showing identical cell populations (overall cell population analyzed) with each treatment group highlighted are shown in **Figure 6B**.

**Figure 6.**
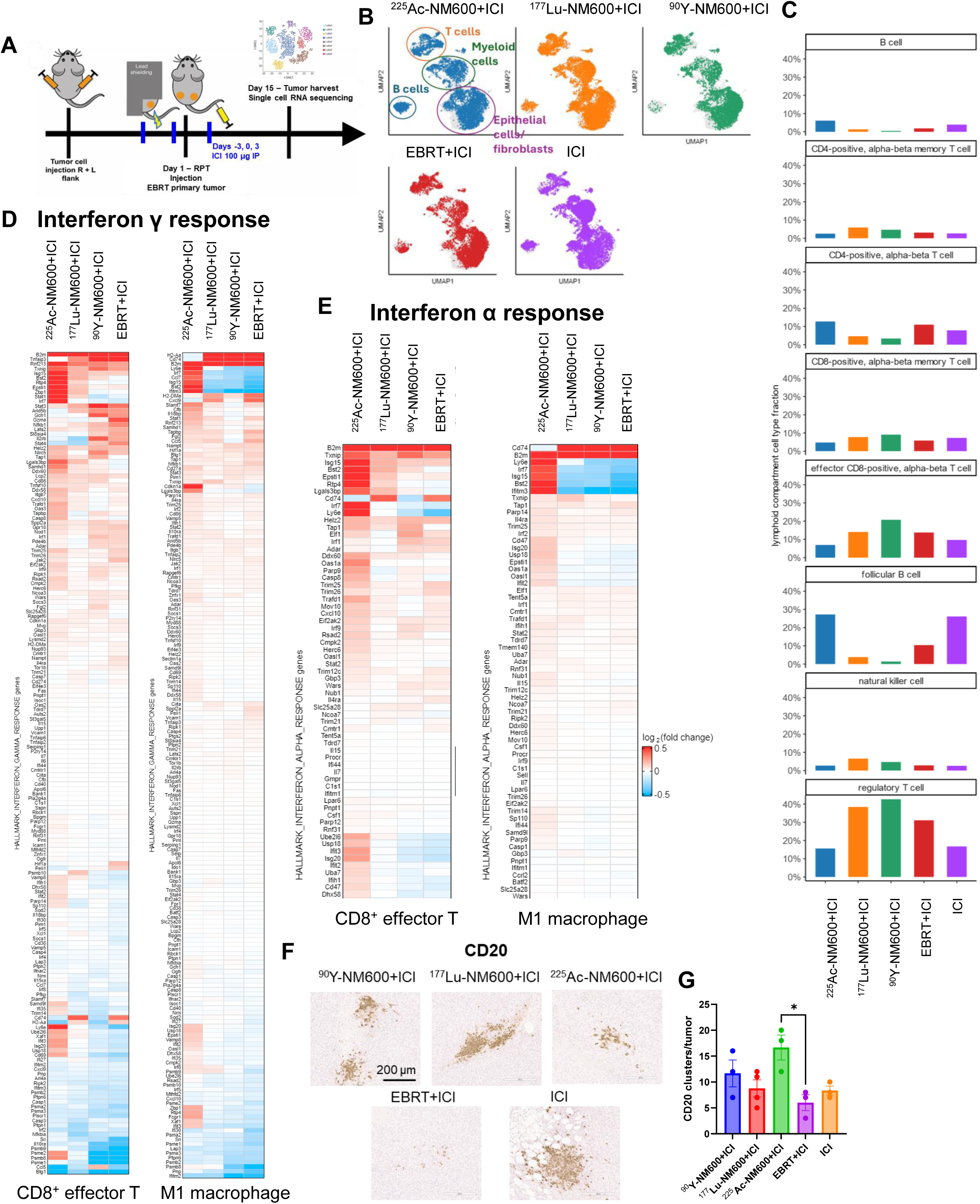
Tumor immune cell infiltrate and gene expression changes following ^90^Y-, ^177^Lu-, ^225^Ac-NM600, or EBRT + ICI combination therapy in B78 melanoma. A) Treatment scheme. B78 tumor-bearing mice received either 2 Gy ^90^Y-, ^177^Lu-, ^225^Ac-NM600 or EBRT + ICI (anti-CTLA4 and anti-PD-L1) on days -3/0/3 or ICI alone. Tumors were harvested on day 15, dissociated, and immune cells were isolated by Ficoll density gradient centrifugation for single cell RNA sequencing analysis. B) UMAP of single cell RNA sequencing data colored by samples. Each panel shows identical cell population with one sample in color and all the others in gray. Regions containing T cells, myeloid cells, B cells, and epithelial cells/fibroblasts are indicated. C) Cell type fractions in lymphoid compartment. Cell types having >5% fraction in at least one sample are represented. D-E) Differential gene expression heat map for interferon γ (D) and interferon α (E) response genes for CD8^+^ effector T cells and M1 macrophages. The color of each box represents the log_2_(fold change) of gene expression of ^225^Ac-, ^177^Lu-, ^90^Y-NM600, or EBRT+ICI compared to ICI alone. N=3 mice/treatment group pooled into one sample/treatment group for single cell RNA sequencing analysis. F) B78 tumor photomicrographs for B cell marker CD20 at day 15 following 2 Gy ^90^Y-, ^177^Lu-, ^225^Ac-NM600 or 2 Gy EBRT + ICI or ICI alone; brown = positive immunolabeling. G) Quantification of CD20^+^ cell clusters/tumor. Cell cluster: ≥ 5 cells. N=3 tumors/treatment group. One-way ANOVA with Tukey’s HSD post hoc test was used to compare CD20 IHC quantification between treatment groups.

Lymphoid cell type fractions are shown in **Figure 6C**. Of note, frequency of follicular B cells, which participate in T-cell-dependent antibody responses, were increased in the ^225^Ac- NM600+ICI and ICI alone treatment groups. The ^225^Ac-NM600 follicular B cells had a 2.57-fold change increase compared to ICI alone in expression of MHC-I gene, *H2-K1*. Frequencies of both CD8^+^ effector T cells and Tregs were highest in the ^90^Y-NM600+ICI treatment group. CD8^+^/Treg ratios for each treatment group were similar between groups in this analysis: ^225^Ac- NM600+ICI: 0.44, ^177^Lu-NM600+ICI: 0.37, ^90^Y-NM600+ICI: 0.49, EBRT+ICI: 0.44, ICI: 0.58.

Differential gene expression analysis was performed to evaluate the relative expression of genes in each combination RT+ICI treatment vs. ICI alone. These results for CD8^+^ effector T cells and M1 macrophages for genes involved in IFNγ and IFNα responses are shown in **Figure 6D-E**. In the ^225^Ac-NM600+ICI treatment group several CD8^+^ effector T cell genes were upregulated to a greater degree than other RT+ICI groups: *Rnf213, Txnip, Isg15, Bst2, Rtp4, Epsti1, Zbp1, Stat1, Irf7,* and *Ly6e*. Of these, *Rnf213, Isg15, Rpt4,* and *Stat1* are interferon-induced genes; IRF7 regulates several interferon α genes; and ZBP1 activates both IFN1 responses and NF-κB signaling^38^. This IFN1 activation reflects the reported IFN1 signaling following radiation due to accumulation of cytosolic micronuclei^39^. TXNIP has been reported to be associated with increased tumor infiltrating immune cells and a carcinostatic role overall^40^. Epithelial-stromal interaction 1 (EPST1) is a protein that promotes monocyte adhesion to endothelial cells^41^. BST2 and LY6E are pro-tumor proteins. *B2m*, which encodes β_2_ microglobulin, a component of MHC-I molecules, was upregulated for all groups relative to ICI alone. This increase in expression for all RT+ICI groups reflects the well documented ability of RT to enhance MHC-I expression^42^.

For M1 macrophages, there were several genes upregulated following ^225^Ac-NM600+ICI but downregulated for other groups, including *Ly6e, Irf7, Isg15, Bst2,* and unique from CD8+ effector T cells, *Ccl7* and *Ifitm3*. *Ccl7* has been shown to facilitate antitumor responses to anti-PD-1 through recruitment of classical dendritic cells^43^, and *Ifitm3* encodes an interferon-inducible transmembrane protein. Conversely, both the MHC-II gene (*H2-Aa)* and *Cd74*, involved in formation and transport of MHC-II^44^, and were downregulated following ^225^Ac- NM600+ICI in M1 macrophages, but upregulated in all other RT+ICI groups. Taken together, genes induced by ^225^Ac-NM600+ICI have mixed roles in tumor progression and immune response, with a clear prevalence of MHC-I expression among all RT groups, and IFN1-inducible genes following ^225^Ac-NM600+ICI, consistent with the greater amount of DNA damage induced by the α-emitting radioisotope.

External beam radiation to tumor draining lymph nodes (TDLNs) decreases the effectiveness of RT + anti-CD25 immunotherapy^13^. In metastatic settings, TDLNs may not be readily identifiable or avoidable. Furthermore, it is unknown what effect EBRT may have on other immune structures including tertiary lymphoid structures (TLSs), which have been shown to be critical to antitumor immunity with ICIs^45^. We hypothesize that RPT, by molecularly rather than spatially targeting tumors, could avoid such structures. EBRT+ICI disrupted existing CD20^+^-rich lymphoid clusters in B78 melanoma (**Figure 6F-G**). These CD20^+^ structures were present in a significantly greater number in the ^225^Ac-NM600+ICI tumor, which is the radioisotope with the shortest emission pathlength (<100 μm)^23^. The short tissue pathlength signifies that irradiation is minimized to non-targeted cells in the tumor including stroma and islands of immune cells, as well as neighboring structures including lymph nodes.

### ^225^Ac-NM600+ICI leads to diversified effector T cell responses with key shared clonotypes

With the same cells used for scRNA-seq analysis, we performed TCR seq analysis to gain insight into the functional status of CD8^+^ T cells following ^90^Y-, ^177^Lu-, ^225^Ac-NM600, or EBRT+ICI. After quality control, TCR sequencing analysis provided data for 3,517 cells with productive V-J spanning pairs. UMAPs of all T cells stratified by sample are shown in **Figure 7A**. Clonotype expansion ranges for **Figure 7** are represented in **Figure 7B**. UMAPs stratified by T cell subtype are shown in **Figure 7C**, verifying that most medium and large T cell clones pertain to CD8^+^ effector and memory subtypes. Clonal expansion by T cell subtype and CDR3 sequence diversity measurement (D50) are shown in **Figures 7D and 7E**. ^177^Lu- and ^90^Y- NM600+ICI have more clonal expansion of CD8^+^ effector and memory T cells, with ^225^Ac-NM600 and EBRT+ICI showing more diversity and a higher D50 score for CD8^+^ effector T cells. **Figure 7F** presents the clonal expansion data in a different way, demonstrating the % of cells with clonotypes per expansion. These results indicate that the majority of effector CD8^+^ clones in the ^177^Lu- and ^90^Y-NM600+ICI groups are medium or large clones. These findings are consistent with previously reported CD8^+^ clonal expansion following low dose ^90^Y- NM600+ICI^15^. Finally, shared clonotypes between various subsets of T cells are shown in **Figure 7G-H**, with ^225^Ac-NM600+ICI treated tumors having the greatest number of shared clonotypes between effector and memory CD8^+^ T cells (11), as compared to seven shared clonotypes for the same T cell subsets in each other treatment group. While ^225^Ac-NM600+ICI treated tumors had greater T cell diversity overall, they exhibited shared clonotypes between effector and memory CD8^+^ T cells. This is consistent with an *in situ* vaccine effect and greater priming of an adaptive T cell response with high LET α-particle RPT, as compared to the greater clonal expansion or propagation of existing immune response in the absence of apparent T cell priming with low dose, low LET β-particle RPT.

**Figure 7.**
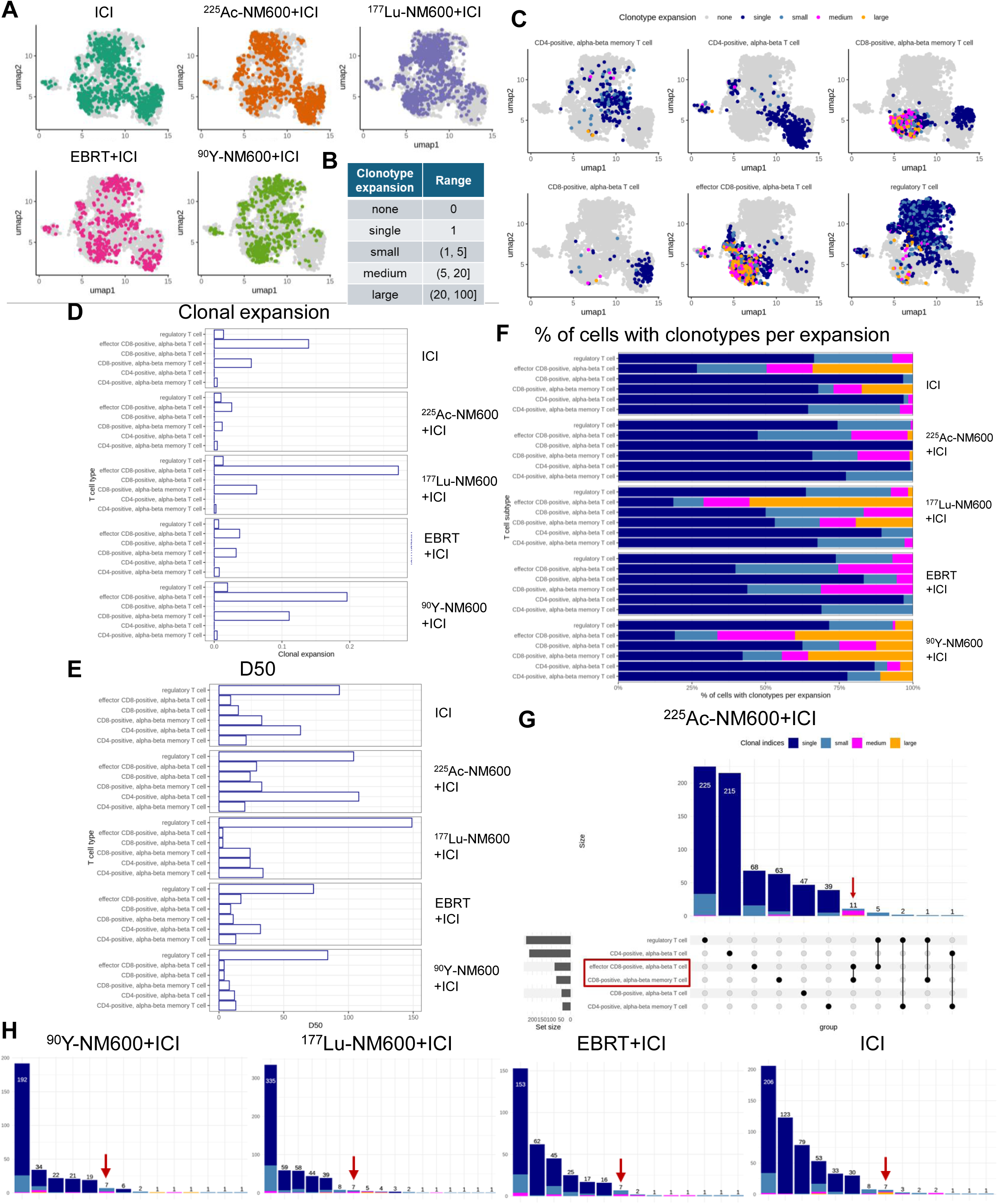
^225^Ac-NM600+ICI leads to diversified effector T cell responses with key shared clonotypes. Paired TCR sequencing analysis performed on samples from B78 tumor-bearing mice used for single cell RNA sequencing. A) UMAP plots of T cells stratified by sample. B) TCR clonotype expansion ranges. C) UMAP plots of T cells stratified by T cell subtype. D) Clonal expansion by treatment group and T cell subtype. E) D50 score by treatment group and T cell subtype. F) Relative abundance of clonal indices for each treatment group and T cell subtype. G) UpSet plot showing shared clonotypes between T cell subtypes for the ^225^Ac- NM600+ICI sample. Red box demarcates CD8^+^ effector and memory subsets; red arrow demarcates # of shared clonotypes between CD8^+^ effector and memory subsets (= 11). H) UpSet plots showing shared clonotypes between T cell subtypes for the ^90^Y-, ^177^Lu-, EBRT+ICI, and ICI alone samples. For each plot, red arrow demarcates # of shared clonotypes between CD8+ effector and memory subsets (= 7 for each sample). N=3 mice/treatment group pooled into one sample/treatment group for TCR sequencing analysis.

## DISCUSSION

Low dose ^90^Y-NM600+ICI was more effective than tumor dose-matched ^225^Ac-NM600 + ICI in the immunogenic MC38 colorectal carcinoma model. This observation was surprising, given that greater STING activation has been observed with ^225^Ac^23^. The MC38 response to RPT alone (**Figure 2E**) was not affected by difference in RBE, suggesting that the therapeutic response in MC38 may be predominantly mediated by tumor-resident immune cells. The higher frequency of tumor infiltrating CD45^+^ cells in the ^90^Y-NM600+ICI group on day 11 may contribute to the superior tumor responses in that group (**Figure 4M**). Additionally, circulating CD8^+^ effector memory cells isolated after 2 Gy mean tumor dose from ^90^Y-NM600+ICI treated mice had higher effector cytokine production than those isolated after the same mean tumor dose from ^225^Ac-NM600+ICI when co-cultured with MC38 tumor cells. This suggests that ^90^Y- NM600+ICI is effective at enhancing the existing effector immune response in immune sensitive MC38 (**Figure 4P-Q**).

The combination of enhanced clonal expansion and preservation of existing tumor immune response following ^90^Y-NM600+ICI may be particularly beneficial in patients with immune-sensitive metastatic cancers. These tumors, like the MC38 model, have an existing degree of immunogenicity (increased tumor mutational burden, increased immune cell infiltration at baseline, increased MHC-I expression, or a combination of these) and may have developed acquired resistance to ICIs. However, response to ICI may be rescued in such settings following RPT+ICI therapy where low dose levels permit immune cell sparing (**Figure 6F-G**), enhanced immune cell infiltration of tumor (**Figure 4M-N**), increased peripheral immune cell activation **(Figure 4P-Q, 5C**), and bolstered immunological memory (**Figure 5I-L**).

Conversely, ^225^Ac-NM600+ICI was more effective than ^90^Y-NM600+ICI and ^177^Lu- NM600+ICI in the poorly immunogenic B78 melanoma model. There are several potential contributing factors to this result. ^225^Ac-NM600 is a high LET radioisotope with a greater RBE in this tumor model as a monotherapy (**Figure 3E**). Additionally, irradiation of B78 melanoma *in vitro* with low dose ^225^Ac was shown to increase expression of type I interferon response-associated genes, which was not observed following ^90^Y or ^177^Lu at the same low dose^23^. Additionally, single cell RNA sequencing and TCR sequencing of B78 tumors following ^225^Ac- NM600+early ICI compared to ^90^Y- and ^177^Lu-NM600+early ICI showed that ^225^Ac-NM600+ICI treated tumors had fewer infiltrating Tregs and greater infiltrating follicular B cells (**Figure 6C**). IFNγ and IFNα response genes were upregulated in the ^225^Ac-NM600+ICI group specifically (**Figure 6D-E**).

Additionally, the short tissue range of α-emitting ^225^Ac-NM600 may result in greater dose heterogeneity and sparing of B cell-rich TLSs compared to β-emitting radioisotopes or EBRT^46^ (**Figure 6F-G**). Radiation dose heterogeneity, tumor infiltrating lymphocytes (TILs), nearby TDLNs, and TLSs have all been implicated in anti-tumor immune responses following RT+ICI^13,45–48^. Low-dose radiotherapy and anti-PD-1 were shown to increase quantity and maturity of TLSs in murine lung cancer, with a strong antitumor effect associated with CD8^+^ number within TLSs^48^. RPT has the potential to maintain and possibly promote TILs, TDLNs, and TLSs by molecularly targeting tumor cells rather than spatially targeting tumors, as with EBRT.

A unique combination of an effector CD8^+^ T cell repertoire with greater diversity overall but increased clonal expansion of shared clonotypes between effector and memory CD8^+^ T cells was noted for the ^225^Ac-NM600+ICI group compared to the ^90^Y- and ^177^Lu-NM600+ICI groups (**Figure 7**). In a report by Fairfax et al., large CD8^+^ T cell clone counts 21 days following treatment, regardless of clonal specificity, were associated with durable responses to combination (anti-CTLA4 and anti-PD-1) ICI in metastatic melanoma patients^49^. The T cell populations analyzed in this study could signal a progressing adaptive immune response. These findings are consistent with ^225^Ac-NM600+ICI leading to an enhanced systemic antitumor immune response in the immune resistant B78 melanoma model. High LET ^225^Ac-NM600 may have a role of priming antitumor immune responses through recruitment of CD8^+^ T cells through type I IFN signaling, whereas low LET may have a larger role in propagating existing immune responses in immune sensitive tumors. All studies here were performed with low-dose RPT. Therefore, it is unknown if low-LET RPT may also prime immune resistant tumors to respond to ICI if administered at higher doses or in a more heterogeneous distribution. Further investigation is warranted with additional RPT agents and tumor models.

Early or intermediate-timed dual anti-CTLA4 and anti-PD-L1 in combination with RPT significantly prolongs overall survival compared to groups receiving delayed dual ICI in both MC38 and B78 (**Figures 2, 3**). Recently, trials have highlighted the importance of timing of RT + ICI to achieve anti-tumor responses ^50,51^ (NCT03519971). In our study, administering a Treg depleting isotype of anti-CTLA4 (Treg depletion shown in **Figure 4N**) before RPT in the early ICI groups may have had an immune priming effect or effect in depleting Tregs^52^. Depleting this immune suppressive population may have aided activated CD8^+^ T cells (**Figure 4P-Q**, **Figure 6D-E**). Additionally, as is shown in **Figure 6**, RPT preserved CD20^+^ tumor immune structures, potentially sparing antigen presentation processes within the tumor and helping to promote clonal expansion of a shared lineage of CD8^+^ effector and memory cells (**Figure 7G**).

This study had several limitations. To perform exploratory investigations of tumor immunomodulation with novel RPT agents, doses, and ICI treatment schedules, syngeneic murine tumor models are a valuable tool. However, murine tumor models do not fully recapitulate the immune microenvironment and metastatic potential of human tumors and disease progression. In this work, we compared equal absorbed doses of radiation delivered to a tumor with the same vector (NM600). However, changing the isotope could alter the biodistribution of the agent, resulting in differing doses for normal tissues, which potentially confounds the immune response. ^225^Ac-NM600, for example, had higher liver uptake and lower tumor:liver uptake ratios than ^90^Y- or ^177^Lu-NM600 (^225^Ac-NM600 tumor:liver dose ratio: 0.23; ^177^Lu- NM600: 1.37; ^90^Y-NM600: 0.66). Given this variation, it is important to use caution when selecting surrogate imaging radioisotopes for dosimetry, as the radioisotope itself may greatly impact the biodistribution and pharmacokinetics of a given RPT agent. Unfortunately, there may not be a better method for comparison of tumor immune effects than using equivalent tumor doses and the same RPT chelator and vector.

Although acute hepatotoxicity was not observed (**Figure S2**), we investigated late toxicities of ^225^Ac-NM600 (**Figure S7**), and noted elevated liver transaminases and serum creatinine, suggesting hepatic and renal dysfunction. One parameter of radiopharmaceuticals that can be adjusted to improve toxicities and lesion:non-lesion residence times is specific activity (A_s_). Frequently, high A_s_ is desired to reduce toxicities associated with certain RPT vectors or with target saturation at low A_s_ ^53^. However, in the case of ^111^I-J591, lowering molar activity of the RPT agent by adding unlabeled antibody increased the lesion residence time and the ratio of lesion:liver residence times^54^. For this reason, we investigated the effect of lowering the A_s_ of ^225^Ac-NM600 (**Figure S7**). The liver uptake of ^225^Ac-NM600 was significantly decreased in the low A_s_ group (0.011 MBq/nmol ^225^Ac-NM600) vs. high A_s_ (0.337 MBq/nmol ^225^Ac-NM600), and the MC38 uptake was not significantly different. However, late (6 month post administration) changes in mouse weight and hepatic and renal toxicity were similar in low and high A_s_ treatment groups. Thus, this approach of lowering the A_s_ did not markedly improve the biodistribution of ^225^Ac-NM600.

In poorly immunogenic tumors with intrinsic resistance to ICIs, combination with high LET RPT may prime adaptive CD8^+^ T cell recognition. Conversely, in immunogenic tumors with acquired resistance to ICIs, low doses of low LET RPT may provide tumor inflammation and tumor cell susceptibility needed to renew clonal expansion of existing tumor-specific CD8^+^ T cells and generate immunologic memory. This study emphasizes the critical importance of understanding the mechanisms of ICI resistance in order to appropriately implement combined modality treatment approaches that may overcome such resistance. In this work, we begin to clarify differential immune effects of distinct radioisotopes when delivered by the same RPT vector. A mechanistic understanding of these effects will be vital to the rational design and optimization of clinical studies combining RPT and ICIs.

## METHODS

### Cell lines and culture

The murine melanoma B78-D14 (B78) cell line, derived from B16 melanoma as previously described, was obtained from Dr. Ralph Reisfeld (Scripps Research Institute) in 2002^55^. The murine colorectal carcinoma MC38 cell line was obtained from American Type Culture Collection (ATCC). B78 cells were grown in a humidified incubator at 37°C with 5% CO_2_, in RPMI-1640 supplemented with 10% FBS, 100 U/mL penicillin, and 100 µg/mL streptomycin; MC38 cells were grown in DMEM supplemented with 10% FBS, 100 U/mL penicillin, and 100 µg/mL streptomycin. Cell line authentication was performed per ATCC guidelines using morphology, growth curves, and Mycoplasma testing within 6 months of use.

### Murine tumor models

Mice were housed and treated under a protocol approved by the Institutional Animal Care and Use Committee at the University of Wisconsin-Madison (protocol: M005670). C57BL/6 mice were purchased at age 6 to 8 weeks from Taconic. Both male and female mice were included in MC38 and B78 therapy studies. Female mice were used for correlate studies.

#### Therapy studies

MC38 tumors were engrafted by subcutaneous flank injection of 1 × 10^6^ tumor cells. B78 tumors were engrafted by subcutaneous flank injection of 2 x 10^6^ tumor cells. Tumor size was determined using calipers and volume approximated as (width^2^ × length)/2. Mice were randomized immediately before treatment by permuted block randomization. Only mice with palpable flank tumors the day before treatment began were included in the study. Treatment began when tumors were well-established, approximately 12 days after tumor implantation for MC38 (∼50-100 mm^3^) and 4 weeks for B78 (∼50-100 mm^3^). The day of EBRT or RPT was defined as “day 1” of treatment. Anti-murine CTLA-4 (IgG2c, clone 9D9, produced by NeoClone) and anti-murine PD-L1 (IgG2b, clone 10F.9G2, BioXCell) were administered by 100 μg intraperitoneal injection on days indicated in figures and figure legends. Mice were euthanized when tumor size exceeded 20 mm in the longest dimension or recommended by an independent animal health monitor for morbidity or moribund behavior. Due to institutional radiation safety protocols and the mice being treated with radioactive isotopes, all investigators were aware of mouse treatment groups. The number of mice in therapy groups was based on anticipated effect size of outcomes to power experiments, determined in collaboration with biostatisticians, and therefore may vary between treatment groups and tumor models. Mouse therapy experiments were repeated in duplicate. Final replicates are presented for tumor response and aggregate data for survival; number of animals per group is indicated in figure legends.

#### CD8^+^ depletion studies

CD8^+^ T cell depletion was performed by injection of anti-murine CD8α (Rat IgG2b, κ; Clone 2.43; BioXCell) or IgG2b control (Rat IgG2b, κ; Clone LTF-2; BioXCell) by 300 µg intraperitoneal injection on days -5, 0, and 5. Depletion was confirmed on treatment day −3 by flow cytometry.

#### Memory studies

Mice without palpable tumors on day 100 were considered complete responders (CR). CR mice were rechallenged with 1*10^6^ MC38 cells or 2*10^6^ B78 cells (whichever cell line matched original tumor) contralateral flank injection on day 100. For each therapy study, n=5 age-matched naïve female C57BL/6 mice were injected with either 1*10^6^ MC38 cells or 2*10^6^ B78 cells as a control. On day 136, CR mice and age-matched naïve controls (n=4) received 2*10^5^ MC38 cells IV. An additional n=4 naïve controls received no cells IV. 24h post antigen re-exposure, blood and splenocytes were harvested for flow cytometry.

### Radionuclides and radiochemistry

^90^Y was purchased as ^90^YCl_3_ from Eckert and Ziegler. ^177^Lu and ^225^Ac were purchased as ^177^LuCl_3_ and solid ^225^Ac(NO_3_)_3_ from Oak Ridge National Laboratory. The radiolabeling of 2- [[hydroxy[[18-[4-[[2-[4,7,10-tris(carboxymethyl)-1,4,7,10-tetraazacyclododec-1- yl]acetyl]amino]phenyl]octadecyl]oxy]phosphinyl]oxy]-N,N,N-trimethyl ethanaminium (NM600) with ^90^Y, ^177^Lu, or ^225^Ac proceeded by mixing the radiometal with NM600 (0.336-3.36 MBq/nmol) in 0.1 M NaOAc buffer (pH 5.5) for 30 min at 90°C, as previously described for ^90^Y-NM600^22^. The labeled compounds were purified using a reverse-phase Waters Oasis HLB Light cartridge (Milford, MA), eluted using 100% ethanol, dried with a stream of N_2_, and reconstituted in 0.9% NaCl with 0.1% v/v Tween 20. Yield and purity were determined with instant thin-layer chromatography using 50 mM EDTA and iTLC-SG –glass microfiber chromatography paper impregnated with silica gel (Agilent Technologies). A cyclone phosphor image reader was used to analyze the chromatograms; the labeled compound remained at the spotting point while free radiometals moved with the solvent front.

### PET/CT imaging

Mice bearing MC38 (n=3 male, n=3 female) flank tumors (100-150 mm^3^) were injected via tail vein with 9.25 MBq of ^86^Y-NM600 and imaged with an Inveon microPET/microCT scanner (Siemens Medical Solutions, Knoxville, TN) at 3, 24, 48, and 72 hours post injection of the radiotracer. For each scan, mice were anesthetized with isoflurane and placed in the prone position on the scanner bed. Sequential CT (80 kVp; 1000 mAs; 220 angles) and static PET scans (80 million coincidence events; time window: 3.432 ns; energy window: 350-650 keV) were collected. A three-dimensional ordered subset expectation maximization algorithm was used to reconstruct the PET images. These were then fused with corresponding CT images for attenuation correction and anatomical referencing. Tumor and organs of interest were contoured for region-of-interest analysis of the PET images to determine the magnitude and kinetics of ^86^Y- NM600 uptake, which is reported as percent injected activity per gram of tissue (%IA/g; mean ± SEM). An *ex vivo* biodistribution study was carried out after the last scan time point.

### SPECT/CT imaging

Animals with subcutaneous MC38 (C57BL/6) tumors were administered 18.5 MBq of ^177^Lu-NM600 in the lateral tail vein. Individual mice (n=4) were placed prone into a MILabs U- SPECT6 /CTUhr system (Houten, The Netherlands) under 2% isoflurane for longitudinal scans at 3, 24, 96, and 180 h post-injection. CT scans (5 min) were acquired for anatomical reference and attenuation correction and fused with the SPECT scans (45 min). Image reconstruction used a similarity-regulated ordered-subset expectation maximization (SROSEM) algorithm. Images were quantitatively analyzed by drawing volumes of interest (VOI) over the tumor and organs of interest to determine the percent injected activity (IA) per gram (%IA/g) for each tissue. An *ex vivo* biodistribution study was carried out after the last scan time point.

### *Ex vivo* biodistribution

Mice bearing MC38 subcutaneous tumors were injected with 9.25 MBq ^86^Y-NM600, 18.5 MBq ^177^Lu-NM600, or 9.25 kBq ^225^Ac-NM600, then euthanized via CO_2_ asphyxiation at 72h (^86^Y-NM600), 180h (^177^Lu-NM600), or 2, 48, and 168 h (n = 3/timepoint) post-injection, and organs of interest were collected for *ex vivo* biodistribution for dosimetry. For comparison between low A_s_ and high A_s_ ^225^Ac-NM600 biodistributions, MC38 tumor bearing mice were injected with 9.25 kBq of low A_s_ (81.1 μg NM600/MBq ^225^Ac; n=3) or high A_s_ (2.70 μg NM600/MBq ^225^Ac; n=2) ^225^Ac-NM600, and organs of interest were collected 12 days after injection. Organs from animals that received ^225^Ac-NM600 were allowed to decay at 4°C overnight to reach secular equilibrium with ^213^Bi, which could be quantified. Organs were wet weighed, analyzed with a Perkin Elmer Wizard2 Gamma Counter (Westham, MA), and decay corrected to calculate the %IA/g for each tissue.

### *In vivo* dosimetry estimation

To estimate dosimetry of ^225^Ac-NM600, *ex vivo* biodistribution results were used. Allometric scaling was first performed to estimate the organ mass with respect to body mass, assuming 20 g for C57BL/6. Then, the residence time in each organ, MBq-sec/MBq(inj), was determined using trapezoidal integration, assuming only physical decay after the last timepoint. Lastly, the residence time for each organ was multiplied by the dose factor, mGy/MBq-sec, for each organ, which was assumed to be a sphere of equal mass that only receives self-dose. The total dose for each organ was computed by summing all contributions from the complete decay chain of ^225^Ac, considering negligible redistribution of daughter isotopes. SPECT/CT-based ^177^Lu-NM600 dosimetry was estimated according to a previously described method^29,31^. Total injected activity within each organ was calculated from extrapolation of %IA/g at a given time point. The standard mouse model was used to convert organ-specific cumulative activity into absorbed dose per injected activity (Gy/MBq). Dose contributions from surrounding organs were also included in calculations.

### Toxicity assessments

To evaluate ^225^Ac-NM600 toxicity, comprehensive metabolic panel (CMP) and complete blood count (CBC) studies were performed in control and therapy mice. Groups of naïve C57BL/6 mice (n=21/group) were administered 18.5 kBq (MTA) high A_s_ (2.70 μg NM600/MBq ^225^Ac) ^225^Ac-NM600 or 20 ng cold NM600 intravenously. Weights were recorded for n=6 mice on days 0, 7, 14, 22, 29, 36, 43, and 6 months post-injection. Three mice were culled from each cohort on days 0 (pre-injection), 7, 22, 28, 43, and 6 months post-injection, and 500 μL of blood was collected via axillary bleed. CBC analysis was completed using whole blood and an Abaxis VetScan HM5 hematology analyzer (Union City, CA). The remaining blood was centrifuged at 4000 rpm for 10 minutes to separate the serum from red blood cells; the serum was then run on an Abaxis VetScan VS2 analyzer (Union City, CA). Additionally, n=3 mice were administered 18.5 kBq low As (81.1 μg NM600/MBq ^225^Ac) ^225^Ac-NM600 intravenously. Weights were recorded for n=3 mice on days 0, 14, 41, and 6 months post-injection. Six months post injection, CBC and CMP analyses were performed by the same method as the other treatment groups.

### Radiation therapy (RT)

#### In vitro

Delivery of EBRT (195 kV) *in vitro* was performed using a RS225 Cell Irradiator (Xstrahl). Delivery of in vitro radionuclide therapy was performed as previously described. Briefly, a stock solution of ^225^Ac (unconjugated radionuclide)-containing cell culture media was prepared based on previously described dosimetry calculations^23^ to deliver either 0.25 Gy or 1 Gy absorbed dose in 24h to a cell monolayer in a six-well plate. For 0.25 Gy, 6.882 kBq ^225^Ac was added to each well; for 1 Gy, 27.75 kBq ^225^Ac was added to each well.

#### In vivo

^90^Y-, ^177^Lu-, or ^225^Ac-NM600 therapy was administered via intravenous tail vein injection on treatment day 1 for MC38 or B78 tumor-bearing mice. **Table 1** shows injected activity for each RPT dose studied.

**Table 1.** Injected RPT activity for tumor dose prescriptions.

| Tumor Model | RPT Agent | Tumor Absorbed Dose Prescription (Gy) | Injected Activity |
| --- | --- | --- | --- |
| MC38 | $^{90}\text{Y}$-NM600 | 2 | 0.925 MBq |
| | $^{177}\text{Lu}$-NM600 | 2 | 1.85 MBq |
| | $^{225}\text{Ac}$-NM600 | 0.2 | 0.4625 kBq |
| | $^{225}\text{Ac}$-NM600 | 2 | 4.625 kBq |
| | $^{225}\text{Ac}$-NM600 | 8 | 18.5 kBq |
| B78 | $^{90}\text{Y}$-NM600 | 2 | 0.733 MBq |
| | $^{177}\text{Lu}$-NM600 | 2 | 1.702 MBq |
| | $^{225}\text{Ac}$-NM600 | 0.2 | 1.85 kBq |
| | $^{225}\text{Ac}$-NM600 | 2 | 18.5 kBq |

Delivery of EBRT (300 kV) *in vivo* was performed using an X-ray biological cabinet irradiator X-RAD CIX3 (Xstrahl). The dose rate for EBRT delivery in all experiments was approximately 2 Gy/min. Dosimetric calibration and monthly quality assurance checks were performed on these irradiators by University of Wisconsin Medical Physics staff. Tumor-targeted EBRT was delivered at a dose of 2 Gy on day 1 with the right flank tumor exposed and the rest of the animal shielded using custom lead blocks.

### Clonogenic assays

*In vitro* clonogenic assay of B78 and MC38 cells was performed as previously described^25^. Briefly cells were plated at a density of 100-4000 cells/well in a six-well plate, and irradiated with either 0, 2, 4, 6, or 8 Gy EBRT or 0.25 or 1 Gy ^225^Ac (n=6 wells/treatment group; n=12 wells/0 Gy control group). For EBRT-treated cells, cells were washed with PBS and cell culture media was exchanged immediately following radiation. For ^225^Ac-treated cells, cells were washed with PBS and radioactive cell culture media was exchanged with non-radioactive cell culture media 24h following start of radiation. Surviving colonies were stained with crystal violet to aid colony counting. Colonies containing >50 cells were scored to determine plating efficiency and the fractions of the cells surviving after each radiation dose. The log surviving fraction of control and irradiated colonies were calculated and plotted.

### *In vitro* co-culture

MC38 cells or no cells were plated in 24 well plates (20,000 cells per well) containing DMEM media. Fresh media was exchanged after 24h of culture and peripheral blood mononuclear cells (PBMCs) harvested from treated or naïve mice were added. Blood (200 µL) was collected by submandibular vein collection. 50 µL of whole blood was used for CBC analysis. The remaining ∼150 µL whole blood treated with RBC lysis buffer (BioLegend) and washed with phosphate-buffered saline prior to co-culture. 200,000 PBMCs were added per well, for a 10:1 effector to target ratio. After 48 hours of co-culture, PBMCs were harvested for analysis of activation markers using flow cytometry. Number of mice/group is indicated in figure legend.

### Flow cytometry

Flow cytometry was performed as previously described^56^, using fluorescent beads (UltraComp Beads eBeads, 176 Invitrogen) to determine compensation, and fluorescence minus one (FMO) methodology to determine gating. Rainbow beads (Spherotech) were used to set voltages for each flow cytometry timepoint. For *in vivo* analysis, tumors were harvested and gently dissociated. Spleens and tumor draining lymph nodes were harvested and manually dissociated. Blood was collected by submandibular vein collection. Spleens and blood were treated with RBC lysis buffer (BioLegend) and washed with phosphate-buffered saline prior to staining. Cells were treated with CD16/32 antibody (BioLegend) to prevent non-specific binding. Live cell staining was performed using Ghost Red Dye 780 (Tonbo Biosciences) according to manufacturer’s instructions. After live-dead staining, a single cell suspension was labeled with the surface antibodies at 4°C for 30 min and washed three times using flow buffer (2% FBS + 2 mM EDTA in PBS). For intracellular staining, the cells were fixed and stained for internal markers with permeabilization solution according to manufacturer’s instructions (BD Cytofix/Cytoperm^TM^). Flow cytometry was performed using an Attune NxT Flow Cytometer (ThermoFisher). Data were analyzed using FlowJo Software. A complete list of antibody targets, clones, and fluorophores is provided in **Table S45**. The number of animals per group is indicated in figure legends.

### Immunohistochemistry

For immunohistochemistry (IHC) sections, anti-CD20 (mAb Rabbit IgG; clone E3N7O; Cell Signaling Technology 70168S, 1:200). Standard IHC methods were performed as previously described^25,57^. All labeling was performed with no primary antibody negative controls. CD20^+^ cell clusters were counted for an entire tumor cross-section, with groupings of ≥5 cells being considered a cell cluster. N=3 (^90^Y-NM600+ICI, ^225^Ac-NM600+ICI, EBRT+ICI, ICI) or n=4 (^177^Lu-NM600+ICI) tumors were quantified for positive labeling by a blinded observer.

### Single-cell RNA sequencing

#### Tumor harvest and dissociation

Tumors were dissected from three mice/treatment group, pooled by treatment, minced, and digested using the Miltenyi Biotech tumor dissociation kit (mouse, tough tumor dissociation protocol) for 40 min at 37 °C. Cells were then strained through a 70 µm filter and washed with RPMI-1640 (Corning), and resuspended in 6 mL of RPMI-1640. Tumor infiltrating lymphocytes were isolated by Ficoll-Paque density gradient centrifugation as previously described^58^. Briefly, 4 mL of Ficoll-Paque (Cytiva) was added to a 15 mL tube, and the entire cell suspension was carefully layered on top. The samples were centrifuged at 600 rcf for 20 minutes with no brake or accelerator. Cells were collected from the layer interface, rinsed with RPMI-1640 at 375 rcf, strained through a 40 µm filter, and counted on a Countess 3 automated cell counter (Invitrogen). All samples had viability > 90% and were resuspended at a concentration of 1.2 *10^6^ cells/mL in RPMI-1640 for single-cell RNA sequencing.

#### Library preparation and sequencing

scRNA-seq was performed as previously described^59^. Briefly, single-cell suspensions from isolated CD45^+^ cells were loaded on a Chromium iX single cell instrument (10x Genomics) to generate single-cell beads in emulsion and scRNA-seq libraries were prepared. For Gel bead in EMulsion (GEM) generation and library preparation, the following materials from 10x Genomics were used: Chromium Next GEM Single Cell 5′ Kit v2 (PN-1000263); VDJ Amplification, Mouse TCR Kit (PN-1000254); 5’ Feature Barcode Kit (PN-1000256); Library Construction Kit (PN-1000190); Chromium Next GEM Chip K Single Cell Kit (PN-1000286); Dual Index Kit TT Set A (PN-1000215); Dual Index Kit TN Set A (PN-1000254). The Chromium Next GEM Single Cell 5′ v2 User Guide was followed. Single-cell barcoded cDNA libraries were quantified using a Qubit Fluorometer with Qubit dsDNA HS reagent (Invitrogen), assayed using a Tapestation D1000 screentape (Agilent) and sequenced on a NovaSeq 6000 S2 flow cell (Illumina) according to 10X Genomics recommendations (26 cycles read 1, 10 cycles i7 index read, 10 cycles i5 index read, and 90 cycles read 2). Cells were sequenced to greater than 50,000 reads per cell for the 5’ gene expression library and greater than 5,000 reads per cell for the V(D)J library as recommended by manufacturer.

#### Analysis of scRNA-seq data

Cell Ranger (version 7.2.0) was used for demultiplexing, barcode processing, and gene counting. Reads were aligned to the mouse genome (version mm10). Doublets were removed by scDblFinder (version 1.14.0). We removed low-quality cells that had total read counts fewer than 1,000, had log(‘total read counts’ + 1), log(‘number of genes with read count’ + 1), or fraction of read counts in the top 20% of genes beyond five times of median absolute deviation (MAD), or had fraction of read counts from mitochondria beyond three times of MAD. Read counts were normalized across all samples by the logNormCounts function in scuttle (version 1.10.1). Dimensionality reduction was performed by Scanpy (version 1.9.1). Cell types were predicted by SingleR (version 2.2.0) using the Immunological Genome Project reference data from celldex (version 1.10.1). Gene differential expression was analyzed by MAST (version 1.26.0). Gene set enrichment analysis was performed by fgsea (version 1.26.0) using the mouse hallmark gene sets from MSigDB (version 2022.1).

#### TCR processing

For the TCR data, for each cell the cellranger multi pipeline assembles the V(D)J transcripts into contigs, aligns the contigs to the TCR reference sequence, and determines whether the contigs correspond to a CDR3 sequence by annotating both ends of the CDR3 sequences with V and J genes.

#### TCR analysis and visualization

We analyzed each sample independently and considered high confidence, productive and full length CDR3 sequences. We defined a clonotype as the unique combination of TRA and TRB CDR3 sequences annotated by different V and J genes. Then, we stratified the T cells according to their clonal expansion (**Figure 7B**). To summarize the diversity and clonal expansion at the T cell type level, we counted the number of T cells annotated with a clonotype in each cell type and then calculated the Shannon diversity, clonal expansion, and the D50 index^60,61^. We examined the T cell combinations sharing the same clonotypes per sample by using the ComplexUpset R package (v1.3.3)^62,63^.

### Statistical analysis

Prism 10 (GraphPad Software) and R 4.1.2 (R Foundation) were used for all statistical analyses. Student’s t-test was used for two-group comparisons. One-way or two-way ANOVA with Tukey’s honestly significant difference (HSD) test was used to assess statistical significance of mean differences in CBC, CMP, flow cytometry, immunohistochemistry, complete response rate, rechallenge rejection rate, and mouse body weight data. For tumor growth analysis, all available data was used. To compare treatment groups, linear mixed models after log base 10 transformation of tumor volume were fitted on cell line, time in days, and their interaction.

Tumor volumes of 0 were present in all experiments except those represented by **Figures 1L** & **3G**. In such cases, a small constant (1*10^-4^) was added prior to log transformation. Mouse ID was also included as a random intercept, accounting for correlation between measurements taken from the same mouse (**Tables S1, S4, S7, S9, S11, S14, S16, S18, S21, S23, S25, S28, S30, S32, S35, S37, S39, S42**). Pairwise contrasts were adjusted using Tukey’s method (**Tables S2, S5, S8, S10, S12, S15, S17, S19, S22, S24, S26, S29, S31, S33, S36, S38, S40, S43**). Kaplan-Meier method was used to estimate the survival distribution for the overall survival. Then, pairwise comparison of the overall survival was made using a log-rank test with Benjamini-Hochberg adjustment of p-values between levels of factors (**Tables S3, S6, S13, S20, S27, S34, S41, S44**). All data are reported as mean ± standard error of the mean (SEM) unless otherwise noted. For all graphs, *, P < 0.05; **, P < 0.01; ***, P < 0.001; and ****, P < 0.0001.

## Data availability

All data reported in this work are available within the Article, Supplementary Information, from the corresponding author upon reasonable request, or in the GEO repository (https://www.ncbi.nlm.nih.gov/geo/). Single cell RNA sequencing and T cell receptor sequencing datasets generated and analyzed during the current study are available in the GEO repository with the accession ID GSE275609 and password srgxkwkehliftcf.

## Acknowledgements

The authors’ work is supported in part by grants from NIH NCI P50CA278595, NIH NCI P01CA250972, NIH NCI P50DE026787, UWCCC Support Grant NIH NCI P30CA014520, University of Wisconsin Small Animal Imaging & Radiotherapy Facility, NIH S10OD028670-01, UWCCC Flow Cytometry Laboratory, UWCCC Cancer Informatics Shared Resource, NIH NCI F30CA268780, NIH T32GM140935, and a UW-Madison Radiology MD-PhD Graduate Student Fellowship. The authors utilized the University of Wisconsin – Madison Biotechnology Gene Expression Center (Research Resource Identifier - RRID:SCR_017757) for single cell RNA library preparation and the DNA Sequencing Facility (RRID:SCR_017759) for sequencing. The authors would like to thank the University of Wisconsin Carbone Cancer Center (UWCCC) for supporting this project. The authors would like to thank Archeus Technologies for providing NM600.

## Author contributions

CPK, AO, PAC, MP: animal work and analysis. CPK, WJJ, MT: flow cytometry staining and analysis. CAF, HCR, MBI, ANP, CFM, RH: RPT radiochemistry. CPK, AO, CC, CFM: RPT biodistribution and toxicity studies. CPK, WJJ, PAC, AKE, AGS: scRNA sequencing sample preparation. JJG, OK, BPB: PET/SPECT analysis and ^90^Y-, ^177^Lu-, ^225^Ac- NM600 dosimetry. CPK, MH: biostatistical analysis. PL, RWS, JMV, IO: bioinformatics analysis. CPK, WJJ, ZSM: experimental design. JPW and ZSM: funding, planning, and supervision of the project and data interpretation. CPK and ZSM wrote the manuscript. All authors read and approved the final manuscript.

## Competing interests

JJG is cofounder and Chief Innovation Officer of Voximetry, Inc. HCR provides consulting services to Archeus Technologies, which holds the license rights to NM600 related technologies. RH is a member of the scientific advisory board for Archeus Technologies. BPB is cofounder and Chief Science Officer of Voximetry, Inc. JPW is a founder and Chief Science Advisor for Archeus Technologies. ZSM has served as a member of the scientific advisory board for Seneca Therapeutics, Archeus Technologies, NorthStar Medical Radioisotopes, and Cali Biomedical, as a consultant for Johnson & Johnson and Telix Pharmaceuticals, and has sponsored research agreements with Point Biopharmaceuticals and Telix Pharmaceuticals. ZSM has received material support for research (drug reagents) from Bayer Pharmaceuticals, BMS, XRD therapeutics, Seneca Therapeutics, AstraZeneca, HiberCell, Apeiron, Nektar Therapeutics, and Invenra. RH, JPW, and ZSM are inventors on patents held by the University of Wisconsin Alumni Research Foundation related to select radiopharmaceutical therapies and the interaction of radiopharmaceutical therapies with immunotherapies. All other authors declare they have no competing interests.

---

**S1.**
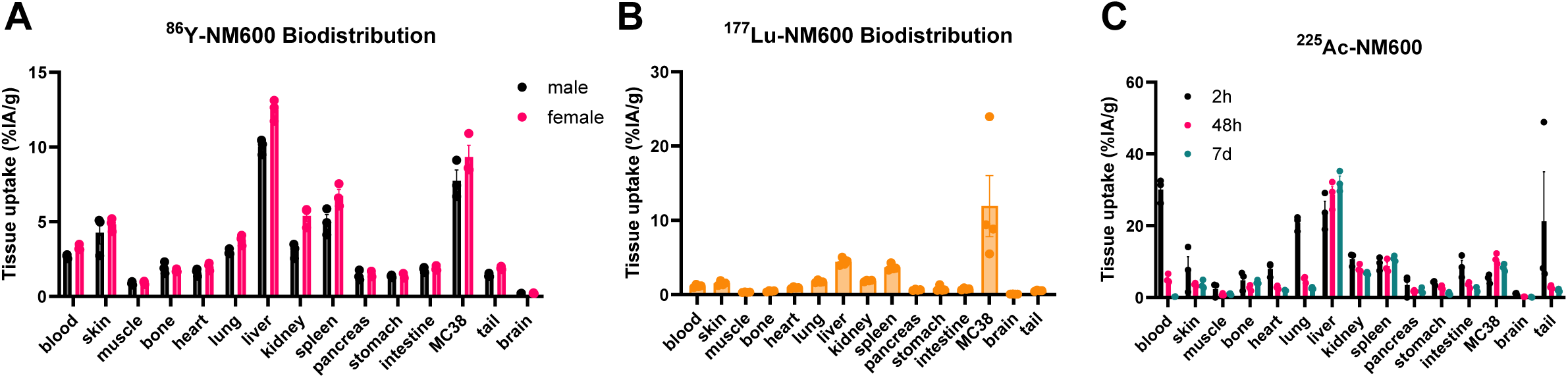
^86^Y-, ^177^Lu-, ^225^Ac-NM600 biodistribution in MC38 tumor-bearing mice

**S2.**
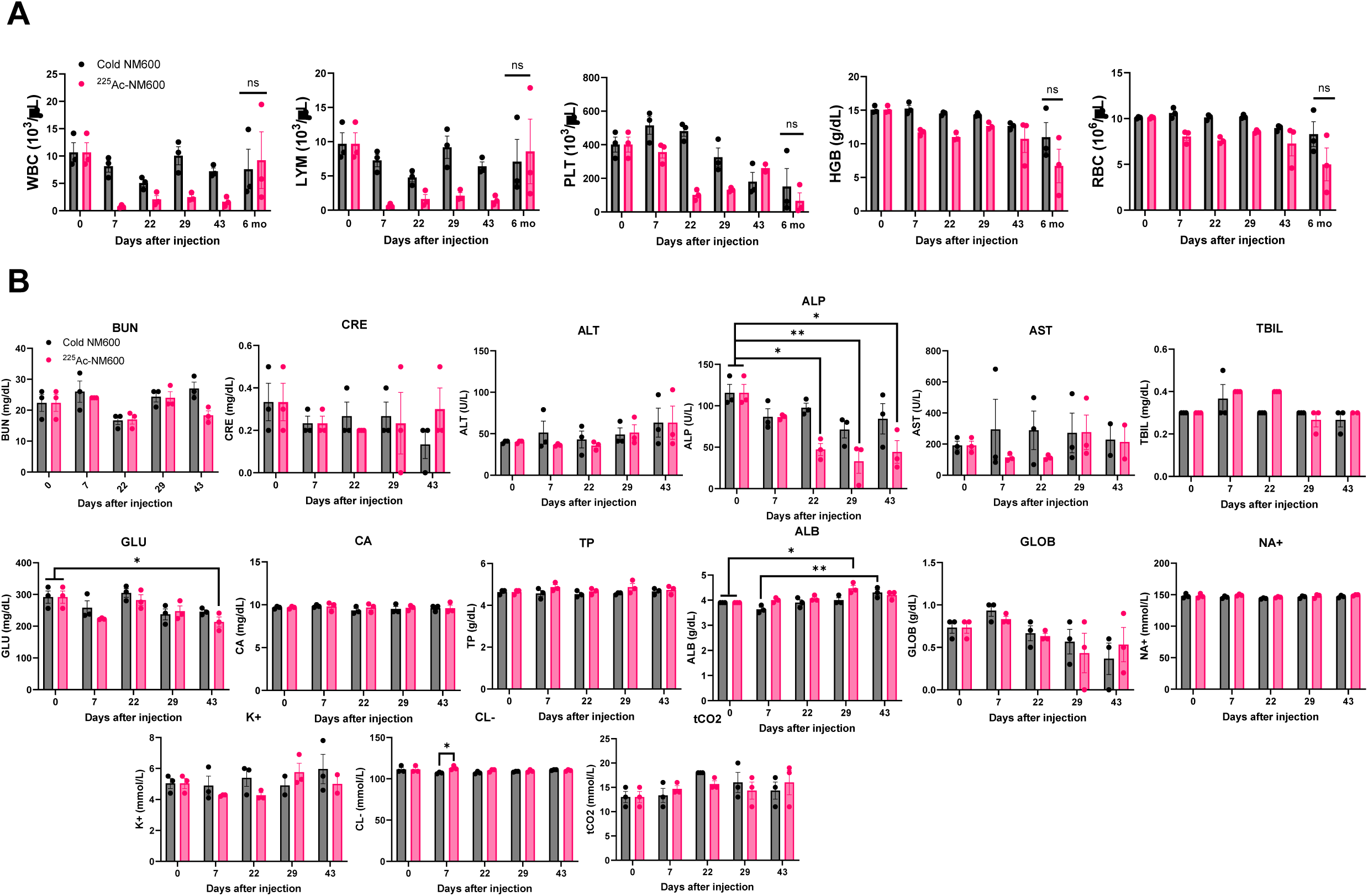
Acute toxicity profile of ^225^Ac-NM600

**S3.**
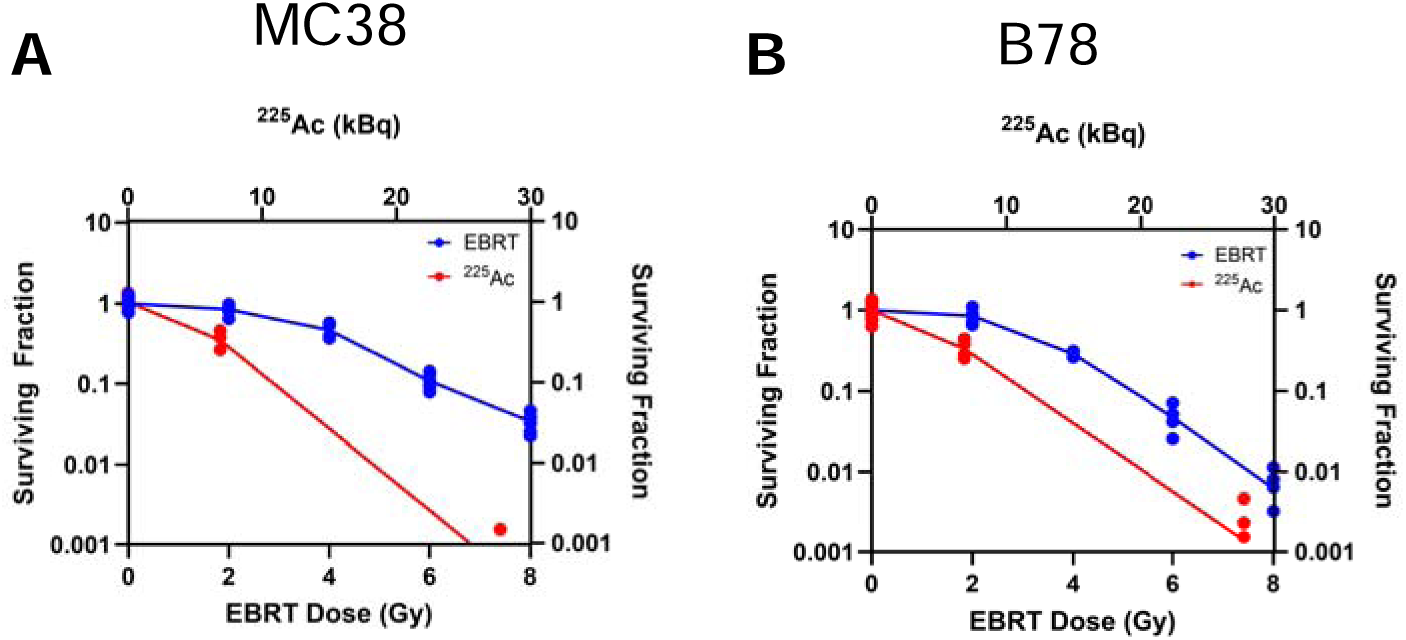
Clonogenic assays of MC38 and B78 following EBRT and ^225^Ac

**S4.**
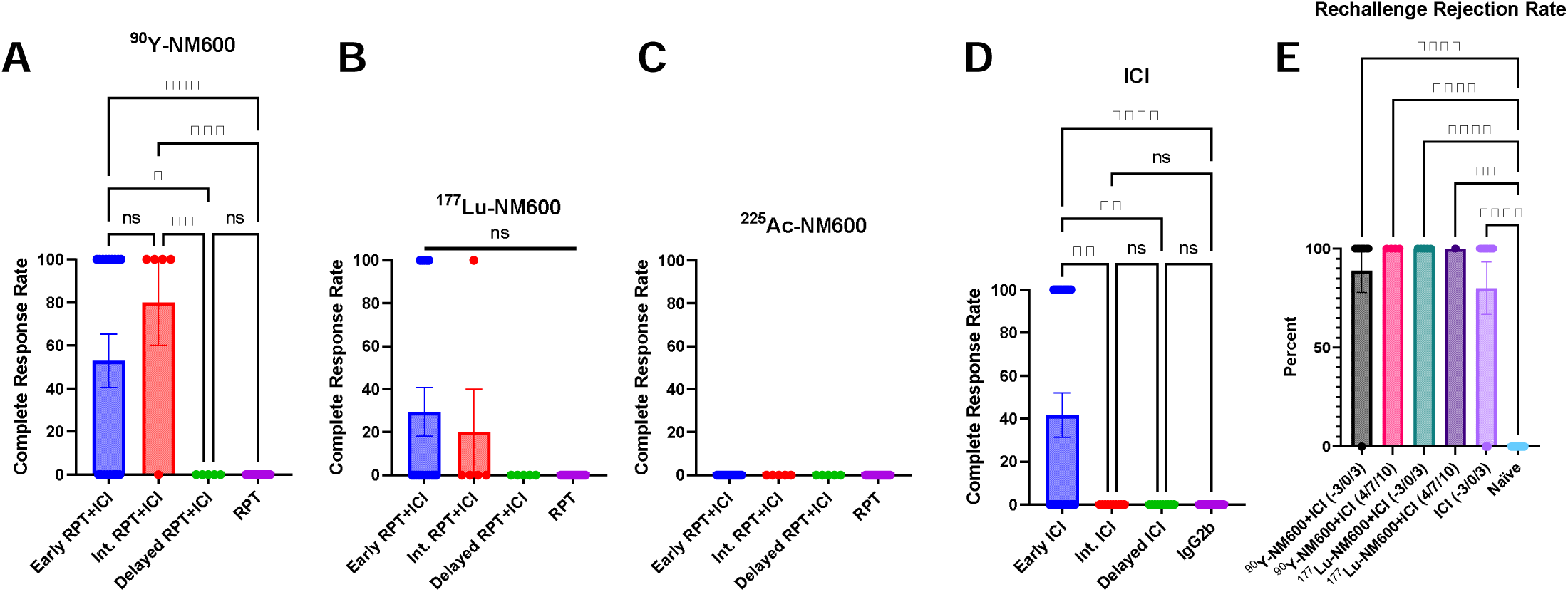
Complete response and rechallenge rejection rates for MC38 colorectal carcinoma

**S5.**
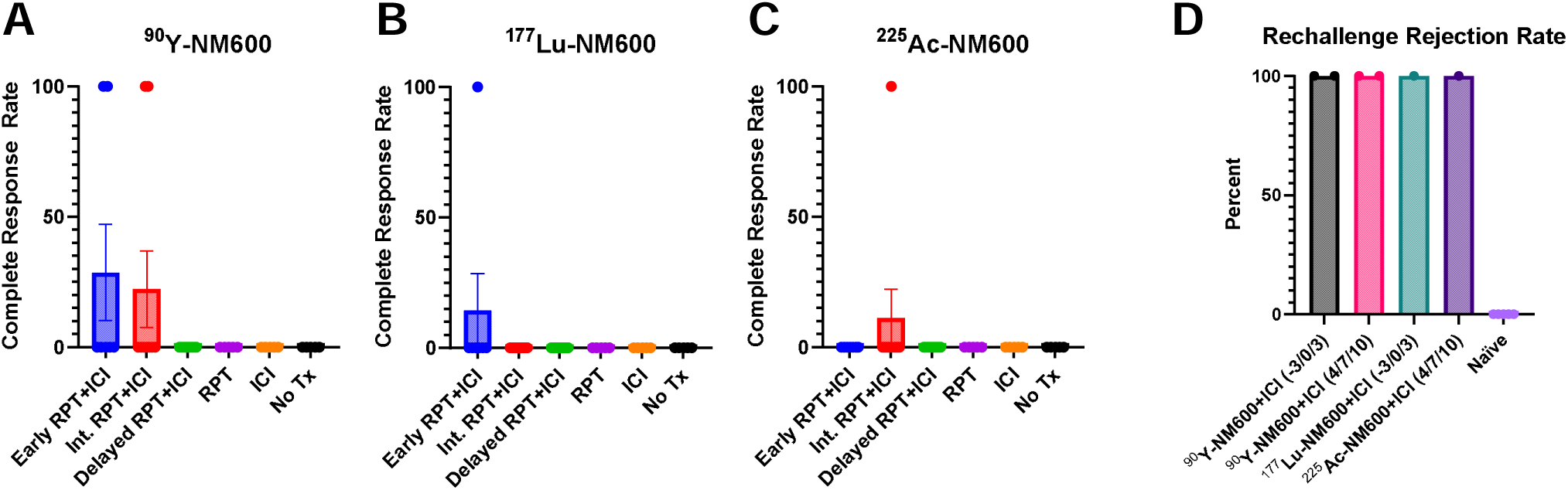
Complete response and rechallenge rejection rates for B78 melanoma

**S6.**
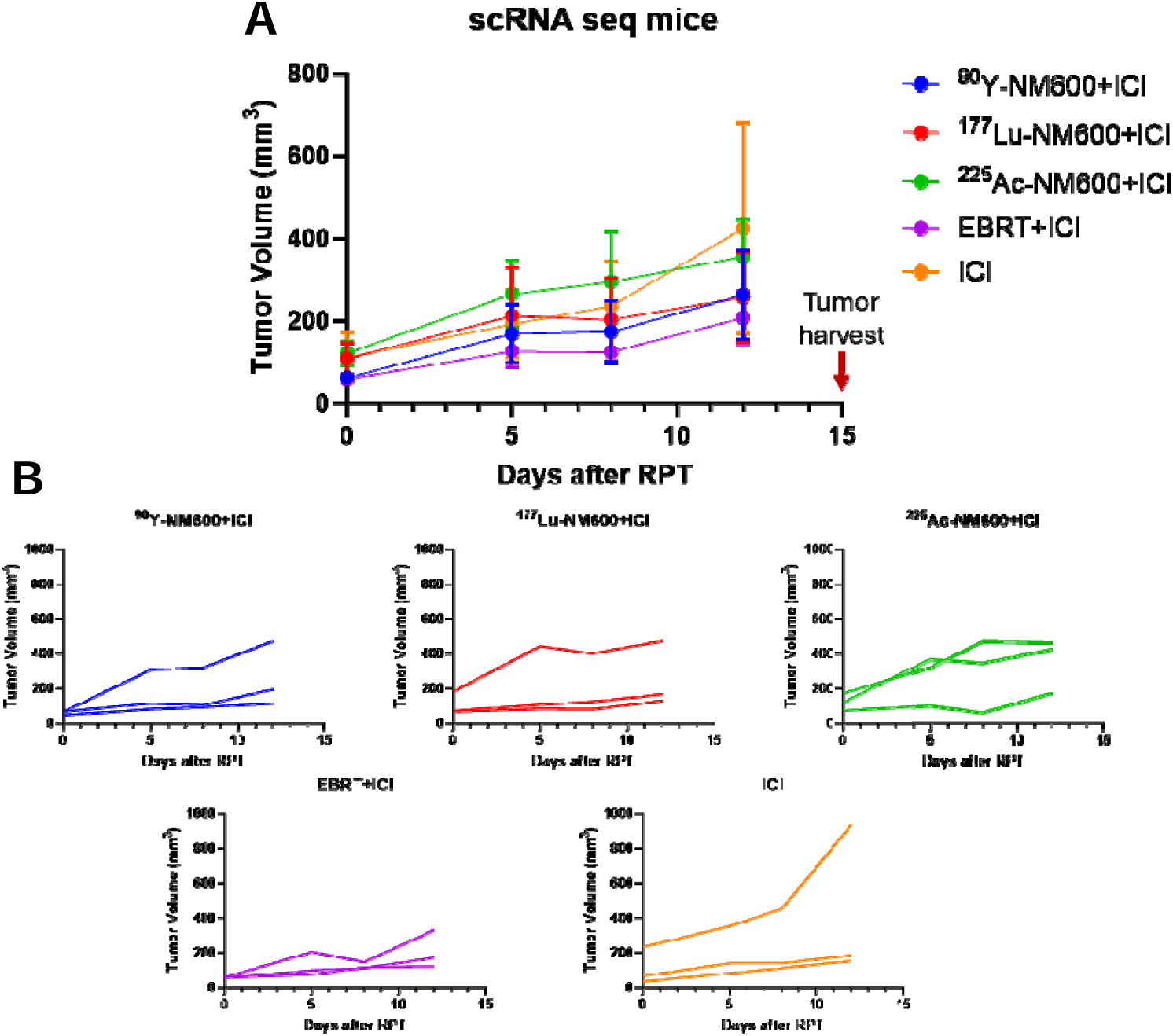
Tumor growth curves of single cell RNA sequencing experiment mice

**S7.**
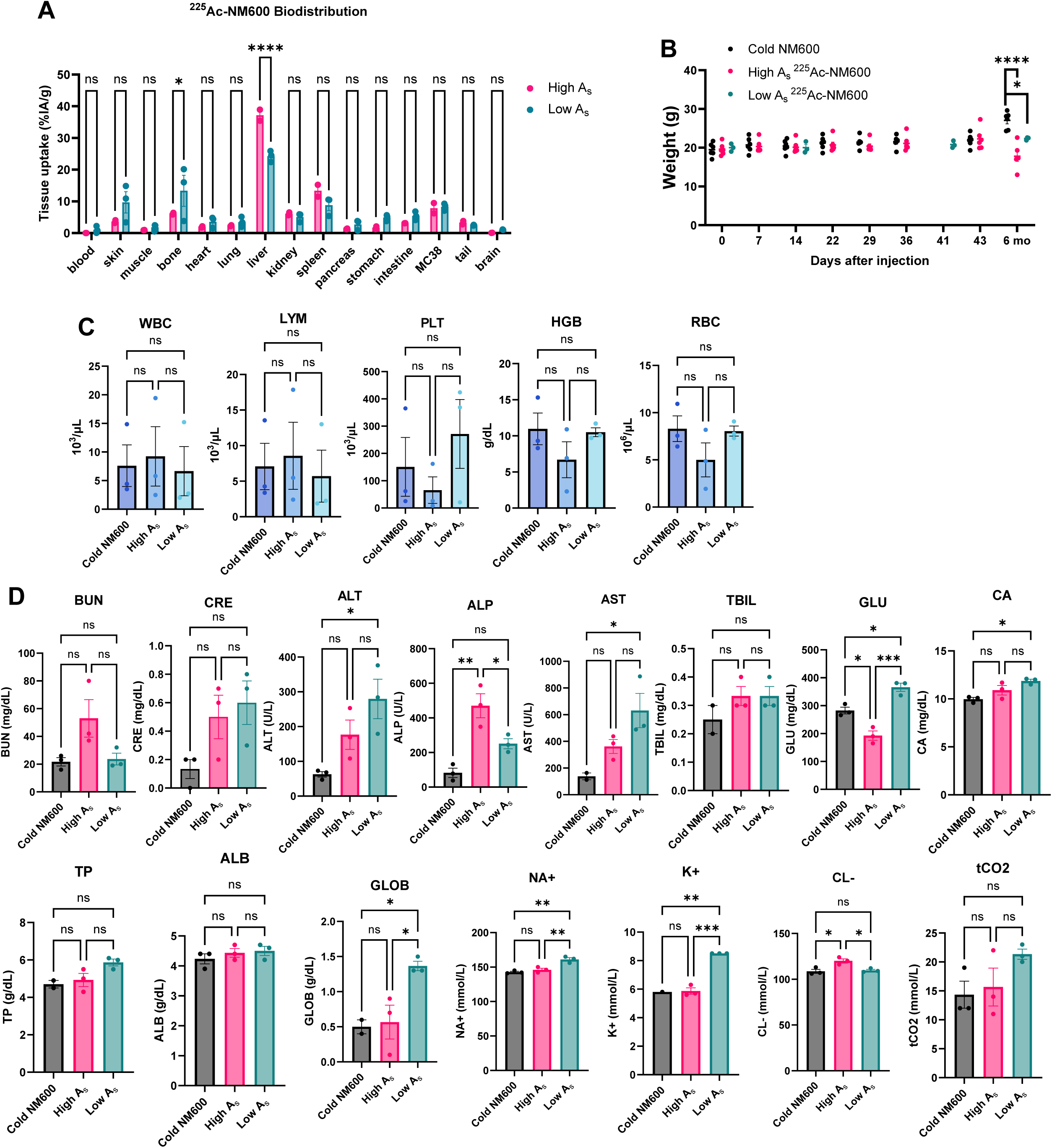
Low vs. high specific activity ^225^Ac-NM600 biodistribution and late (six month) toxicity profile

