## Supplemental Figures and Data for "Priming versus propagating: distinct immune effects of an alpha- versus beta-particle emitting radiopharmaceutical when combined with immune checkpoint inhibition"

**Table S1.** Linear mixed model coefficient estimates for fixed effects in **Figure 1H**

| Log_10_(vol+0.0001) ~ treat + day + treat*day + (1 \| id) | | | |
| --- | --- | --- | --- |
|  | **Estimate** | **95% CI** | **P-value** |
| Day | 0.0608 | (-0.0121, 0.1326) | 0.1048 |
| ICI | -0.5851 | (-1.8118, 0.6279) | 0.3548 |
| 0.2 Gy | -0.0851 | (-1.3932, 1.2075) | 0.8995 |
| 0.2 Gy+ICI | -0.1194 | (-1.3445, 1.0846) | 0.8495 |
| 2 Gy | 0.2150 | (-1.0510, 1.4600) | 0.7408 |
| 2 Gy+ICI | -0.5757 | (-1.7880, 0.6210) | 0.3563 |
| 8 Gy | -0.2159 | (-1.6102, 1.1595) | 0.7633 |
| 8 Gy+ICI | -0.3328 | (-1.5461, 0.8666) | 0.5945 |
| Day : ICI | -0.1182 | (-0.2048, -0.0305) | 0.0091 |
| Day : 0.2 Gy | 0.0068 | (-0.0996, 0.1139) | 0.9020 |
| Day : 0.2 Gy+ICI | -0.1222 | (-0.2065, -0.0372) | 0.0055 |
| Day : 2 Gy | -0.0240 | (-0.1181, 0.0716) | 0.6246 |
| Day : 2 Gy+ICI | -0.0553 | (-0.1390, 0.0294) | 0.2054 |
| Day : 8 Gy | -0.0445 | (-0.1356, 0.0476) | 0.3478 |
| Day : 8 Gy+ICI | -0.1017 | (-0.1850, -0.0172) | 0.0196 |

**Table S2.** Tukey-adjusted pairwise comparisons of estimated marginal means in **Figure 1H**

| Log_10_(vol+0.0001) ~ treat + day + treat*day + (1 \| id) | | | | | |
| --- | --- | --- | --- | --- | --- |
| **contrast** | **estimate** | **SE** | **df** | **t.ratio** | **p.value** |
| (IgG2b) - (ICI) | 0.1182 | 0.0451 | 431 | 2.619 | 0.1518 |
| (IgG2b) - (0.2 Gy) | -0.0068 | 0.0553 | 431 | -0.123 | 0.9999 |
| (IgG2b) - (0.2 Gy+ICI) | 0.1223 | 0.0438 | 431 | 2.788 | 0.1006 |
| (IgG2b) - (2 Gy) | 0.0240 | 0.0491 | 432 | 0.489 | 0.9997 |
| (IgG2b) - (2 Gy+ICI) | 0.0553 | 0.0436 | 431 | 1.268 | 0.9103 |
| (IgG2b) - (8 Gy) | 0.0446 | 0.0474 | 431 | 0.939 | 0.9820 |
| (IgG2b) - (8 Gy+ICI) | 0.1017 | 0.0434 | 431 | 2.341 | 0.2741 |
| (ICI) - (0.2 Gy) | -0.1250 | 0.0480 | 431 | -2.604 | 0.1571 |
| (ICI) - (0.2 Gy+ICI) | 0.0041 | 0.0340 | 430 | 0.119 | 0.9999 |
| (ICI) - (2 Gy) | -0.0942 | 0.0406 | 432 | -2.320 | 0.2852 |
| (ICI) - (2 Gy+ICI) | -0.0629 | 0.0337 | 430 | -1.865 | 0.5757 |
| (ICI) - (8 Gy) | -0.0737 | 0.0385 | 430 | -1.912 | 0.5431 |
| (ICI) - (8 Gy+ICI) | -0.0165 | 0.0335 | 430 | -0.494 | 0.9997 |
| (0.2 Gy) - (0.2 Gy+ICI) | 0.1291 | 0.0467 | 431 | 2.761 | 0.1078 |
| (0.2 Gy) - (2 Gy) | 0.0309 | 0.0516 | 431 | 0.598 | 0.9989 |
| (0.2 Gy) - (2 Gy+ICI) | 0.0621 | 0.0465 | 431 | 1.335 | 0.8851 |
| (0.2 Gy) - (8 Gy) | 0.0514 | 0.0501 | 431 | 1.025 | 0.9705 |
| (0.2 Gy) - (8 Gy+ICI) | 0.1085 | 0.0463 | 431 | 2.341 | 0.2738 |
| (0.2 Gy+ICI) - (2 Gy) | -0.0982 | 0.0392 | 432 | -2.507 | 0.1951 |
| (0.2 Gy+ICI) - (2 Gy+ICI) | -0.0669 | 0.0320 | 430 | -2.094 | 0.4208 |
| (0.2 Gy+ICI) - (8 Gy) | -0.0777 | 0.0370 | 430 | -2.100 | 0.4163 |
| (0.2 Gy+ICI) - (8 Gy+ICI) | -0.0206 | 0.0317 | 430 | -0.649 | 0.9981 |
| (2 Gy) - (2 Gy+ICI) | 0.0313 | 0.0389 | 431 | 0.804 | 0.9929 |
| (2 Gy) - (8 Gy) | 0.0205 | 0.0431 | 431 | 0.476 | 0.9998 |
| (2 Gy) - (8 Gy+ICI) | 0.0776 | 0.0387 | 431 | 2.008 | 0.4775 |
| (2 Gy+ICI) - (8 Gy) | -0.0108 | 0.0367 | 430 | -0.293 | 0.9999 |
| (2 Gy+ICI) - (8 Gy+ICI) | 0.0464 | 0.0314 | 430 | 1.477 | 0.8196 |
| (8 Gy) - (8 Gy+ICI) | 0.0571 | 0.0365 | 430 | 1.566 | 0.7706 |

**Table S3.** Log-rank test of MC38 overall survival (^225^Ac-NM600 dose) in **Figure 1J**

| **contrast** | **p.value** |
| --- | --- |
| (IgG2b) - (ICI) | 0.0006 *** |
| (IgG2b) - (0.2 Gy) | 0.6551 |
| (IgG2b) - (0.2 Gy+ICI) | 0.0010 *** |
| (IgG2b) - (2 Gy) | 0.0007 *** |
| (IgG2b) - (2 Gy+ICI) | <0.0001 **** |
| (IgG2b) - (8 Gy) | <0.0001 **** |
| (IgG2b) - (8 Gy+ICI) | <0.0001 **** |
| (ICI) - (0.2 Gy) | 0.0380 * |
| (ICI) - (0.2 Gy+ICI) | 0.7708 |
| (ICI) - (2 Gy) | 0.0503 |
| (ICI) - (2 Gy+ICI) | 0.8680 |
| (ICI) - (8 Gy) | 0.9210 |
| (ICI) - (8 Gy+ICI) | 0.0444 * |
| (0.2 Gy) - (0.2 Gy+ICI) | 0.0201 * |
| (0.2 Gy) - (2 Gy) | 0.3540 |
| (0.2 Gy) - (2 Gy+ICI) | 0.0008 *** |
| (0.2 Gy) - (8 Gy) | <0.0001 **** |
| (0.2 Gy) - (8 Gy+ICI) | <0.0001 **** |
| (0.2 Gy+ICI) - (2 Gy) | 0.0307 * |
| (0.2 Gy+ICI) - (2 Gy+ICI) | 0.2818 |
| (0.2 Gy+ICI) - (8 Gy) | 0.3437 |
| (0.2 Gy+ICI) - (8 Gy+ICI) | 0.1488 |
| (2 Gy) - (2 Gy+ICI) | <0.0001 **** |
| (2 Gy) - (8 Gy) | <0.0001 **** |
| (2 Gy) - (8 Gy+ICI) | <0.0001 **** |
| (2 Gy+ICI) - (8 Gy) | 0.9020 |
| (2 Gy+ICI) - (8 Gy+ICI) | 0.0006 *** |
| (8 Gy) - (8 Gy+ICI) | 0.0015 ** |

**Table S4.** Linear mixed model coefficient estimates for fixed effects in **Figure 1L**

| Log_10_(vol) ~ treat + day + treat*day + (1 \| id) | | | |
| --- | --- | --- | --- |
|  | **Estimate** | **95% CI** | **P-value** |
| Day | 0.0151 | (0.0127, 0.0176) | <.0001 |
| 2 Gy | 0.0776 | (-0.0369, 0.1924) | 0.1873 |
| 0.2 Gy+ICI | 0.3173 | (0.2009, 0.4341) | <.0001 |
| 0.2 Gy | 0.2045 | (0.0882, 0.3213) | 0.0007 |
| ICI | 0.3568 | (0.2413, 0.4742) | <.0001 |
| No Tx | 0.2416 | (0.1199, 0.3665) | 0.0001 |
| Day : 2 Gy | 0.0030 | (-0.0006, 0.0067) | 0.1024 |
| Day : 0.2 Gy+ICI | 0.0056 | (0.0017, 0.0095) | 0.0049 |
| Day : 0.2 Gy | 0.0086 | (0.0048, 0.0125) | <.0001 |
| Day : ICI | 0.0052 | (0.0014, 0.0090) | 0.0085 |
| Day : No Tx | 0.0159 | (0.0111, 0.0205) | <.0001 |

**Table S5.** Tukey-adjusted pairwise comparisons of estimated marginal means in **Figure 1L**

| Log_10_(vol) ~ treat + day + treat*day + (1 \| id) | | | | | |
| --- | --- | --- | --- | --- | --- |
| **contrast** | **estimate** | **SE** | **df** | **t.ratio** | **p.value** |
| (2 Gy+ICI) - (2 Gy) | -0.0030 | 0.0019 | 791 | -1.635 | 0.5755 |
| (2 Gy+ICI) - (0.2 Gy+ICI) | -0.0056 | 0.0020 | 791 | -2.823 | 0.0549 |
| (2 Gy+ICI) - (0.2 Gy) | -0.0086 | 0.0020 | 791 | -4.332 | 0.0002 |
| (2 Gy+ICI) - (ICI) | -0.0052 | 0.0020 | 793 | -2.637 | 0.0897 |
| (2 Gy+ICI) - (No Tx) | -0.0159 | 0.0024 | 795 | -6.622 | <.0001 |
| (2 Gy) - (0.2 Gy+ICI) | -0.0026 | 0.0021 | 791 | -1.260 | 0.8064 |
| (2 Gy) - (0.2 Gy) | -0.0056 | 0.0021 | 791 | -2.730 | 0.0706 |
| (2 Gy) - (ICI) | -0.0021 | 0.0020 | 794 | -1.058 | 0.8978 |
| (2 Gy) - (No Tx) | -0.0128 | 0.0024 | 795 | -5.243 | <.0001 |
| (0.2 Gy+ICI) - (0.2 Gy) | -0.0030 | 0.0022 | 791 | -1.389 | 0.7336 |
| (0.2 Gy+ICI) - (ICI) | 0.0004 | 0.0022 | 793 | 0.204 | 0.9999 |
| (0.2 Gy+ICI) - (No Tx) | -0.0102 | 0.0026 | 795 | -4.016 | 0.0009 |
| (0.2 Gy) - (ICI) | 0.0034 | 0.0022 | 794 | 1.603 | 0.5965 |
| (0.2 Gy) - (No Tx) | -0.0072 | 0.0026 | 795 | -2.833 | 0.0533 |
| (ICI) - (No Tx) | -0.0107 | 0.0025 | 796 | -4.220 | 0.0004 |

**Table S6.** Log-rank test of B78 overall survival (^225^Ac-NM600 dose) in **Figure 1N**

| **contrast** | **p.value** |
| --- | --- |
| (2 Gy+ICI) - (2 Gy) | 0.0507 |
| (2 Gy+ICI) - (0.2 Gy+ICI) | 0.0077 ** |
| (2 Gy+ICI) - (0.2 Gy) | 0.0013 ** |
| (2 Gy+ICI) - (ICI) | 0.0034 ** |
| (2 Gy+ICI) - (No Tx) | <0.0001 **** |
| (2 Gy) - (0.2 Gy+ICI) | 0.3721 |
| (2 Gy) - (0.2 Gy) | 0.1123 |
| (2 Gy) - (ICI) | 0.2205 |
| (2 Gy) - (No Tx) | 0.0040 ** |
| (0.2 Gy+ICI) - (0.2 Gy) | 0.4415 |
| (0.2 Gy+ICI) - (ICI) | 0.8861 |
| (0.2 Gy+ICI) - (No Tx) | 0.0628 |
| (0.2 Gy) - (ICI) | 0.8607 |
| (0.2 Gy) - (No Tx) | 0.0481 * |
| (ICI) - (No Tx) | 0.0941 |

**Table S7.** Linear mixed model coefficient estimates for fixed effects in **Figure 2F** (^90^Y-NM600)

| **2F.90Y:** Log_10_(vol+0.0001) ~ treat + day + treat*day + (1 \| id) | | | |
| --- | --- | --- | --- |
|  | **Estimate** | **95% CI** | **P-value** |
| Day | -0.1565 | (-0.1809, -0.1321) | <.0001 |
| Intermediate 90Y+ICI | 0.1465 | (-0.4963, 0.7891) | 0.6593 |
| Delayed 90Y+ICI | 1.3257 | (0.6824, 1.9691) | 0.0001 |
| 90Y | 1.1562 | (0.5672, 1.7508) | 0.0002 |
| Intermediate ICI | 1.4279 | (0.7802, 2.0760) | <.0001 |
| IgG2b | 1.5612 | (0.9527, 2.1679) | <.0001 |
| Day : Intermediate 90Y+ICI | 0.0430 | (0.0053, 0.0808) | 0.0280 |
| Day : Delayed 90Y+ICI | 0.2079 | (0.1692, 0.2465) | <.0001 |
| Day : 90Y | 0.2052 | (0.1672, 0.2433) | <.0001 |
| Day : Intermediate ICI | 0.2195 | (0.1746, 0.2641) | <.0001 |
| Day : IgG2b | 0.2444 | (0.1923, 0.2964) | <.0001 |

**Table S8.** Tukey-adjusted pairwise comparisons of estimated marginal means in **Figure 2F** (^90^Y-NM600)

| **2F.90Y:** Log_10_(vol+0.0001) ~ treat + day + treat*day + (1 \| id) | | | | | |
| --- | --- | --- | --- | --- | --- |
| **contrast** | **estimate** | **SE** | **df** | **t.ratio** | **p.value** |
| (Early 90Y+ICI) - (Intermediate 90Y+ICI) | -0.0430 | 0.0195 | 361 | -2.206 | 0.2373 |
| (Early 90Y+ICI) - (Delayed 90Y+ICI) | -0.2079 | 0.0200 | 361 | -10.413 | <.0001 |
| (Early 90Y+ICI) - (90Y) | -0.2052 | 0.0196 | 361 | -10.446 | <.0001 |
| (Early 90Y+ICI) - (Intermediate ICI) | -0.2195 | 0.0231 | 361 | -9.500 | <.0001 |
| (Early 90Y+ICI) - (IgG2b) | -0.2444 | 0.0269 | 361 | -9.088 | <.0001 |
| (Intermediate 90Y+ICI) - (Delayed 90Y+ICI) | -0.1649 | 0.0215 | 361 | -7.671 | <.0001 |
| (Intermediate 90Y+ICI) - (90Y) | -0.1622 | 0.0212 | 361 | -7.651 | <.0001 |
| (Intermediate 90Y+ICI) - (Intermediate ICI) | -0.1765 | 0.0244 | 361 | -7.222 | <.0001 |
| (Intermediate 90Y+ICI) - (IgG2b) | -0.2014 | 0.0280 | 361 | -7.180 | <.0001 |
| (Delayed 90Y+ICI) - (90Y) | 0.0027 | 0.0216 | 361 | 0.125 | 0.9999 |
| (Delayed 90Y+ICI) - (Intermediate ICI) | -0.0116 | 0.0248 | 361 | -0.469 | 0.9972 |
| (Delayed 90Y+ICI) - (IgG2b) | -0.0365 | 0.0284 | 361 | -1.287 | 0.7923 |
| (90Y) - (Intermediate ICI) | -0.0143 | 0.0245 | 361 | -0.584 | 0.9921 |
| (90Y) - (IgG2b) | -0.0392 | 0.0282 | 361 | -1.392 | 0.7318 |
| (Intermediate ICI) - (IgG2b) | -0.0249 | 0.0306 | 361 | -0.813 | 0.9651 |

**Table S9.** Linear mixed model coefficient estimates for fixed effects in **Figure 2F** (^177^Lu-NM600)

| **2F.177Lu:** Log_10_(vol+0.0001) ~ treat + day + treat*day + (1 \| id) | | | |
| --- | --- | --- | --- |
|  | **Estimate** | **95% CI** | **P-value** |
| Day | -0.1003 | (-0.1417, -0.0594) | <.0001 |
| Intermediate 177Lu+ICI | 1.1178 | (0.0058, 2.2233) | 0.0532 |
| Delayed 177Lu+ICI | 0.7532 | (-0.3999, 1.8964) | 0.2073 |
| 177Lu | 0.2443 | (-0.8153, 1.3045) | 0.6575 |
| Intermediate ICI | 0.7143 | (-0.4970, 1.9190) | 0.2557 |
| IgG2b | 0.8437 | (-0.2394, 1.9156) | 0.1325 |
| Day : Intermediate 177Lu+ICI | 0.0499 | (-0.0135, 0.1139) | 0.1321 |
| Day : Delayed 177Lu+ICI | 0.1414 | (0.0670, 0.2169) | 0.0003 |
| Day : 177Lu | 0.1822 | (0.1072, 0.2570) | <.0001 |
| Day : Intermediate ICI | 0.1604 | (0.0712, 0.2502) | 0.0006 |
| Day : IgG2b | 0.1568 | (0.0807, 0.2332) | 0.0001 |

**Table S10.** Tukey-adjusted pairwise comparisons of estimated marginal means in **Figure 2F** (^177^Lu-NM600)

| **2F.177Lu:** Log_10_(vol+0.0001) ~ treat + day + treat*day + (1 \| id) | | | | | |
| --- | --- | --- | --- | --- | --- |
| **contrast** | **estimate** | **SE** | **df** | **t.ratio** | **p.value** |
| (Early 177Lu+ICI) - (Intermediate 177Lu+ICI) | -0.0499 | 0.0330 | 257 | -1.510 | 0.6581 |
| (Early 177Lu+ICI) - (Delayed 177Lu+ICI) | -0.1414 | 0.0389 | 258 | -3.634 | 0.0045 |
| (Early 177Lu+ICI) - (177Lu) | -0.1822 | 0.0389 | 258 | -4.684 | 0.0001 |
| (Early 177Lu+ICI) - (Intermediate ICI) | -0.1604 | 0.0465 | 257 | -3.452 | 0.0085 |
| (Early 177Lu+ICI) - (IgG2b) | -0.1568 | 0.0396 | 258 | -3.958 | 0.0014 |
| (Intermediate 177Lu+ICI) - (Delayed 177Lu+ICI) | -0.0915 | 0.0412 | 258 | -2.223 | 0.2309 |
| (Intermediate 177Lu+ICI) - (177Lu) | -0.1323 | 0.0413 | 258 | -3.200 | 0.0191 |
| (Intermediate 177Lu+ICI) - (Intermediate ICI) | -0.1105 | 0.0483 | 257 | -2.285 | 0.2041 |
| (Intermediate 177Lu+ICI) - (IgG2b) | -0.1069 | 0.0419 | 258 | -2.552 | 0.1132 |
| (Delayed 177Lu+ICI) - (177Lu) | -0.0408 | 0.0461 | 258 | -0.885 | 0.9498 |
| (Delayed 177Lu+ICI) - (Intermediate ICI) | -0.0189 | 0.0525 | 257 | -0.361 | 0.9992 |
| (Delayed 177Lu+ICI) - (IgG2b) | -0.0154 | 0.0466 | 258 | -0.330 | 0.9995 |
| (177Lu) - (Intermediate ICI) | 0.0218 | 0.0526 | 258 | 0.415 | 0.9984 |
| (177Lu) - (IgG2b) | 0.0254 | 0.0463 | 257 | 0.548 | 0.9941 |
| (Intermediate ICI) - (IgG2b) | 0.0036 | 0.0530 | 257 | 0.067 | 0.9999 |

**Table S11.** Linear mixed model coefficient estimates for fixed effects in **Figure 2F** (^225^Ac-NM600)

| **2F.225Ac:** Log_10_(vol+0.0001) ~ treat + day + treat*day + (1 \| id) | | | |
| --- | --- | --- | --- |
|  | **Estimate** | **95% CI** | **P-value** |
| Day | 0.0052 | (-0.0118, 0.0222) | 0.5548 |
| Intermediate 225Ac+ICI | 0.5987 | (0.1154, 1.0863) | 0.0180 |
| Delayed 225Ac+ICI | 0.5362 | (0.0435, 1.0319) | 0.0375 |
| 225Ac | 0.7164 | (0.2550, 1.1827) | 0.0031 |
| Intermediate ICI | 0.6014 | (0.0699, 1.1353) | 0.0303 |
| IgG2b | 0.6723 | (0.1973, 1.1476) | 0.0068 |
| Day : Intermediate 225Ac+ICI | 0.0243 | (-0.0025, 0.0511) | 0.0811 |
| Day : Delayed 225Ac+ICI | 0.0376 | (0.0098, 0.0656) | 0.0098 |
| Day : 225Ac | 0.0345 | (0.0049, 0.0641) | 0.0254 |
| Day : Intermediate ICI | 0.0563 | (0.0171, 0.0955) | 0.0060 |
| Day : IgG2b | 0.0493 | (0.0161, 0.0825) | 0.0045 |

**Table S12.** Tukey-adjusted pairwise comparisons of estimated marginal means in **Figure 2F** (^225^Ac-NM600)

| **2F.225Ac:** Log_10_(vol+0.0001) ~ treat + day + treat*day + (1 \| id) | | | | | |
| --- | --- | --- | --- | --- | --- |
| **contrast** | **estimate** | **SE** | **df** | **t.ratio** | **p.value** |
| (Early 225Ac+ICI) - (Intermediate 225Ac+ICI) | -0.0243 | 0.0139 | 276 | -1.750 | 0.4999 |
| (Early 225Ac+ICI) - (Delayed 225Ac+ICI) | -0.0377 | 0.0145 | 277 | -2.599 | 0.1009 |
| (Early 225Ac+ICI) - (225Ac) | -0.0345 | 0.0154 | 277 | -2.246 | 0.2202 |
| (Early 225Ac+ICI) - (Intermediate ICI) | -0.0563 | 0.0203 | 276 | -2.770 | 0.0655 |
| (Early 225Ac+ICI) - (IgG2b) | -0.0493 | 0.0172 | 277 | -2.862 | 0.0511 |
| (Intermediate 225Ac+ICI) - (Delayed 225Ac+ICI) | -0.0133 | 0.0157 | 277 | -0.848 | 0.9580 |
| (Intermediate 225Ac+ICI) - (225Ac) | -0.0102 | 0.0165 | 277 | -0.617 | 0.9897 |
| (Intermediate 225Ac+ICI) - (Intermediate ICI) | -0.0319 | 0.0212 | 276 | -1.505 | 0.6612 |
| (Intermediate 225Ac+ICI) - (IgG2b) | -0.0250 | 0.0183 | 277 | -1.367 | 0.7469 |
| (Delayed 225Ac+ICI) - (225Ac) | 0.0031 | 0.0170 | 278 | 0.185 | 0.9999 |
| (Delayed 225Ac+ICI) - (Intermediate ICI) | -0.0186 | 0.0216 | 277 | -0.861 | 0.9553 |
| (Delayed 225Ac+ICI) - (IgG2b) | -0.0117 | 0.0187 | 278 | -0.624 | 0.9892 |
| (225Ac) - (Intermediate ICI) | -0.0217 | 0.0222 | 276 | -0.981 | 0.9236 |
| (225Ac) - (IgG2b) | -0.0148 | 0.0194 | 278 | -0.763 | 0.9734 |
| (Intermediate ICI) - (IgG2b) | 0.0069 | 0.0235 | 277 | 0.294 | 0.9997 |

**Table S13.** Log-rank test of MC38 overall survival (Varied ICI Timing) in **Figure 2H**

| **contrast** | **p.value** |
| --- | --- |
| (Early 90Y+ICI) - (Intermediate 90Y+ICI) | 0.2711 |
| (Early 90Y+ICI) - (Delayed 90Y+ICI) | 0.0016 ** |
| (Early 90Y+ICI) - (90Y) | <0.0001 **** |
| (Early 90Y+ICI) - (Intermediate ICI) | <0.0001 **** |
| (Early 90Y+ICI) - (IgG2b) | <0.0001 **** |
| (Intermediate 90Y+ICI) - (Delayed 90Y+ICI) | 0.0034 ** |
| (Intermediate 90Y+ICI) - (90Y) | 0.0006 *** |
| (Intermediate 90Y+ICI) - (Intermediate ICI) | 0.0005 *** |
| (Intermediate 90Y+ICI) - (IgG2b) | <0.0001 **** |
| (Delayed 90Y+ICI) - (90Y) | 0.1591 |
| (Delayed 90Y+ICI) - (Intermediate ICI) | 0.0772 |
| (Delayed 90Y+ICI) - (IgG2b) | 0.0006 *** |
| (90Y) - (Intermediate ICI) | 0.1125 |
| (90Y) - (IgG2b) | 0.0021 ** |
| (Early 177Lu+ICI) - (Intermediate 177Lu+ICI) | 0.8789 |
| (Early 177Lu+ICI) - (Delayed 177Lu+ICI) | 0.0231 * |
| (Early 177Lu+ICI) - (177Lu) | <0.0001 **** |
| (Early 177Lu+ICI) - (Intermediate ICI) | 0.0005 *** |
| (Early 177Lu+ICI) - (IgG2b) | <0.0001 **** |
| (Intermediate 177Lu+ICI) - (Delayed 177Lu+ICI) | 0.0423 * |
| (Intermediate 177Lu+ICI) - (177Lu) | 0.0050 ** |
| (Intermediate 177Lu+ICI) - (Intermediate ICI) | 0.0105 * |
| (Intermediate 177Lu+ICI) - (IgG2b) | 0.0003 *** |
| (Delayed 177Lu+ICI) - (177Lu) | 0.1740 |
| (Delayed 177Lu+ICI) - (Intermediate ICI) | 0.5043 |
| (Delayed 177Lu+ICI) - (IgG2b) | 0.0755 |
| (177Lu) - (Intermediate ICI) | 0.9268 |
| (177Lu) - (IgG2b) | 0.0813 |
| (Early 225Ac+ICI) - (Intermediate 225Ac+ICI) | 0.0066 ** |
| (Early 225Ac+ICI) - (Delayed 225Ac+ICI) | 0.0003 *** |
| (Early 225Ac+ICI) - (225Ac) | <0.0001 **** |
| (Early 225Ac+ICI) - (Intermediate ICI) | <0.0001 **** |
| (Early 225Ac+ICI) - (IgG2b) | <0.0001 **** |
| (Intermediate 225Ac+ICI) - (Delayed 225Ac+ICI) | 0.1969 |
| (Intermediate 225Ac+ICI) - (225Ac) | 0.1032 |
| (Intermediate 225Ac+ICI) - (Intermediate ICI) | 0.0637 |
| (Intermediate 225Ac+ICI) - (IgG2b) | 0.0001 *** |
| (Delayed 225Ac+ICI) - (225Ac) | 0.7706 |
| (Delayed 225Ac+ICI) - (Intermediate ICI) | 0.1643 |
| (Delayed 225Ac+ICI) - (IgG2b) | 0.0010 *** |
| (225Ac) - (Intermediate ICI) | 0.1771 |
| (225Ac) - (IgG2b) | <0.0001 **** |
| (Intermediate ICI) - (IgG2b) | 0.1601 |

**Table S14.** Linear mixed model coefficient estimates for fixed effects in **Figure 2G** (Early ICI)

| Log_10_(vol+0.0001) ~ treat + day + treat*day + (1 \| id) | | | |
| --- | --- | --- | --- |
|  | **Estimate** | **95% CI** | **P-value** |
| Day | -0.1565 | (-0.1926, -0.1204) | <.0001 |
| Early 177Lu+ICI | 1.1909 | (-0.5032, 2.8851) | 0.1965 |
| Early 225Ac+ICI | 1.2866 | (-0.3998, 2.9740) | 0.1629 |
| Early ICI | 1.2385 | (-0.4596, 2.9370) | 0.1809 |
| IgG2b | 1.8097 | (0.0578, 3.5616) | 0.0624 |
| Day : Early 177Lu+ICI | 0.0544 | (-0.0089, 0.1175) | 0.0959 |
| Day : Early 225Ac+ICI | 0.1623 | (0.1019, 0.2227) | <.0001 |
| Day : Early ICI | 0.1075 | (0.0420, 0.1728) | 0.0016 |
| Day : IgG2b | 0.2240 | (0.1351, 0.3126) | <.0001 |

**Table S15.** Tukey-adjusted pairwise comparisons of estimated marginal means **Figure 2G** (Early ICI)

| Log_10_(vol+0.0001) ~ treat + day + treat*day + (1 \| id) | | | | | |
| --- | --- | --- | --- | --- | --- |
| **contrast** | **estimate** | **SE** | **df** | **t.ratio** | **p.value** |
| (Early 90Y+ICI) - (Early 177Lu+ICI) | -0.0544 | 0.0326 | 304 | -1.670 | 0.4542 |
| (Early 90Y+ICI) - (Early 225Ac+ICI) | -0.1623 | 0.0311 | 304 | -5.218 | <.0001 |
| (Early 90Y+ICI) - (Early ICI) | -0.1075 | 0.0337 | 304 | -3.190 | 0.0135 |
| (Early 90Y+ICI) - (IgG2b) | -0.2240 | 0.0457 | 304 | -4.897 | <.0001 |
| (Early 177Lu+ICI) - (Early 225Ac+ICI) | -0.1079 | 0.0365 | 304 | -2.956 | 0.0276 |
| (Early 177Lu+ICI) - (Early ICI) | -0.0531 | 0.0387 | 304 | -1.373 | 0.6456 |
| (Early 177Lu+ICI) - (IgG2b) | -0.1696 | 0.0496 | 304 | -3.422 | 0.0063 |
| (Early 225Ac+ICI) - (Early ICI) | 0.0548 | 0.0376 | 304 | 1.459 | 0.5899 |
| (Early 225Ac+ICI) - (IgG2b) | -0.0617 | 0.0486 | 304 | -1.268 | 0.7110 |
| (Early ICI) - (IgG2b) | -0.1165 | 0.0503 | 304 | -2.315 | 0.1429 |

**Table S16.** Linear mixed model coefficient estimates for fixed effects **Figure 2G** (Intermediate ICI)

| Log_10_(vol+0.0001) ~ treat + day + treat*day + (1 \| id) | | | |
| --- | --- | --- | --- |
|  | **Estimate** | **95% CI** | **P-value** |
| Day | -0.1135 | (-0.1401, -0.0868) | <.0001 |
| Intermediate 177Lu+ICI | 2.0336 | (0.1021, 3.9649) | 0.0708 |
| Intermediate 225Ac+ICI | 1.6520 | (-0.2770, 3.5810) | 0.1315 |
| Intermediate ICI | 1.6988 | (-0.2637, 3.6615) | 0.1265 |
| IgG2b | 1.8520 | (-0.0706, 3.7756) | 0.0947 |
| Day : Intermediate 177Lu+ICI | 0.0673 | (0.0207, 0.1139) | 0.0057 |
| Day : Intermediate 225Ac+ICI | 0.1437 | (0.0983, 0.1891) | <.0001 |
| Day : Intermediate ICI | 0.1690 | (0.1011, 0.2371) | <.0001 |
| Day : IgG2b | 0.1605 | (0.1032, 0.2180) | <.0001 |

**Table S17.** Tukey-adjusted pairwise comparisons of estimated marginal means **Figure 2G** (Intermediate ICI)

| Log_10_(vol+0.0001) ~ treat + day + treat*day + (1 \| id) | | | | | |
| --- | --- | --- | --- | --- | --- |
| **contrast** | **estimate** | **SE** | **df** | **t.ratio** | **p.value** |
| (Intermediate 90Y+ICI) - (Intermediate 177Lu+ICI) | -0.0673 | 0.0241 | 217 | -2.794 | 0.0445 |
| (Intermediate 90Y+ICI) - (Intermediate 225Ac+ICI) | -0.1438 | 0.0235 | 217 | -6.122 | <.0001 |
| (Intermediate 90Y+ICI) - (Intermediate ICI) | -0.1690 | 0.0352 | 217 | -4.806 | <.0001 |
| (Intermediate 90Y+ICI) - (IgG2b) | -0.1606 | 0.0297 | 217 | -5.410 | <.0001 |
| (Intermediate 177Lu+ICI) - (Intermediate 225Ac+ICI) | -0.0764 | 0.0274 | 217 | -2.789 | 0.0451 |
| (Intermediate 177Lu+ICI) - (Intermediate ICI) | -0.1017 | 0.0379 | 217 | -2.681 | 0.0601 |
| (Intermediate 177Lu+ICI) - (IgG2b) | -0.0932 | 0.0329 | 217 | -2.831 | 0.0403 |
| (Intermediate 225Ac+ICI) - (Intermediate ICI) | -0.0253 | 0.0375 | 217 | -0.674 | 0.9619 |
| (Intermediate 225Ac+ICI) - (IgG2b) | -0.0168 | 0.0324 | 217 | -0.518 | 0.9855 |
| (Intermediate ICI) - (IgG2b) | 0.0085 | 0.0416 | 217 | 0.204 | 0.9996 |

**Table S18.** Linear mixed model coefficient estimates for fixed effects **Figure 2G** (Delayed ICI)

| Log_10_(vol+0.0001) ~ treat + day + treat*day + (1 \| id) | | | |
| --- | --- | --- | --- |
|  | **Estimate** | **95% CI** | **P-value** |
| Day | 0.0506 | (0.0459, 0.0554) | <.0001 |
| Delayed 177Lu+ICI | 0.4341 | (0.2904, 0.5783) | <.0001 |
| Delayed 225Ac+ICI | 0.4261 | (0.2879, 0.5643) | <.0001 |
| Delayed ICI | 0.4790 | (0.3190, 0.6395) | <.0001 |
| IgG2b | 0.4811 | (0.3478, 0.6154) | <.0001 |
| Day : Delayed 177Lu+ICI | 0.0025 | (-0.0073, 0.0121) | 0.6265 |
| Day : Delayed 225Ac+ICI | -0.0071 | (-0.0154, 0.0011) | 0.0969 |
| Day : Delayed ICI | 0.0265 | (0.0118, 0.0411) | 0.0006 |
| Day : IgG2b | 0.0060 | (-0.0040, 0.0158) | 0.2475 |

**Table S19.** Tukey-adjusted pairwise comparisons of estimated marginal means **Figure 2G** (Delayed ICI)

| Log_10_(vol+0.0001) ~ treat + day + treat*day + (1 \| id) | | | | | |
| --- | --- | --- | --- | --- | --- |
| **contrast** | **estimate** | **SE** | **df** | **t.ratio** | **p.value** |
| (Delayed 90Y+ICI) - (Delayed 177Lu+ICI) | -0.0025 | 0.0050 | 201 | -0.487 | 0.9885 |
| (Delayed 90Y+ICI) - (Delayed 225Ac+ICI) | 0.0071 | 0.0043 | 201 | 1.667 | 0.4566 |
| (Delayed 90Y+ICI) - (Delayed ICI) | -0.0265 | 0.0076 | 201 | -3.481 | 0.0055 |
| (Delayed 90Y+ICI) - (IgG2b) | -0.0060 | 0.0051 | 201 | -1.159 | 0.7744 |
| (Delayed 177Lu+ICI) - (Delayed 225Ac+ICI) | 0.0096 | 0.0056 | 201 | 1.705 | 0.4332 |
| (Delayed 177Lu+ICI) - (Delayed ICI) | -0.0240 | 0.0084 | 201 | -2.865 | 0.0368 |
| (Delayed 177Lu+ICI) - (IgG2b) | -0.0035 | 0.0063 | 201 | -0.557 | 0.9809 |
| (Delayed 225Ac+ICI) - (Delayed ICI) | -0.0336 | 0.0080 | 201 | -4.197 | 0.0004 |
| (Delayed 225Ac+ICI) - (IgG2b) | -0.0131 | 0.0057 | 201 | -2.300 | 0.1492 |
| (Delayed ICI) - (IgG2b) | 0.0205 | 0.0085 | 201 | 2.422 | 0.1136 |

**Table S20.** Log-rank test of MC38 overall survival (Varied Radionuclide) in **Figure 2I**

| **contrast** | **p.value** |
| --- | --- |
| (Early 90Y+ICI) - (Early Lu+ICI) | 0.1085 |
| (Early 90Y+ICI) - (Early 225Ac+ICI) | 0.0123 * |
| (Early 90Y+ICI) – (Early ICI) | 0.3373 |
| (Early Lu+ICI) – (Early 225Ac+ICI) | 0.2264 |
| (Early Lu+ICI) – (Early ICI) | 0.5706 |
| (Early 225Ac+ICI) – (Early ICI) | 0.0857 |
| (Early ICI) – (IgG2b) | 0.0006 *** |
| (Intermediate 90Y+ICI) - (Intermediate Lu+ICI) | 0.0612 |
| (Intermediate 90Y+ICI) - (Intermediate 225Ac+ICI) | 0.0018 ** |
| (Intermediate Lu+ICI) – (Intermediate 225Ac+ICI) | 0.4769 |
| (Delayed 90Y+ICI) - (Delayed Lu+ICI) | 0.4107 |
| (Delayed 90Y+ICI) - (Delayed 225Ac+ICI) | 0.5158 |
| (Delayed 90Y+ICI) – (Delayed ICI) | 0.0679 |
| (Delayed Lu+ICI) – (Delayed 225Ac+ICI) | 0.7332 |
| (Delayed Lu+ICI) – (Delayed ICI) | 0.3239 |
| (Delayed 225Ac+ICI) - (Delayed ICI) | 0.1338 |
| (Delayed ICI) – (IgG2b) | 0.4683 |

**Table S21.** Linear mixed model coefficient estimates for fixed effects in **Figure 3F** (^90^Y-NM600)

| Log_10_(vol+0.0001) ~ treat + day + treat*day + (1 \| id) | | | |
| --- | --- | --- | --- |
|  | **Estimate** | **95% CI** | **P-value** |
| Day | -0.0169 | (-0.0305, -0.0033) | 0.0159 |
| Intermediate 90Y+ICI | 0.6084 | (-0.0624, 1.2726) | 0.0766 |
| Delayed 90Y+ICI | 0.6529 | (-0.0130, 1.3166) | 0.0565 |
| 90Y | 0.4644 | (-0.3210, 1.2451) | 0.2491 |
| ICI | 1.1362 | (0.3289, 1.9354) | 0.0061 |
| No Tx | 0.6538 | (-0.1555, 1.4609) | 0.1161 |
| Day : Intermediate 90Y+ICI | 0.0126 | (-0.0059, 0.0312) | 0.1861 |
| Day : Delayed 90Y+ICI | 0.0390 | (0.0204, 0.0577) | 0.0001 |
| Day : 90Y | 0.0442 | (0.0225, 0.0659) | 0.0001 |
| Day : ICI | 0.0354 | (0.0116, 0.0593) | 0.0040 |
| Day : No Tx | 0.0513 | (0.0260, 0.0766) | 0.0001 |

**Table S22.** Tukey-adjusted pairwise comparisons of estimated marginal means in **Figure 3F** (^90^Y-NM600)

| Log_10_(vol+0.0001) ~ treat + day + treat*day + (1 \| id) | | | | | |
| --- | --- | --- | --- | --- | --- |
| **contrast** | **estimate** | **SE** | **df** | **t.ratio** | **p.value** |
| (Early 90Y+ICI) - (Intermediate 90Y+ICI) | -0.0126 | 0.0095 | 601 | -1.324 | 0.7718 |
| (Early 90Y+ICI) - (Delayed 90Y+ICI) | -0.0390 | 0.0096 | 601 | -4.074 | 0.0007 |
| (Early 90Y+ICI) - (90Y) | -0.0442 | 0.0112 | 601 | -3.963 | 0.0012 |
| (Early 90Y+ICI) - (ICI) | -0.0354 | 0.0123 | 602 | -2.892 | 0.0456 |
| (Early 90Y+ICI) - (No Tx) | -0.0513 | 0.0130 | 601 | -3.947 | 0.0012 |
| (Intermediate 90Y+ICI) - (Delayed 90Y+ICI) | -0.0264 | 0.0092 | 601 | -2.864 | 0.0493 |
| (Intermediate 90Y+ICI) - (90Y) | -0.0316 | 0.0109 | 601 | -2.909 | 0.0434 |
| (Intermediate 90Y+ICI) - (ICI) | -0.0228 | 0.0120 | 602 | -1.906 | 0.3991 |
| (Intermediate 90Y+ICI) - (No Tx) | -0.0387 | 0.0128 | 601 | -3.034 | 0.0302 |
| (Delayed 90Y+ICI) - (90Y) | -0.0052 | 0.0109 | 601 | -0.474 | 0.9970 |
| (Delayed 90Y+ICI) - (ICI) | 0.0036 | 0.0120 | 602 | 0.299 | 0.9997 |
| (Delayed 90Y+ICI) - (No Tx) | -0.0123 | 0.0128 | 601 | -0.960 | 0.9303 |
| (90Y) - (ICI) | 0.0088 | 0.0133 | 602 | 0.659 | 0.9863 |
| (90Y) - (No Tx) | -0.0071 | 0.0140 | 601 | -0.508 | 0.9959 |
| (ICI) - (No Tx) | -0.0159 | 0.0149 | 602 | -1.062 | 0.8961 |

**Table S23.** Linear mixed model coefficient estimates for fixed effects in **Figure 3F** (^177^Lu-NM600)

| Log_10_(vol+0.0001) ~ treat + day + treat*day + (1 \| id) | | | |
| --- | --- | --- | --- |
|  | **Estimate** | **95% CI** | **P-value** |
| Day | -0.0087 | (-0.0174, 0.0001) | 0.0543 |
| Intermediate 177Lu+ICI | 0.1648 | (-0.2627, 0.5854) | 0.4494 |
| Delayed 177Lu+ICI | 0.1205 | (-0.3057, 0.5453) | 0.5823 |
| 177Lu | -0.0088 | (-0.5124, 0.4949) | 0.9729 |
| ICI | 0.1057 | (-0.3960, 0.6127) | 0.6835 |
| No Tx | 0.1581 | (-0.3607, 0.6698) | 0.5507 |
| Day : Intermediate 177Lu+ICI | 0.0228 | (0.0111, 0.0346) | 0.0002 |
| Day : Delayed 177Lu+ICI | 0.0233 | (0.0112, 0.0352) | 0.0002 |
| Day : 177Lu | 0.0413 | (0.0258, 0.0569) | <.0001 |
| Day : ICI | 0.0282 | (0.0129, 0.0433) | 0.0003 |
| Day : No Tx | 0.0413 | (0.0251, 0.0574) | <.0001 |

**Table S24.** Tukey-adjusted pairwise comparisons of estimated marginal means in **Figure 3F** (^177^Lu-NM600)

| Log_10_(vol+0.0001) ~ treat + day + treat*day + (1 \| id) | | | | | |
| --- | --- | --- | --- | --- | --- |
| **contrast** | **estimate** | **SE** | **df** | **t.ratio** | **p.value** |
| (Early 177Lu+ICI) - (Intermediate 177Lu+ICI) | -0.0228 | 0.0060 | 588 | -3.775 | 0.0024 |
| (Early 177Lu+ICI) - (Delayed 177Lu+ICI) | -0.0233 | 0.0062 | 588 | -3.774 | 0.0024 |
| (Early 177Lu+ICI) - (177Lu) | -0.0413 | 0.0080 | 589 | -5.168 | <.0001 |
| (Early 177Lu+ICI) - (ICI) | -0.0282 | 0.0078 | 590 | -3.598 | 0.0047 |
| (Early 177Lu+ICI) - (No Tx) | -0.0413 | 0.0083 | 589 | -4.974 | <.0001 |
| (Intermediate 177Lu+ICI) - (Delayed 177Lu+ICI) | -0.0005 | 0.0058 | 588 | -0.077 | 0.9999 |
| (Intermediate 177Lu+ICI) - (177Lu) | -0.0185 | 0.0077 | 589 | -2.389 | 0.1618 |
| (Intermediate 177Lu+ICI) - (ICI) | -0.0054 | 0.0076 | 590 | -0.709 | 0.9809 |
| (Intermediate 177Lu+ICI) - (No Tx) | -0.0185 | 0.0081 | 589 | -2.292 | 0.1990 |
| (Delayed 177Lu+ICI) - (177Lu) | -0.0180 | 0.0079 | 590 | -2.298 | 0.1965 |
| (Delayed 177Lu+ICI) - (ICI) | -0.0049 | 0.0077 | 589 | -0.643 | 0.9877 |
| (Delayed 177Lu+ICI) - (No Tx) | -0.0180 | 0.0082 | 589 | -2.208 | 0.2355 |
| (177Lu) - (ICI) | 0.0131 | 0.0092 | 590 | 1.421 | 0.7141 |
| (177Lu) - (No Tx) | 0.00003 | 0.0096 | 588 | 0.003 | 0.9999 |
| (ICI) - (No Tx) | -0.0131 | 0.0095 | 591 | -1.378 | 0.7405 |

**Table S25.** Linear mixed model coefficient estimates for fixed effects in **Figure 3F** (^225^Ac-NM600)

| Log_10_(vol+0.0001) ~ treat + day + treat*day + (1 \| id) | | | |
| --- | --- | --- | --- |
|  | **Estimate** | **95% CI** | **P-value** |
| Day | -0.0136 | (-0.0205, -0.0067) | 0.0001 |
| Intermediate 225Ac+ICI | -0.0216 | (-0.3591, 0.3147) | 0.9006 |
| Delayed 225Ac+ICI | -0.0415 | (-0.3761, 0.2948) | 0.8100 |
| 225Ac | -0.2861 | (-0.6779, 0.1104) | 0.1583 |
| ICI | 0.2200 | (-0.1806, 0.6172) | 0.2834 |
| No Tx | 0.0461 | (-0.3669, 0.4562) | 0.8277 |
| Day : Intermediate 225Ac+ICI | 0.0231 | (0.0139, 0.0323) | <.0001 |
| Day : Delayed 225Ac+ICI | 0.0244 | (0.0152, 0.0336) | <.0001 |
| Day : 225Ac | 0.0325 | (0.0219, 0.0432) | <.0001 |
| Day : ICI | 0.0266 | (0.0146, 0.0388) | <.0001 |
| Day : No Tx | 0.0486 | (0.0357, 0.0614) | <.0001 |

**Table S26.** Tukey-adjusted pairwise comparisons of estimated marginal means in **Figure 3F** (^225^Ac-NM600)

| Log_10_(vol+0.0001) ~ treat + day + treat*day + (1 \| id) | | | | | |
| --- | --- | --- | --- | --- | --- |
| **contrast** | **estimate** | **SE** | **df** | **t.ratio** | **p.value** |
| (Early 225Ac+ICI) - (Intermediate 225Ac+ICI) | -0.0231 | 0.0047 | 626 | -4.888 | <.0001 |
| (Early 225Ac+ICI) - (Delayed 225Ac+ICI) | -0.0244 | 0.0047 | 626 | -5.168 | <.0001 |
| (Early 225Ac+ICI) - (225Ac) | -0.0325 | 0.0055 | 626 | -5.933 | <.0001 |
| (Early 225Ac+ICI) - (ICI) | -0.0266 | 0.0062 | 628 | -4.281 | 0.0003 |
| (Early 225Ac+ICI) - (No Tx) | -0.0486 | 0.0066 | 628 | -7.371 | <.0001 |
| (Intermediate 225Ac+ICI) - (Delayed 225Ac+ICI) | -0.0013 | 0.0044 | 626 | -0.300 | 0.9997 |
| (Intermediate 225Ac+ICI) - (225Ac) | -0.0095 | 0.0052 | 626 | -1.812 | 0.4584 |
| (Intermediate 225Ac+ICI) - (ICI) | -0.0035 | 0.0060 | 628 | -0.589 | 0.9918 |
| (Intermediate 225Ac+ICI) - (No Tx) | -0.0255 | 0.0064 | 629 | -4.001 | 0.0010 |
| (Delayed 225Ac+ICI) - (225Ac) | -0.0081 | 0.0052 | 626 | -1.559 | 0.6261 |
| (Delayed 225Ac+ICI) - (ICI) | -0.0022 | 0.0060 | 628 | -0.367 | 0.9991 |
| (Delayed 225Ac+ICI) - (No Tx) | -0.0242 | 0.0064 | 629 | -3.794 | 0.0022 |
| (225Ac) - (ICI) | 0.0060 | 0.0066 | 628 | 0.900 | 0.9464 |
| (225Ac) - (No Tx) | -0.0161 | 0.0070 | 628 | -2.305 | 0.1934 |
| (ICI) - (No Tx) | -0.0220 | 0.0076 | 630 | -2.904 | 0.0440 |

**Table S27.** Log-rank test of B78 overall survival (Varied ICI Timing) in **Figure 3H**

| **contrast** | **p.value** |
| --- | --- |
| (Early 90Y+ICI) - (Intermediate 90Y+ICI) | 0.5471 |
| (Early 90Y+ICI) - (Delayed 90Y+ICI) | 0.0234 * |
| (Early 90Y+ICI) - (90Y) | 0.0371 * |
| (Early 90Y+ICI) - (ICI) | 0.0953 |
| (Early 90Y+ICI) - (No Tx) | 0.0003 *** |
| (Intermediate 90Y+ICI) - (Delayed 90Y+ICI) | 0.0437 * |
| (Intermediate 90Y+ICI) - (90Y) | 0.0768 |
| (Intermediate 90Y+ICI) - (ICI) | 0.2531 |
| (Intermediate 90Y+ICI) - (No Tx) | 0.0024 ** |
| (Delayed 90Y+ICI) - (90Y) | 0.3923 |
| (Delayed 90Y+ICI) - (ICI) | 0.7741 |
| (Delayed 90Y+ICI) - (No Tx) | 0.0041 ** |
| (90Y) - (ICI) | 0.7360 |
| (90Y) - (No Tx) | 0.0127 * |
| (Early 177Lu+ICI) - (Intermediate 177Lu+ICI) | 0.5665 |
| (Early 177Lu+ICI) - (Delayed 177Lu+ICI) | 0.3093 |
| (Early 177Lu+ICI) - (177Lu) | 0.0010 ** |
| (Early 177Lu+ICI) - (ICI) | 0.0714 |
| (Early 177Lu+ICI) - (No Tx) | 0.0003 *** |
| (Intermediate 177Lu+ICI) - (Delayed 177Lu+ICI) | 0.6140 |
| (Intermediate 177Lu+ICI) - (177Lu) | 0.0034 ** |
| (Intermediate 177Lu+ICI) - (ICI) | 0.3171 |
| (Intermediate 177Lu+ICI) - (No Tx) | 0.0018 ** |
| (Delayed 177Lu+ICI) - (177Lu) | 0.0205 * |
| (Delayed 177Lu+ICI) - (ICI) | 0.3091 |
| (Delayed 177Lu+ICI) - (No Tx) | 0.0027 ** |
| (177Lu) - (ICI) | 0.6058 |
| (177Lu) - (No Tx) | 0.2133 |
| (Early 225Ac+ICI) - (Intermediate 225Ac+ICI) | 0.1971 |
| (Early 225Ac+ICI) - (Delayed 225Ac+ICI) | 0.4211 |
| (Early 225Ac+ICI) - (225Ac) | 0.0051 ** |
| (Early 225Ac+ICI) - (ICI) | 0.0379 * |
| (Early 225Ac+ICI) - (No Tx) | 0.0003 *** |
| (Intermediate 225Ac+ICI) - (Delayed 225Ac+ICI) | 0.8124 |
| (Intermediate 225Ac+ICI) - (225Ac) | 0.1499 |
| (Intermediate 225Ac+ICI) - (ICI) | 0.0924 |
| (Intermediate 225Ac+ICI) - (No Tx) | <0.0001 **** |
| (Delayed 225Ac+ICI) - (225Ac) | 0.1076 |
| (Delayed 225Ac+ICI) - (ICI) | 0.0684 |
| (Delayed 225Ac+ICI) - (No Tx) | <0.0001 **** |
| (225Ac) - (ICI) | 0.5700 |
| (225Ac) - (No Tx) | 0.0018 ** |
| (ICI) - (No Tx) | 0.4358 |

**Table S28.** Linear mixed model coefficient estimates for fixed effects in **Figure 3G** (Early ICI)

| Log_10_(vol+0.0001) ~ treat + day + treat*day + (1 \| id) | | | |
| --- | --- | --- | --- |
|  | **Estimate** | **95% CI** | **P-value** |
| Day | -0.0135 | (-0.0291, 0.0020) | 0.0914 |
| Early 177Lu+ICI | 0.8576 | (0.0565, 1.6511) | 0.0368 |
| Early 225Ac+ICI | 0.9408 | (0.1392, 1.7340) | 0.0221 |
| ICI | 1.5078 | (0.5913, 2.4094) | 0.0013 |
| No Tx | 0.7990 | (-0.1324, 1.7216) | 0.0943 |
| Day : Early 177Lu+ICI | 0.0054 | (-0.0166, 0.0275) | 0.6312 |
| Day : Early 225Ac+ICI | -0.0001 | (-0.0220, 0.0219) | 0.9957 |
| Day : ICI | 0.0240 | (-0.0026, 0.0507) | 0.0802 |
| Day : No Tx | 0.0512 | (0.0224, 0.0800) | 0.0006 |

**Table S29.** Tukey-adjusted pairwise comparisons of estimated marginal means in **Figure 3G** (Early ICI)

| Log_10_(vol+0.0001) ~ treat + day + treat*day + (1 \| id) | | | | | |
| --- | --- | --- | --- | --- | --- |
| **contrast** | **estimate** | **SE** | **df** | **t.ratio** | **p.value** |
| (Early 90Y+ICI) - (Early 177Lu+ICI) | -0.0054 | 0.0113 | 476 | -0.480 | 0.9891 |
| (Early 90Y+ICI) - (Early 225Ac+ICI) | 0.0001 | 0.0113 | 476 | 0.005 | 0.9999 |
| (Early 90Y+ICI) - (ICI) | -0.0240 | 0.0137 | 477 | -1.753 | 0.4028 |
| (Early 90Y+ICI) - (No Tx) | -0.0512 | 0.0148 | 477 | -3.459 | 0.0053 |
| (Early 177Lu+ICI) - (Early 225Ac+ICI) | 0.0055 | 0.0113 | 476 | 0.488 | 0.9884 |
| (Early 177Lu+ICI) - (ICI) | -0.0186 | 0.0137 | 477 | -1.356 | 0.6564 |
| (Early 177Lu+ICI) - (No Tx) | -0.0458 | 0.0148 | 477 | -3.092 | 0.0179 |
| (Early 225Ac+ICI) - (ICI) | -0.0241 | 0.0136 | 477 | -1.765 | 0.3956 |
| (Early 225Ac+ICI) - (No Tx) | -0.0513 | 0.0148 | 477 | -3.474 | 0.0051 |
| (ICI) - (No Tx) | -0.0272 | 0.0167 | 477 | -1.626 | 0.4813 |

**Table S30.** Linear mixed model coefficient estimates for fixed effects in **Figure 3G** (Intermediate ICI)

| Log_10_(vol+0.0001) ~ treat + day + treat*day + (1 \| id) | | | |
| --- | --- | --- | --- |
|  | **Estimate** | **95% CI** | **P-value** |
| Day | -0.0087 | (-0.0180, 0.0006) | 0.0691 |
| Intermediate 177Lu+ICI | 0.1437 | (-0.3172, 0.6086) | 0.5456 |
| Intermediate 225Ac+ICI | 0.0485 | (-0.3924, 0.4940) | 0.8311 |
| ICI | 0.0967 | (-0.4556, 0.6573) | 0.7348 |
| No Tx | 0.1469 | (-0.4276, 0.7271) | 0.6205 |
| Day : Intermediate 177Lu+ICI | 0.0226 | (0.0096, 0.0356) | 0.0008 |
| Day : Intermediate 225Ac+ICI | 0.0201 | (0.0079, 0.0324) | 0.0014 |
| Day : ICI | 0.0276 | (0.0104, 0.0447) | 0.0018 |
| Day : No Tx | 0.0402 | (0.0220, 0.0585) | <.0001 |

**Table S31.** Tukey-adjusted pairwise comparisons of estimated marginal means in **Figure 3G** (Intermediate ICI)

| Log_10_(vol+0.0001) ~ treat + day + treat*day + (1 \| id) | | | | | |
| --- | --- | --- | --- | --- | --- |
| **contrast** | **estimate** | **SE** | **df** | **t.ratio** | **p.value** |
| (Intermediate 90Y+ICI) - (Intermediate 177Lu+ICI) | -0.0226 | 0.0067 | 592 | -3.387 | 0.0067 |
| (Intermediate 90Y+ICI) - (Intermediate 225Ac+ICI) | -0.0201 | 0.0063 | 592 | -3.201 | 0.0125 |
| (Intermediate 90Y+ICI) - (ICI) | -0.0276 | 0.0088 | 595 | -3.134 | 0.0155 |
| (Intermediate 90Y+ICI) - (No Tx) | -0.0402 | 0.0094 | 594 | -4.289 | 0.0002 |
| (Intermediate 177Lu+ICI) - (Intermediate 225Ac+ICI) | 0.0025 | 0.0062 | 592 | 0.400 | 0.9946 |
| (Intermediate 177Lu+ICI) - (ICI) | -0.0050 | 0.0088 | 595 | -0.572 | 0.9791 |
| (Intermediate 177Lu+ICI) - (No Tx) | -0.0176 | 0.0093 | 594 | -1.891 | 0.3230 |
| (Intermediate 225Ac+ICI) - (ICI) | -0.0075 | 0.0085 | 595 | -0.885 | 0.9024 |
| (Intermediate 225Ac+ICI) - (No Tx) | -0.0201 | 0.0090 | 594 | -2.222 | 0.1728 |
| (ICI) - (No Tx) | -0.0126 | 0.0110 | 596 | -1.145 | 0.7826 |

**Table S32.** Linear mixed model coefficient estimates for fixed effects in **Figure 3G** (Delayed ICI)

| Log_10_(vol) ~ treat + day + treat*day + (1 \| id) | | | |
| --- | --- | --- | --- |
|  | **Estimate** | **95% CI** | **P-value** |
| Day | 0.0192 | (0.0153, 0.0231) | <.0001 |
| Delayed 177Lu+ICI | 0.0111 | (-0.1800, 0.2034) | 0.9100 |
| Delayed 225Ac+ICI | -0.0042 | (-0.1953, 0.1866) | 0.9655 |
| ICI | -0.0373 | (-0.2722, 0.2097) | 0.7596 |
| No Tx | -0.0130 | (-0.2509, 0.2260) | 0.9158 |
| Day : Delayed 177Lu+ICI | -0.0049 | (-0.0105, 0.0006) | 0.0861 |
| Day : Delayed 225Ac+ICI | -0.0084 | (-0.0138, -0.0031) | 0.0023 |
| Day : ICI | -0.0024 | (-0.0095, 0.0046) | 0.5038 |
| Day : No Tx | 0.0131 | (0.0056, 0.0208) | 0.0008 |

**Table S33.** Tukey-adjusted pairwise comparisons of estimated marginal means in **Figure 3G** (Delayed ICI)

| Log_10_(vol) ~ treat + day + treat*day + (1 \| id) | | | | | |
| --- | --- | --- | --- | --- | --- |
| **contrast** | **estimate** | **SE** | **df** | **t.ratio** | **p.value** |
| (Delayed 90Y+ICI) - (Delayed 177Lu+ICI) | 0.0049 | 0.0029 | 556 | 1.719 | 0.4232 |
| (Delayed 90Y+ICI) - (Delayed 225Ac+ICI) | 0.0084 | 0.0028 | 556 | 3.067 | 0.0192 |
| (Delayed 90Y+ICI) - (ICI) | 0.0024 | 0.0036 | 558 | 0.668 | 0.9631 |
| (Delayed 90Y+ICI) - (No Tx) | -0.0132 | 0.0039 | 557 | -3.365 | 0.0073 |
| (Delayed 177Lu+ICI) - (Delayed 225Ac+ICI) | 0.0035 | 0.0028 | 556 | 1.278 | 0.7050 |
| (Delayed 177Lu+ICI) - (ICI) | -0.0025 | 0.0036 | 557 | -0.689 | 0.9589 |
| (Delayed 177Lu+ICI) - (No Tx) | -0.0181 | 0.0039 | 558 | -4.597 | 0.0001 |
| (Delayed 225Ac+ICI) - (ICI) | -0.0060 | 0.0035 | 558 | -1.700 | 0.4346 |
| (Delayed 225Ac+ICI) - (No Tx) | -0.0216 | 0.0039 | 557 | -5.611 | <.0001 |
| (ICI) - (No Tx) | -0.0156 | 0.0045 | 559 | -3.442 | 0.0056 |

**Table S34.** Log-rank test of B78 overall survival (Varied Radionuclide) in **Figure 3I**

| **contrast** | **p.value** |
| --- | --- |
| (Early 90Y+ICI) - (Early Lu+ICI) | 0.8875 |
| (Early 90Y+ICI) - (Early 225Ac+ICI) | 0.7042 |
| (Early Lu+ICI) – (Early 225Ac+ICI) | 0.6773 |
| (Intermediate 90Y+ICI) - (Intermediate Lu+ICI) | 0.9916 |
| (Intermediate 90Y+ICI) - (Intermediate 225Ac+ICI) | 0.5537 |
| (Intermediate Lu+ICI) – (Intermediate 225Ac+ICI) | 0.4707 |
| (Delayed 90Y+ICI) - (Delayed Lu+ICI) | 0.3719 |
| (Delayed 90Y+ICI) - (Delayed 225Ac+ICI) | 0.0124 * |
| (Delayed Lu+ICI) – (Delayed 225Ac+ICI) | 0.1753 |

**Table S35.** Linear mixed model coefficient estimates for fixed effects in **Figure 4B**

| Log_10_(vol+0.0001) ~ treat + day + treat*day + (1 \| id) | | | |
| --- | --- | --- | --- |
|  | **Estimate** | **95% CI** | **P-value** |
| Day | -0.1565 | (-0.1791, -0.1339) | <.0001 |
| 90Y | 1.2645 | (0.7198, 1.8176) | <.0001 |
| 90Y+ICI+aCD8 | 1.5201 | (0.9245, 2.1203) | <.0001 |
| aCD8 | 1.5557 | (0.9525, 2.1644) | <.0001 |
| Day : 90Y | 0.2066 | (0.1716, 0.2421) | <.0001 |
| Day : 90Y+ICI+aCD8 | 0.1983 | (0.1602, 0.2367) | <.0001 |
| Day : aCD8 | 0.2401 | (0.1898, 0.2906) | <.0001 |

**Table S36.** Tukey-adjusted pairwise comparisons of estimated marginal means in **Figure 4B**

| Log_10_(vol+0.0001) ~ treat + day + treat*day + (1 \| id) | | | | | |
| --- | --- | --- | --- | --- | --- |
| **contrast** | **estimate** | **SE** | **df** | **t.ratio** | **p.value** |
| (90Y+ICI) - (90Y) | -0.2066 | 0.0182 | 240 | -11.364 | <.0001 |
| (90Y+ICI) - (90Y+ICI+aCD8) | -0.1983 | 0.0197 | 240 | -10.049 | <.0001 |
| (90Y+ICI) - (aCD8) | -0.2401 | 0.0260 | 240 | -9.231 | <.0001 |
| (90Y) - (90Y+ICI+aCD8) | 0.0083 | 0.0211 | 239 | 0.393 | 0.9794 |
| (90Y) - (aCD8) | -0.0335 | 0.0271 | 240 | -1.235 | 0.6053 |
| (90Y+ICI+aCD8) - (aCD8) | -0.0418 | 0.0282 | 240 | -1.483 | 0.4496 |

**Table S37.** Linear mixed model coefficient estimates for fixed effects in **Figure 4C**

| Log_10_(vol+0.0001) ~ treat + day + treat*day + (1 \| id) | | | |
| --- | --- | --- | --- |
|  | **Estimate** | **95% CI** | **P-value** |
| Day | -0.0974 | (-0.1396, -0.0556) | <.0001 |
| 177Lu | 0.1886 | (-0.8910, 1.2709) | 0.7369 |
| 177Lu+ICI+aCD8 | 0.8701 | (-0.3777, 2.1148) | 0.1802 |
| aCD8 | 1.0839 | (-0.2136, 2.3748) | 0.1083 |
| Day : 177Lu | 0.1909 | (0.1140, 0.2673) | <.0001 |
| Day : 177Lu+ICI+aCD8 | 0.1425 | (0.0460, 0.2398) | 0.0052 |
| Day : aCD8 | 0.1275 | (0.0089, 0.2476) | 0.0409 |

**Table S38.** Tukey-adjusted pairwise comparisons of estimated marginal means in **Figure 4C**

| Log_10_(vol+0.0001) ~ treat + day + treat*day + (1 \| id) | | | | | |
| --- | --- | --- | --- | --- | --- |
| **contrast** | **estimate** | **SE** | **df** | **t.ratio** | **p.value** |
| (177Lu+ICI) - (177Lu) | -0.1909 | 0.0398 | 154 | -4.798 | <.0001 |
| (177Lu+ICI) - (177Lu+ICI+aCD8) | -0.1425 | 0.0503 | 153 | -2.835 | 0.0264 |
| (177Lu+ICI) - (aCD8) | -0.1275 | 0.0619 | 154 | -2.059 | 0.1712 |
| (177Lu) - (177Lu+ICI+aCD8) | 0.0484 | 0.0569 | 154 | 0.852 | 0.8296 |
| (177Lu) - (aCD8) | 0.0635 | 0.0668 | 154 | 0.950 | 0.7778 |
| (177Lu+ICI+aCD8) - (aCD8) | 0.0150 | 0.0736 | 154 | 0.204 | 0.9970 |

**Table S39.** Linear mixed model coefficient estimates for fixed effects in **Figure 4D**

| Log_10_(vol+0.0001) ~ treat + day + treat*day + (1 \| id) | | | |
| --- | --- | --- | --- |
|  | **Estimate** | **95% CI** | **P-value** |
| Day | 0.0054 | (-0.0159, 0.0266) | 0.6229 |
| 225Ac | 0.7256 | (0.1508, 1.3069) | 0.0163 |
| 225Ac+ICI+aCD8 | 0.9033 | (0.2749, 1.5400) | 0.0064 |
| aCD8 | 0.8780 | (0.1453, 1.6135) | 0.0221 |
| Day : 225Ac | 0.0336 | (-0.0032, 0.0706) | 0.0801 |
| Day : 225Ac+ICI+aCD8 | 0.0259 | (-0.0148, 0.0663) | 0.2182 |
| Day : aCD8 | 0.0544 | (-0.0225, 0.1320) | 0.1754 |

**Table S40.** Tukey-adjusted pairwise comparisons of estimated marginal means in **Figure 4D**

| Log_10_(vol+0.0001) ~ treat + day + treat*day + (1 \| id) | | | | | |
| --- | --- | --- | --- | --- | --- |
| **contrast** | **estimate** | **SE** | **df** | **t.ratio** | **p.value** |
| (225Ac+ICI) - (225Ac) | -0.0336 | 0.0191 | 171 | -1.757 | 0.2977 |
| (225Ac+ICI) - (225Ac+ICI+aCD8) | -0.0259 | 0.0210 | 171 | -1.234 | 0.6061 |
| (225Ac+ICI) - (aCD8) | -0.0544 | 0.0400 | 170 | -1.360 | 0.5260 |
| (225Ac) - (225Ac+ICI+aCD8) | 0.0077 | 0.0240 | 172 | 0.322 | 0.9885 |
| (225Ac) - (aCD8) | -0.0208 | 0.0416 | 171 | -0.500 | 0.9590 |
| (225Ac+ICI+aCD8) - (aCD8) | -0.0285 | 0.0424 | 170 | -0.671 | 0.9077 |

**Table S41.** Log-rank test of MC38 overall survival (CD8 depletion) in **Figure 4I-K**

| **contrast** | **p.value** |
| --- | --- |
| (90Y+ICI) - (90Y) | <0.0001 **** |
| (90Y+ICI) - (90Y+ICI+aCD8) | 0.0044 ** |
| (90Y+ICI) – (aCD8) | <0.0001 **** |
| (90Y) – (90Y+ICI+aCD8) | 0.7725 |
| (90Y) – (aCD8) | <0.0001 **** |
| (90Y+ICI+aCD8) – (aCD8) | 0.0022 ** |
| (Lu+ICI) - (Lu) | <0.0001 **** |
| (Lu+ICI) - (Lu+ICI+aCD8) | 0.0003 *** |
| (Lu+ICI) – (aCD8) | <0.0001 **** |
| (Lu) – (Lu+ICI+aCD8) | 0.2594 |
| (Lu) – (aCD8) | 0.0012 ** |
| (Lu+ICI+aCD8) – (aCD8) | 0.1591 |
| (225Ac+ICI) - (225Ac) | <0.0001 **** |
| (225Ac+ICI) - (225Ac+ICI+aCD8) | <0.0001 **** |
| (225Ac+ICI) – (aCD8) | <0.0001 **** |
| (225Ac) – (225Ac+ICI+aCD8) | 0.3030 |
| (225Ac) – (aCD8) | <0.0001 **** |
| (225Ac+ICI+aCD8) – (aCD8) | 0.0028 ** |

**Table S42.** Linear mixed model coefficient estimates for fixed effects in **Figure 5E**

| Log_10_(vol+0.0001) ~ treat + day + treat*day + (1 \| id) | | | |
| --- | --- | --- | --- |
|  | **Estimate** | **95% CI** | **P-value** |
| Day | -0.1771 | (-0.2331, -0.1211) | <.0001 |
| 225Ac | 0.9587 | (-0.4937, 2.4112) | 0.2047 |
| ICI | 0.4389 | (-1.0135, 1.8914) | 0.5607 |
| No Tx | 1.2959 | (-0.2138, 2.7871) | 0.0974 |
| Day : 225Ac | 0.1957 | (0.1166, 0.2749) | <.0001 |
| Day: ICI | 0.0562 | (-0.0230, 0.1353) | 0.1731 |
| Day: No Tx | 0.2028 | (0.1134, 0.2945) | <.0001 |

**Table S43.** Tukey-adjusted pairwise comparisons of estimated marginal means in **Figure 5E**

| Log_10_(vol+0.0001) ~ treat + day + treat*day + (1 \| id) | | | | | |
| --- | --- | --- | --- | --- | --- |
| **contrast** | **estimate** | **SE** | **df** | **t.ratio** | **p.value** |
| (225Ac+ICI) - (225Ac) | -0.1957 | 0.0410 | 158 | -4.769 | <.0001 |
| (225Ac+ICI) - (ICI) | -0.0562 | 0.0410 | 158 | -1.368 | 0.5211 |
| (225Ac+ICI) - (No Tx) | -0.2028 | 0.0469 | 159 | -4.327 | 0.0002 |
| (225Ac) - (ICI) | 0.1396 | 0.0410 | 158 | 3.401 | 0.0047 |
| (225Ac) - (No Tx) | -0.0071 | 0.0469 | 159 | -0.150 | 0.9988 |
| (ICI) - (No Tx) | -0.1466 | 0.0469 | 159 | -3.129 | 0.0111 |

**Table S44.** Log-rank test of MC38 overall survival in **Figure 5F**

| **contrast** | **p.value** |
| --- | --- |
| (225Ac+ICI) - (225Ac) | 0.0027 ** |
| (225Ac+ICI) - (ICI) | 0.1343 |
| (225Ac+ICI) – (No Tx) | 0.0018 ** |
| (225Ac) – (ICI) | 0.0263 * |
| (225Ac) – (No Tx) | 0.0830 |
| (ICI) – (No Tx) | 0.0064 ** |

**Table S45.** List of flow cytometry antibody targets, clones, and fluorophores

| **Name** | **Clone** | **Fluorophore** | **Catalog number** |
| --- | --- | --- | --- |
| CD4 | RM4-5 | FITC | BioLegend 100510 |
| NK1.1 | PK136 | PE-Cy5 | BioLegend 108716 |
| FOXP3 | FJK-16s | PE-Cy7 | Thermo Scientific 25-5773-82 |
| CD45 | 30-F11 | BV605 | BioLegend 103140 |
| CD11b | M1/70 | BV711 | BioLegend 101242 |
| CD25 | PC61 | BV510 | BioLegend 102041 |
| CD8a | 53-6.7 | PE-Dazzle594 | BioLegend 100761 |
| CD69 | H1.2F3 | Alexa700 | BioLegend 104539 |
| IFNγ | XMG1.2 | BV421 | BioLegend 505830 |
| TNFα | MP6-XT22 | PE | BioLegend 506306 |
| CD44 | IM7 | BV510 | BioLegend 103043 |
| CD62L | MEL-14 | BV711 | BioLegend 104445 |
| CD3 | 17A2 | FITC | BioLegend 100204 |
| CD8a | 53-6.7 | PerCP-Cy5.5 | BioLegend 100733 |
| CD4 | RM4-5 | PE-Dazzle594 | BioLegend 100565 |
| CD45 | 30-F11 | PE-Cy7 | BioLegend 103113 |
| IFNγ | XMG1.2 | APC | BioLegend 505809 |
| TNFα | MP6-XT22 | PE-Dazzle594 | BioLegend 506346 |
| CD62L | MEL-14 | APC | BioLegend 104412 |

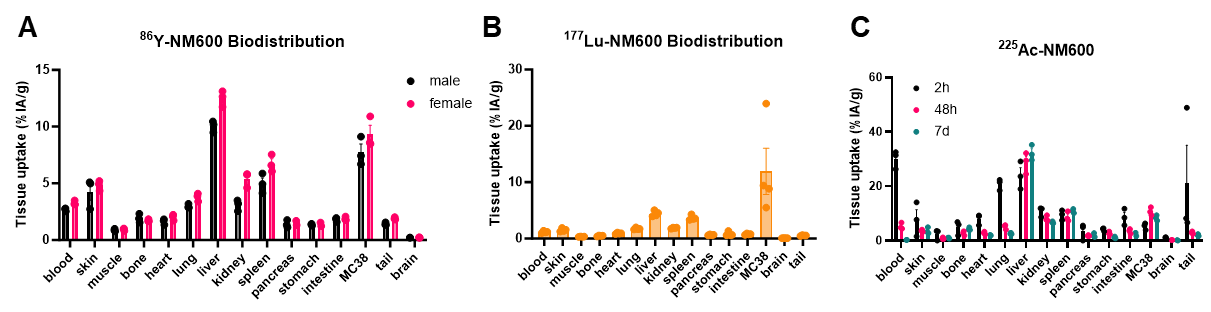
**Figure S1.** ^86^Y-, ^177^Lu-, ^225^Ac-NM600 biodistribution in MC38 tumor-bearing mice. A) *Ex vivo* biodistribution 72h post injection of 9.25 MBq ^86^Y-NM600 in male or female MC38 tumor-bearing mice (n=3/sex). B) *Ex vivo* biodistribution 180h post injection of 18.5 MBq ^177^Lu-NM600 in MC38 tumor-bearing mice (n=4). C) Longitudinal *ex vivo* biodistribution of ^225^Ac-NM600 2h, 48h, or 7d following tail vein injection of 9.25 kBq ^225^Ac-NM600 in MC38 tumor-bearing mice (n=3/timepoint).

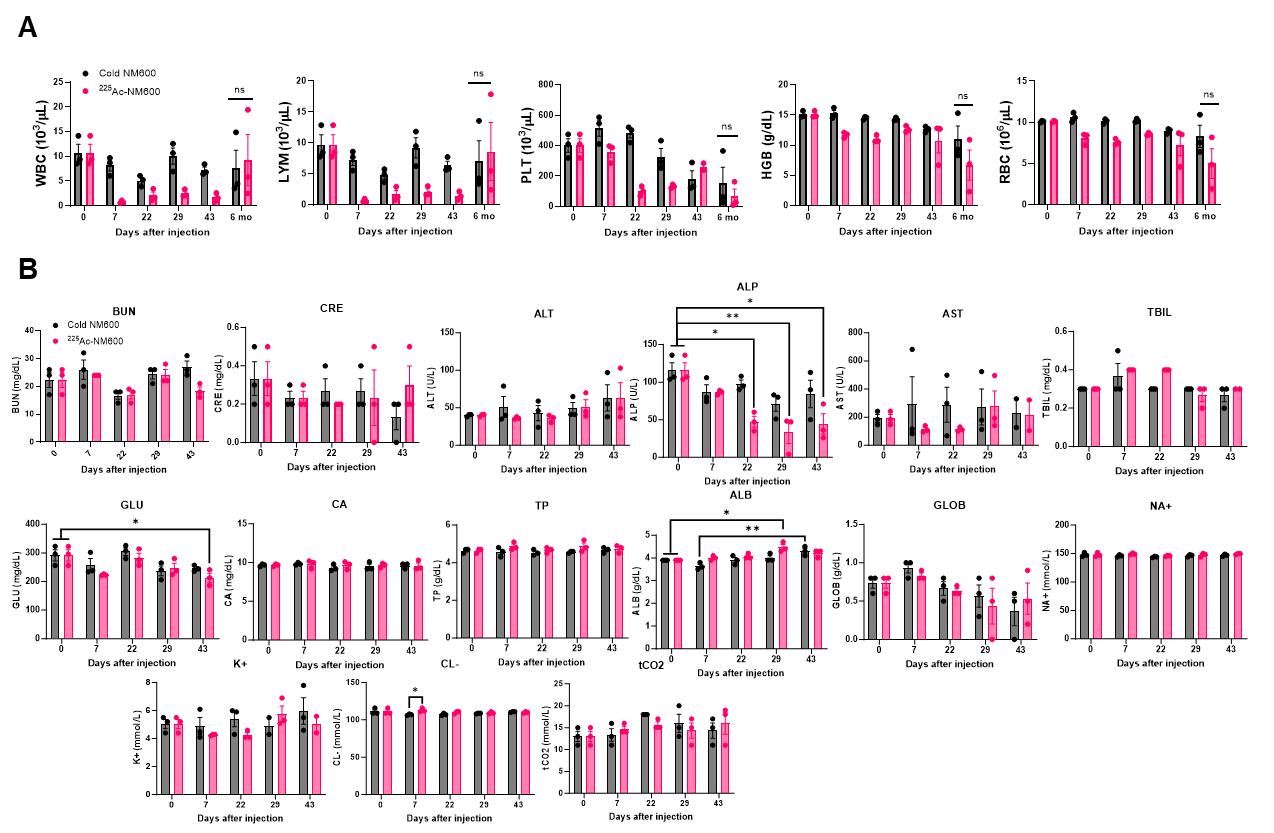
**Figure S2.** Acute toxicity profile of ^225^Ac-NM600. Naïve C57BL/6 mice received either 18.5 kBq ^225^Ac-NM600 or equivalent mass unlabeled NM600 (50 ng cold NM600). A) Serial complete blood counts (CBCs), n=3/timepoint. WBC: white blood cells, LYM: lymphocytes, PLT: platelets, HGB: hemoglobin, RBC: red blood cells. B) Serial comprehensive metabolic panels (CMPs), n=3/timepoint. BUN: blood urea nitrogen, CRE: creatinine, ALT: alanine aminotransferase, ALP: alkaline phosphatase, AST: aspartate aminotransferase, TBIL: total bilirubin, GLU: glucose, CA: calcium, TP: total protein, ALB: albumin, GLOB: hemoglobin, NA+: sodium, K+: potassium, CL-: chloride, tCO2: total carbon dioxide. A) Unpaired two-tailed t-test was used to compare 6 month mouse CBCs between treatment groups. B) Two-way ANOVA with Tukey’s HSD post hoc test was used to compare CMPs between treatment days and treatment groups.

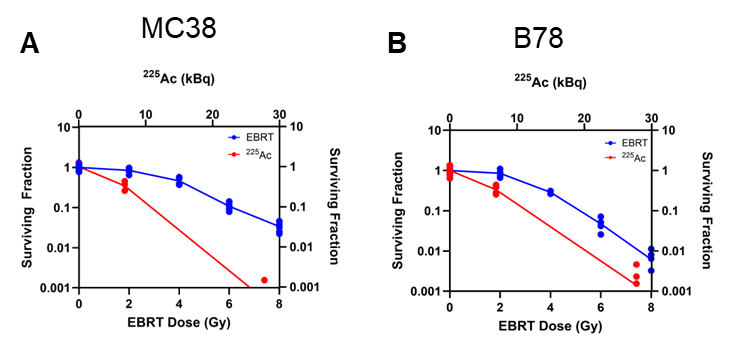
**Figure S3.** Clonogenic assays of MC38 and B78 following EBRT and ^225^Ac. Pre-plated tumor cells were irradiated with indicated doses (0, 2, 4, 6, or 8 Gy external beam radiation therapy or 0, 0.25, or 1 Gy ^225^Ac delivered in 24 hours), then grown for five days. The clonogenic colonies were quantified and are displayed as a percent of total plating efficiency. N=6/treatment group; n=12: 0 Gy control.

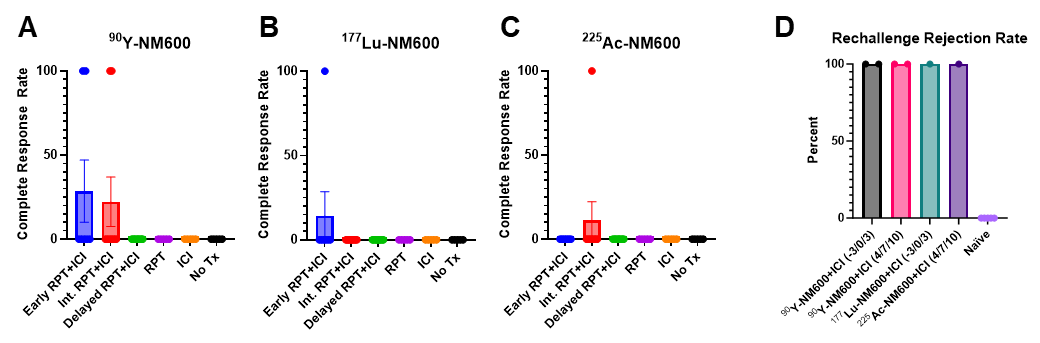
**Figure S4.** Complete response and rechallenge rejection rates for B78 melanoma. A-C) Complete response rates for all B78 treatment groups. Data points at 100 indicate a mouse with no palpable tumor on day 100; 0 indicate mouse deaths or palpable tumors on day 100. D) All complete responder mice and naïve controls were rechallenged with 2*10^6^ B78 cells on the left flank on day 120. Data points at 100 indicate no palpable tumor; 0 indicate palpable tumors four weeks following rechallenge. A-C) N=9: ^90^Y-, ^177^Lu-, ^225^Ac-NM600 + ICI 4/7/10 or + ICI 11/14/17; n=7: ^90^Y-, ^177^Lu-, ^225^Ac-NM600 + ICI -3/0/3; n=5: ^90^Y-, ^177^Lu-, ^225^Ac-NM600, ICI 4/7/10, No Tx. D) N=2: ^90^Y-NM600 + ICI -3/0/3 or + ICI 4/7/10; n=1: ^177^Lu-NM600+ICI -3/0/3, ^225^Ac-NM600+ICI 4/7/10; n=5: naïve.

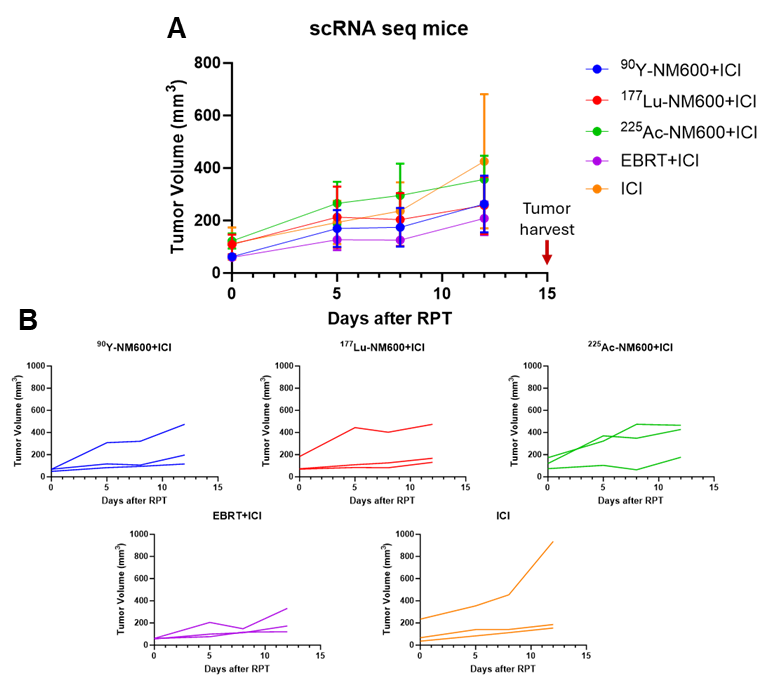
**Figure S5.** Tumor growth curves of single cell RNA sequencing experiment mice. A) Mean+SEM tumor volumes prior to euthanasia and tumor harvest on treatment day 15. B) Individual tumor growth curves. N=3/treatment group.

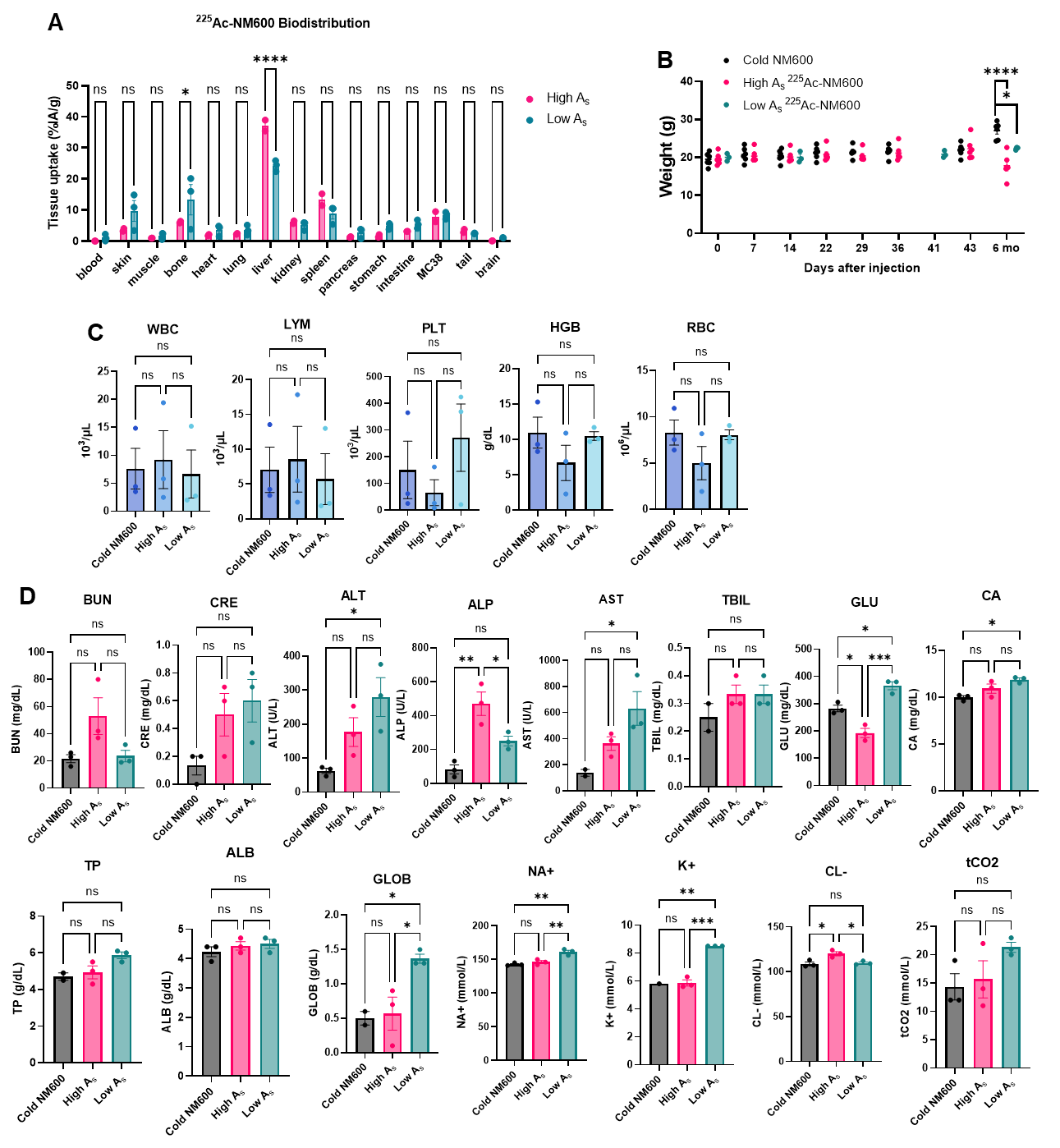
**Figure S6.** Low vs. high specific activity ^225^Ac-NM600 biodistribution and late (six month) toxicity profile A) *Ex vivo* biodistribution of high specific activity (high A_s_; 2.70 μg NM600/MBq ^225^Ac; n=2) or low A_s_ (81.1 μg NM600/MBq ^225^Ac; n=3) 11 days following injection of 9.25 kBq ^225^Ac-NM600 in MC38 tumor-bearing mice. B-D) Naïve C57BL/6 mice received either cold NM600 (50 ng NM600/mouse; equivalent mass NM600 of high A_s_ ^225^Ac-NM600), 18.5 kBq high A_s_ or low A_s_ ^225^Ac-NM600. B) Overall mouse body weights over time. Cold NM600/high A_s_ ^225^Ac-NM600: n=6; low A_s_ ^225^Ac-NM600: n=3. C) Complete blood counts (CBCs) six months post ^225^Ac-NM600 administration. N=3/treatment group. WBC: white blood cells, LYM: lymphocytes, PLT: platelets, HGB: hemoglobin, RBC: red blood cells. D) Comprehensive metabolic panel (CMP) six months post ^225^Ac-NM600 administration. N=3/treatment group. BUN: blood urea nitrogen, CRE: creatinine, ALT: alanine aminotransferase, ALP: alkaline phosphatase, AST: aspartate aminotransferase, TBIL: total bilirubin, GLU: glucose, CA: calcium, TP: total protein, ALB: albumin, GLOB: hemoglobin, NA+: sodium, K+: potassium, CL-: chloride, tCO2: total carbon dioxide. A) Unpaired two-tailed t-test was used to compare tissue biodistribution between treatment groups. B-D) One-way ANOVA with Tukey’s HSD post hoc test was used to compare between treatment groups.
